# βIII-Tubulin is a Brake on Extrinsic Cell-Death in Pancreatic Cancer

**DOI:** 10.1101/2022.09.29.510034

**Authors:** John Kokkinos, George Sharbeen, Rosa Mistica C. Ignacio, Elvis Pandzic, Janet Youkhana, Cyrille Boyer, Koroush S. Haghighi, Matthew Gunawarman, David Goldstein, Val Gebski, Marina Pajic, Omali Pitiyarachchi, Meagan E. Davis, Grace Schulstad, Oliver S.M. Arkell, Chantal Kopecky, Estrella Gonzales-Aloy, Mert Erkan, Jennifer P. Morton, Maria Kavallaris, Peter W. Gunning, Edna C. Hardeman, Amber Johns, Anthony J. Gill, Renee M. Whan, Amanda Mawson, Australian Pancreatic Cancer Genome Initiative, Joshua A. McCarroll, Phoebe A. Phillips

## Abstract

The microtubule protein, βIII-tubulin, has been implicated as a prognostic, pro-survival, and chemoresistance factor in some of the most lethal malignancies including pancreatic ductal adenocarcinoma (PDAC). However, precise survival mechanisms controlled by βIII-tubulin in cancer cells are unknown. Here, we report an unexpected role of βIII-tubulin as a brake on extrinsic caspase 8-dependent apoptosis in PDAC. We show that βIII-tubulin knockdown frees death-receptor DR5 to increase its membrane diffusion, clustering, and activation of cell-death. We demonstrate that βIII-ubulin silencing increases sensitivity of PDAC cells to chemotherapeutic and microenvironment-derived extrinsic cell-death signals including TRAIL, TNFα, and FasL. Finally, nanoparticle delivery of βIII-tubulin siRNA to mouse orthotopic PDAC tumours *in vivo* and human patient-derived PDAC tumour explants *ex vivo* increases extrinsic apoptosis and reduces tumour progression. Thus, silencing of βIII-tubulin represents an innovative strategy to unleash a suicide signal in PDAC cells and render them sensitive to microenvironment and chemotherapy-derived death signals.

## Introduction

Pancreatic ductal adenocarcinoma (PDAC) is one of the deadliest malignancies with 5-year survival of <11%[1]. This is attributed to the disease being diagnosed at an advanced stage, with chemoresistance, metastases, and a complex multi-cellular microenvironment that can drive tumour progression[2-6]. In fact, the highly fibrotic stroma produced by cancer-associated fibroblasts (CAFs) can hinder drug delivery and together with pro-tumourigenic crosstalk between PDAC cells and CAFs promote the metastatic and chemoresistant nature of PDAC[7, 8]. Novel therapeutic strategies should not simply target PDAC cells only but address the complex and heterogeneous multicellular microenvironment that characterises PDAC[6].

The microtubule protein, βIII-tubulin, is upregulated in solid tumours and correlates with poor patient outcome[9-15]. In PDAC, βIII-tubulin is upregulated in patient tumour specimens[16-18]. Several studies in multiple tumour types have reported links between βIII-tubulin and pro-tumour pathways including the PTEN/AKT pathway[19], protection against nutrient deprivation and microenvironment-derived stress[20], and promotion of chemoresistance[16, 21-24] and metastasis[24, 25]. However, despite the wealth of evidence establishing βIII-tubulin as a broad promoter of tumour progression, the mechanism by which βIII-tubulin regulates these survival pathways and protects cancer cells from chemotherapy and microenvironment-derived stress remains unknown. This information is critical for application of βIII-tubulin inhibition in the clinic. Given we previously identified βIII-tubulin silencing in PDAC cells as an activator of apoptosis[16], we investigated apoptotic pathways controlled by βIII-tubulin in PDAC.

Here, we showed that βIII-tubulin regulates the extrinsic caspase 8-dependent apoptotic pathway in PDAC. We also identified that silencing βIII-tubulin increased sensitivity to microenvironment and chemotherapeutic-derived extrinsic apoptosis inducers, including TNF-related apoptosis-inducing ligand (TRAIL). We observed that βIII-tubulin forms a mesh-like structure at the membrane of PDAC cells and silencing βIII-tubulin increases membrane diffusion of the TRAIL-receptor DR5 to trigger its clustering and activation of cell-death.

We used polymeric nanoparticles (Star 3)[26] to investigate the therapeutic effects of βIII-tubulin silencing *in vivo* and in patient-derived PDAC explants *ex vivo*[27]. In orthotopic PDAC mouse tumours, Star 3-βIII-tubulin siRNA significantly reduced tumour growth. Additionally, in patient-derived PDAC explants, βIII-tubulin silencing decreased tumour cell frequency and increased sensitivity to TRAIL. Finally, we showed for the first time that high stromal expression of βIII-tubulin in PDAC patients is independently prognostic of poorer overall patient survival.

## Results

### Knockdown of βIII-tubulin in PDAC cells activated extrinsic apoptosis

We used a smartpool of siRNAs to inhibit βIII-tubulin expression in PDAC cells and confirmed knockdown by qPCR (**Supplementary Figure 1A-B**) and western blot (**Supplementary Figure 1C-H**). Knockdown of βIII-tubulin in MiaPaCa2 cells significantly increased caspase-9 (**Figure 1A**) and caspase-8 activities (**Figure 1B**), relative to controls. To delineate the functional relevance of each caspase, we blocked their individual activities with inhibitors in MiaPaCa2 and PANC1 cells. Apoptosis induced by βIII-tubulin knockdown in both PDAC cell lines, was unaffected by caspase-9 inhibition (intrinsic pathway; **Figure 1C-D**) but was completely blocked by caspase-8 inhibition (extrinsic pathway; **Figure 1E-F**).

**Figure 1.**
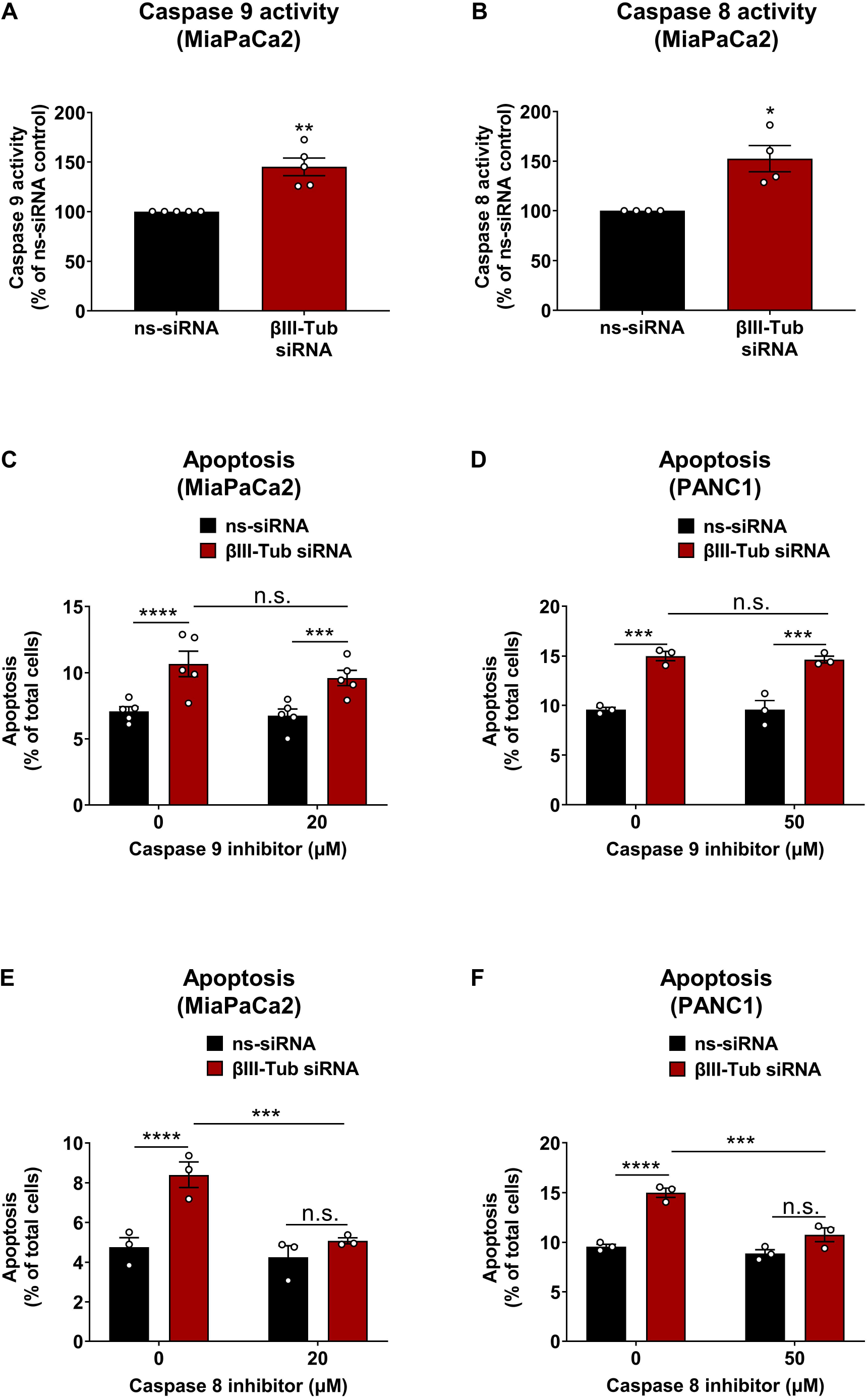
Knockdown of βIII-tubulin in PDAC cells activated the extrinsic pathway of apoptosis. **(A-B)** Caspase 9 (n=5) **(A)** and caspase 8 (n=4) **(B)** activities in MiaPaCa2 cells measured using CaspaseGlo assays, 72 hours post-transfection with non-silencing (ns) or βIII-tubulin (βIII-Tub) siRNA. Activity normalised to cell numbers. **(C-F)** Apoptosis measured using flow cytometry for Annexin V/DAPI in PDAC cells, 72 hours post-transfection with βIII-tubulin siRNA, and 24 hours post-treatment with caspase 9 inhibitor (Z-LEHD-FMK) **(C-D)** or caspase 8 inhibitor (Q-IETD-OPh) **(E-F)**. Results from panels D and F were obtained using the same controls for 0 μM caspase inhibitor. Bars represent mean of n≥3 independent experiments (data points shown from independent experiments) ± SEM. Asterisks indicate significance as assessed by two-tailed paired t-tests or one-way ANOVA, Bonferroni’s multiple comparisons test (*p≤0.05, **p≤0.01, ***p≤0.001, ****p≤0.0001, n.s.; non-significant).

### Knockdown of βIII-tubulin in mouse orthotopic pancreatic tumours decreased tumour progression and increased extrinsic apoptosis

To test if effects we observed *in vitro* would carry into an *in vivo* setting, we utilised an orthotopic PDAC tumour mouse model. Once established (4 weeks post-implant), tumours were randomised based on tumour luminescence (**Figure 2A-B**) then treated with control-siRNA or βIII-tubulin siRNA complexed to Star 3 nanoparticles (**Figure 2A**). βIII-tubulin knockdown in tumours was confirmed at endpoint (**Figure 2C-D**). Star 3+βIII-tubulin siRNA treatment significantly decreased tumour volume (**Figure 2E**) and increased intratumoural cleaved caspase 8-positive cells compared to controls (**Figure 2F**).

**Figure 2.**
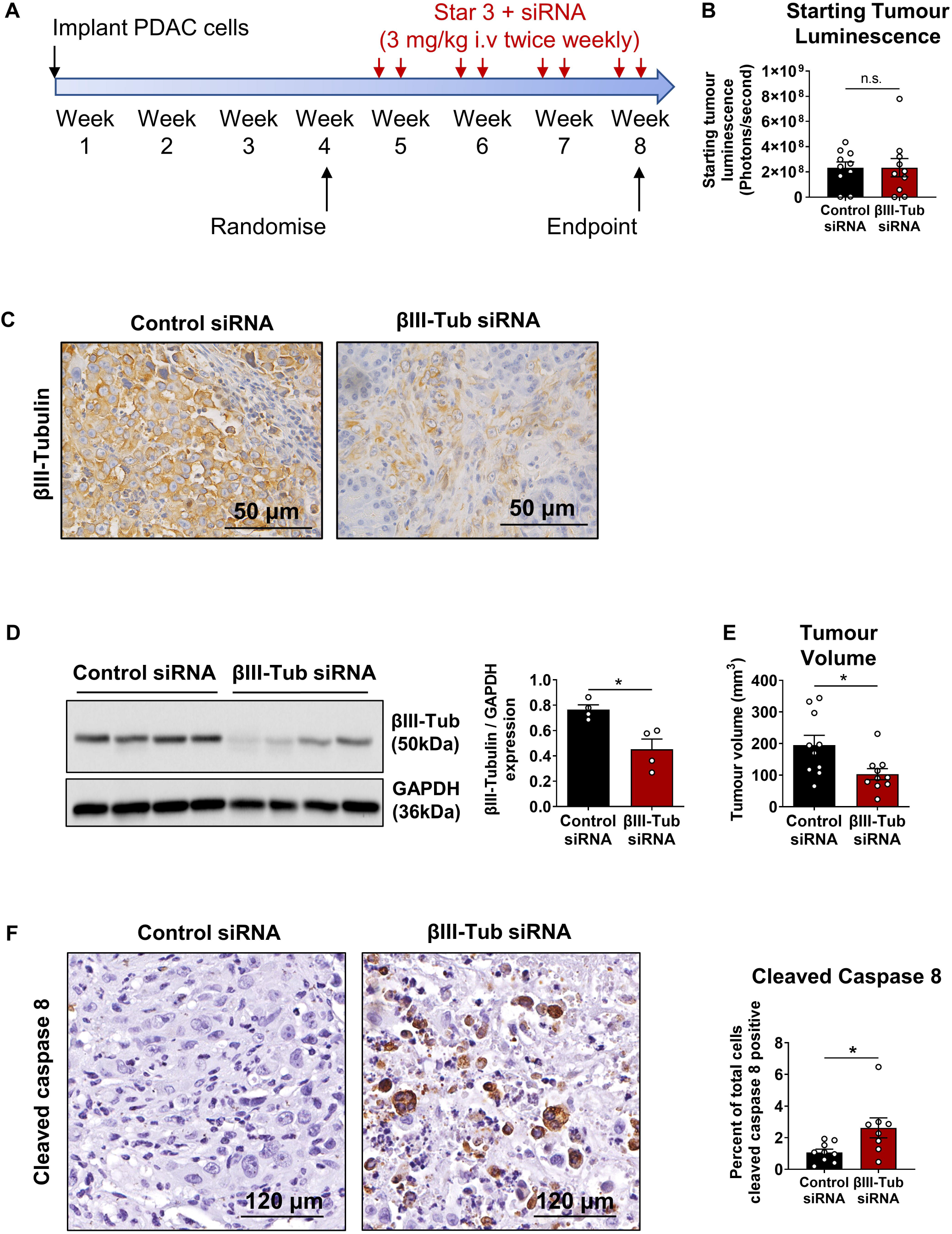
Star 3+βIII-tubulin siRNA decreased tumour growth and increased extrinsic apoptosis in mouse orthotopic PDAC tumours. **(A)** Treatment regimen used in mouse model. **(B)** Mice were randomised based on tumour luminescence. **(C)** Representative immunohistochemistry for βIII-tubulin (βIII-Tub) in sections from tumours treated with Star 3+control siRNA or βIII-tubulin siRNA. **(D)** Western blot of protein extracts from 8 mouse tumours treated with Star 3+control siRNA (n=4) or βIII-tubulin siRNA (n=4) and corresponding densitometry analysis. GAPDH was used as loading control. **(E)** Tumour volume at endpoint. **(F)** Cleaved caspase 8 immunohistochemistry representative images and quantification in Star 3+control siRNA (n=9) and Star 3+βIII-tubulin siRNA (n=8). One tumour was excluded from controls and 2 tumours excluded from βIII-tubulin siRNA group due to insufficient tumour tissue. Bars represent mean (data points indicate each mouse tumour) ± SEM. Asterisks indicate significance by two-tailed unpaired t-tests (*p≤0.05; n.s.; non-significant; n=10 mice per group).

### Knockdown of βIII-tubulin in PDAC cells increased sensitivity to TRAIL

We next investigated the effect of combining βIII-tubulin knockdown with TRAIL as an inducer of extrinsic apoptosis. All cell lines expressed both TRAIL receptors DR4 and DR5 (**Supplementary Figure 2A**) but expression level did not always correlate with relative TRAIL sensitivity of these cell lines in the experiments described below. βIII-tubulin knockdown using smartpool siRNA+TRAIL treatment in PDAC cells increased apoptosis (**Figure 3A-D**) and decreased viability (**Supplementary Figure 2B-D**) compared to either treatment alone. Likewise, live-cell analysis of MiaPaCa2 cells showed βIII-tubulin knockdown alone and TRAIL treatment alone reduced proliferation relative to controls, but combination of βIII-tubulin siRNA with TRAIL completely blocked proliferation (**Figure 3E**). MiaPaCa2 cells transfected with a single-sequence βIII-tubulin siRNA also significantly increased apoptosis in the presence of TRAIL (**Supplementary Figure 2E-F**).

**Figure 3.**
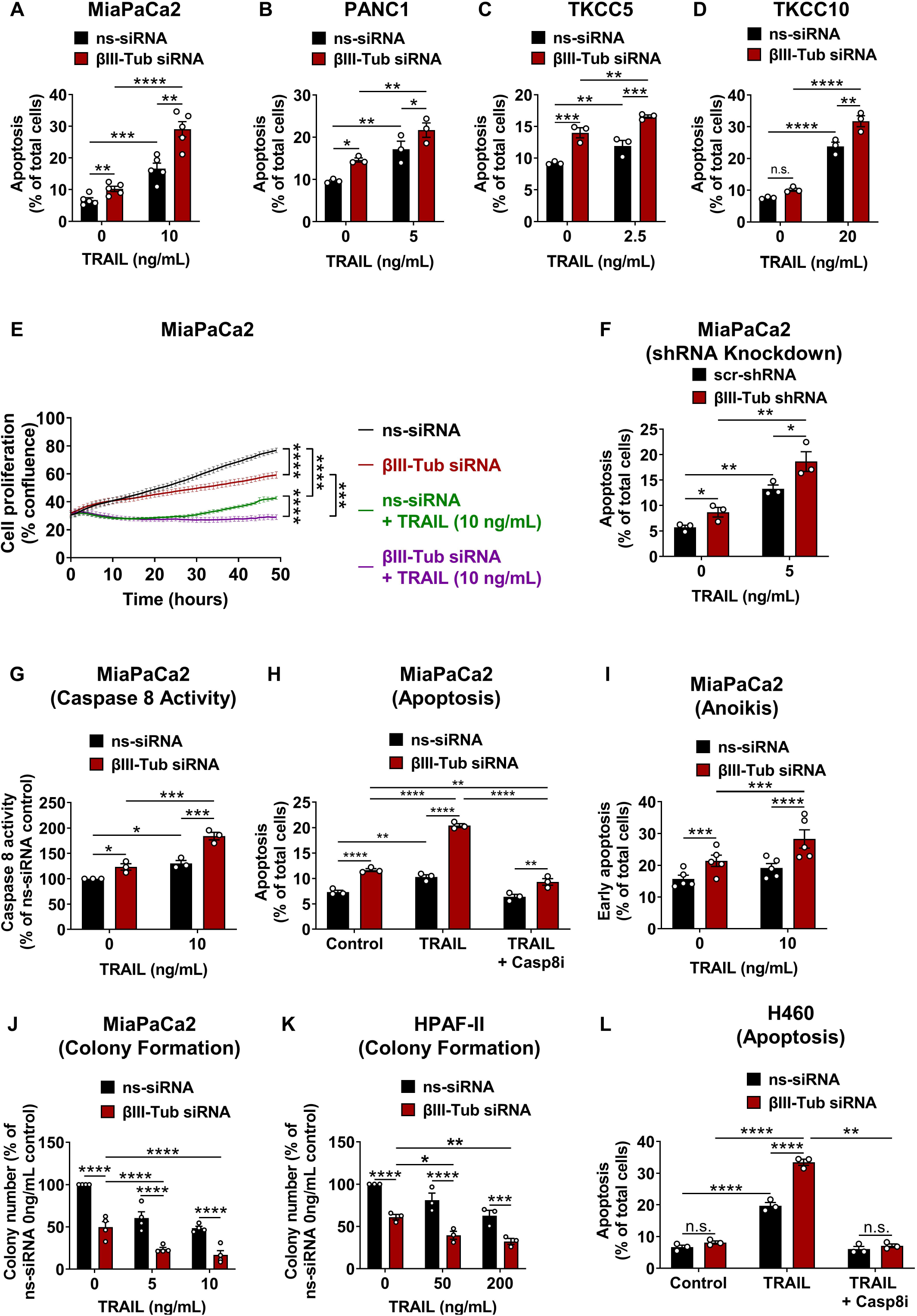
Knockdown of βIII-tubulin in PDAC cells sensitised them to TRAIL. **(A-D)** Apoptosis measured using flow cytometry for Annexin V/DAPI in PDAC cells transfected with non-silencing (ns) or βIII-tubulin (βIII-Tub) siRNA and treated ± TRAIL in MiaPaCa2 (n=5) **(A)**, PANC1 (n=3) **(B)**, TKCC5 (n=3) **(C)**, and TKCC10 (n=3) **(D)** cells. **(E)** Live cell proliferation analysis in MiaPaCa2 cells (n=4) with βIII-tubulin knockdown ± TRAIL. **(F)** Apoptosis measured in MiaPaCa2 cells (n=3) ±TRAIL with stable expression of βIII-tubulin or scramble (scr) shRNA. **(G)** Caspase 8 activity measured in MiaPaCa2 cells with βIII-tubulin knockdown ± TRAIL in MiaPaCa2 cells (n=3). Caspase 8 activity normalised to cell numbers. **(H)** Apoptosis was measured in MiaPaCa2 cells (n=3) with βIII-tubulin knockdown ± TRAIL (10 ng/mL) including a 1 hour pre-incubation with caspase 8 inhibitor (casp8i; 20 μM). **(I)** MiaPaCa2 cells (n=5) transfected with ns-siRNA or βIII-tubulin siRNA were cultured under anchorage independent conditions for 24 hours, then treated with TRAIL for 9 hours. Early apoptosis (Annexin V+/DAPI-) was measured using flow cytometry. **(J-K)** MiaPaCa2 (n=4) **(J)** and HPAF-II (n=3) **(K)** colonies formed from low seeding density, post-transfection with ns-siRNA or βIII-tubulin siRNA, with or without TRAIL (72 hours). **(L)** Apoptosis measured in non-small cell lung cancer (NSCLC) H460 cells (n=3) transfected with ns-siRNA or βIII-tubulin siRNA and treated ± TRAIL (2.5 ng/mL) and ± casp8i (50 μM). Bars represent mean of n≥3 independent experiments (data points shown from independent experiments) ± SEM. Asterisks indicate significance as assessed by one-way ANOVA, Bonferroni’s multiple comparisons test (*p ≤0.05, **p≤0.01, ***p≤0.001, ****p≤0.0001, n.s.; non-significant).

To confirm these effects, we also used MiaPaCa2 cells that stably express βIII-tubulin shRNA (**Supplementary Figure 2E**). Stable knockdown of βIII-tubulin increased apoptosis compared to scramble-shRNA controls, and apoptosis was further enhanced in the presence of TRAIL (**Figure 3F**). βIII-tubulin knockdown-mediated TRAIL sensitisation in PDAC cells was caspase-8 dependent, as demonstrated by significantly increased caspase-8 activity with combination treatment relative to either treatment alone (**Figure 3G**) and by the ability of caspase-8 inhibition to completely block apoptosis induced by the combination treatment (**Figure 3H**). Knockdown of βIII-tubulin+TRAIL also significantly increased sensitivity to anoikis (anchorage-independent growth) (**Figure 3I**) and reduced cell colony formation in PDAC cells compared to either treatment alone (**Figure 3J-3K, Supplementary Figure 2G-2F**).

We next examined if these effects were PDAC specific or extended to other cancer cell types which also express high levels of βIII-tubulin. We used H460 lung cancer cells and demonstrated comparable sensitisation to TRAIL-induced apoptosis (**Figure 3L**). As observed in MiaPaCa2 cells, this increase in apoptosis was blocked with caspase 8 inhibition (**Figure 3L**). Knockdown of βIII-tubulin+TRAIL in H460 cells also reduced cell proliferation to a greater extent than either treatment alone (**Supplementary Figure 2H**).

Finally, we used primary PDAC-derived cancer-associated fibroblasts (CAFs) to test the combination of βIII-tubulin knockdown with TRAIL in non-neoplastic cells. CAFs expressed βIII-tubulin to a similar degree as PDAC cells (**Supplementary Figure 3A**). In contrast to PDAC cells, silencing βIII-tubulin in CAFs in the absence of TRAIL had no effect on apoptosis (**Supplementary Figure 3B**). TRAIL treatment alone induced a downward trend in apoptosis in control transfected CAFs, but no difference in apoptosis between CAFs with βIII-tubulin knockdown alone, and those treated with TRAIL+βIII-tubulin knockdown (**Supplementary Figure 3B**). The statistically significant differences observed between ns-siRNA and βIII-tubulin-siRNA treated CAFs in the presence of TRAIL (**Supplementary Figure 3B**) were likely due to the downward trend in apoptosis induced by TRAIL alone, rather than sensitisation to TRAIL. Importantly, these changes in CAF apoptosis were minor compared to those observed in PDAC cells (see **Supplementary Figure 3B** vs **Figure 3A**). CAF cell proliferation was reduced with βIII-tubulin knockdown but showed no further reduction with TRAIL (**Supplementary Figure 3C**). CAFs from 5 PDAC patients had varying levels of TRAIL death receptor 4 (DR4), but levels were markedly lower than PDAC cells (**Supplementary Figure 3D**), while TRAIL death receptor 5 (DR5) expression was either comparable or increased in CAFs compared to PDAC cells (**Supplementary Figure 3D**). There was no link between expression of TRAIL receptors and TRAIL sensitivity.

### Silencing βIII-tubulin in PDAC cells increased sensitivity to extrinsic apoptosis inducers

We next investigated if βIII-tubulin silencing is a broad sensitiser to extrinsic apoptosis inducers including TNFα and FasL. βIII-tubulin knockdown combined with TNFα in PDAC cells increased apoptosis more than either treatment alone (**Figure 4A-D**). Combination of βIII-tubulin knockdown with TNFα in MiaPaCa2 cells also blocked proliferation, compared to single treatments and control cells (**Figure 4E**). βIII-tubulin knockdown in MiaPaCa2 (**Figure 4F**) and TKCC10 cells (**Figure 4G**) similarly enhanced FasL-induced apoptosis, relative to single treatments and control cells.

**Figure 4:**
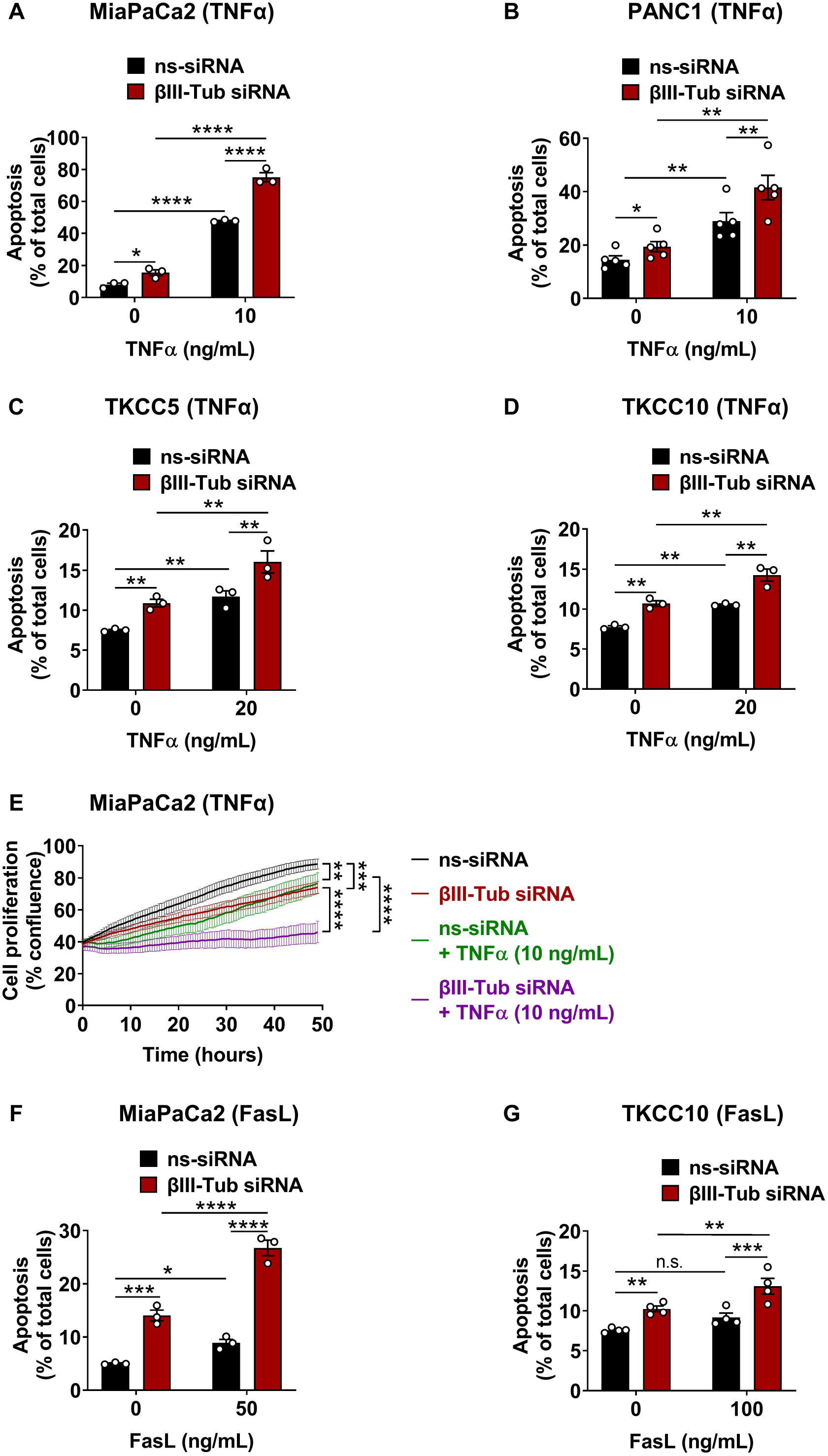
Knockdown of βIII-tubulin in PDAC cells sensitised them to TNFα and FasL-induced apoptosis. **(A-D)** Apoptosis measured using flow cytometry for Annexin V/DAPI in PDAC cells transfected with non-silencing (ns) or βIII-tubulin (βIII-Tub) siRNA and treated ± TNFα in MiaPaCa2 (n=3) **(A)**, PANC1 (n=5) **(B)**, TKCC5 (n=3) **(C)** and TKCC10 (n=3) **(D)** cells. **(E)** Cell proliferation analysis in MiaPaCa2 cells (n=4) with βIII-tubulin knockdown ±TNFα, measured on IncuCyte® S3 Live-Cell Analysis. **(F-G)** Apoptosis measured in MiaPaCa2 (n=3) **(F)** and TKCC10 (n=4) **(G)** cells transfected with ns-siRNA or βIII-tubulin siRNA and treated ± FasL. Bars represent mean of n≥3 independent experiments (data points shown from independent experiments) ± SEM. Asterisks indicate significance as assessed by one-way ANOVA, Bonferroni’s multiple comparisons test (*p≤0.05, **p≤0.01, ***p≤0.001, ****p ≤0.0001, n.s.; non-significant).

### Knockdown of βIII-tubulin in PDAC cells triggered death receptor 5 clustering

A key TRAIL receptor, death receptor 5 (DR5), requires clustering on the cell membrane for activation[28] and has been shown to bind to microtubules[29]. Using immunofluorescence, we consistently observed that βIII-tubulin knockdown, TRAIL, or combination treatment in PDAC cells triggered large membrane clusters of DR5 (**Figure 5A, Supplementary Figure 4A**). Quantification of DR5 cluster size showed that βIII-tubulin knockdown and TRAIL increased average cluster size compared to controls, with the greatest increase in cells with βIII-tubulin knockdown combined with TRAIL (**Figure 5B-C, Supplementary Figure 4B**). We evaluated the effect of βII-tubulin knockdown, which is also known to be upregulated in PDAC cells[16]. Unlike βIII-tubulin knockdown, βII-tubulin (upregulated in PDAC cancer cells) knockdown did not promote clustering of DR5 and did not increase the size of DR5 clusters (**Figure 5D, Supplementary Figure 4C**). Western blot for DR5 in MiaPaCa2 cells where surface proteins had been crosslinked, showed that βIII-tubulin silencing did not change expression of monomeric DR5 (between 37-50 kDa) but cells with βIII-tubulin siRNA+TRAIL had increased higher molecular-weight multimeric DR5 clusters (50-250 kDa), relative to controls (**Figure 5E, Supplementary Figure 5A**). Interestingly, βIII-tubulin silencing did not trigger clustering of the second TRAIL receptor, DR4, which rather showed a decrease in total DR4 expression with βIII-tubulin knockdown and no visible clusters of DR4 (**Supplementary Figures 5B-C**).

**Figure 5:**
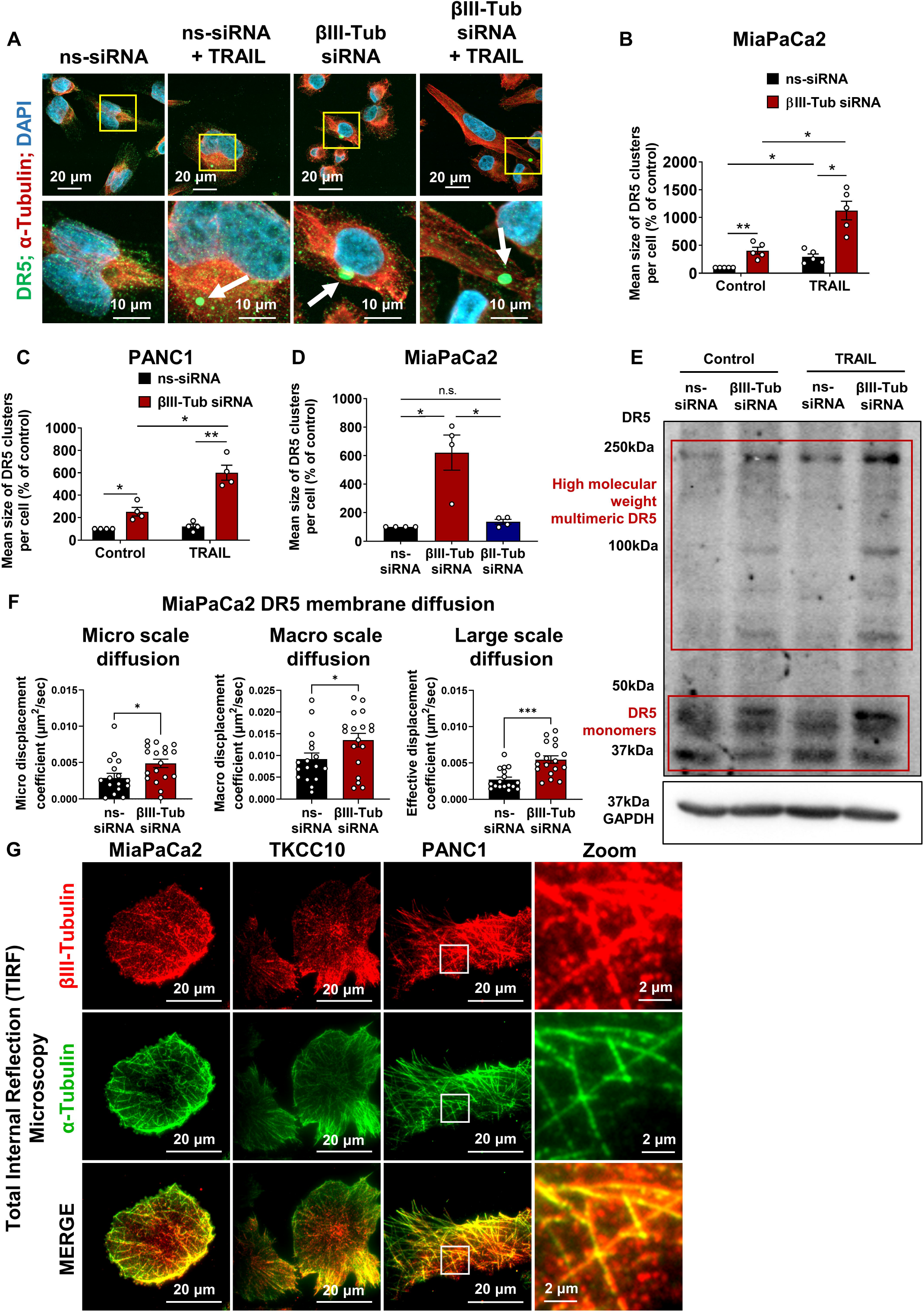
Knockdown of βIII-tubulin in PDAC cells triggered DR5 clustering. **(A)** Immunofluorescence staining for DR5 and α-tubulin in MiaPaCa2 cells transfected with non-silencing (ns) or βIII-tubulin (βIII-Tub) siRNA and treated ±TRAIL (250 ng/mL, 2 hours). βIII-tubulin knockdown and TRAIL triggered formation of large membrane clusters of DR5 (white arrows). Merged images shown again for reference in **Supplementary Figure 4A** with individual channel images. **(B-C)** Quantification of DR5 cluster size in MiaPaCa2 (n=5) **(B)** and PANC1 (n=4) **(C)** cells from Z-stack maximum intensity projections. **(D)** Quantification of DR5 cluster size in MiaPaCa2 cells transfected with ns-, βIII-tubulin, or βII-tubulin siRNA (n=4). **(E)** Western blot (non-reducing conditions) using protein extracted from MiaPaCa2 cells transfected with ns- or βIII-tubulin siRNA and treated ±TRAIL (250 ng/mL; 2 hours). Blot shown is representative of n=5 independent experiments; quantification shown in **Supplementary Figure 5A**. GAPDH used as a loading control. **(F)** Analysis of DR5 membrane diffusion using live imaging of GFP-tagged DR5 in MiaPaCa2 cells with total internal reflection (TIRF) microscopy and k-space image correlation spectroscopy. See **Supplementary Movie 2** for representative video of live cell imaging. Each data point indicates diffusion coefficient from analysis of a single cell. **(G)** Immunofluorescence for βIII-tubulin and α-tubulin and TIRF microscopy in PDAC cells. Bars represent mean of n≥3 independent experiments (data points shown from independent experiments) ± SEM. Asterisks indicate significance as assessed by one-way ANOVA, Bonferroni’s multiple comparisons test (**A, C**) or two-tailed t-test (**D, F**) (*p≤0.05, **p≤0.01, ***p≤0.001, n.s.; non-significant).

We subsequently assessed the effect of βIII-tubulin silencing on DR5 dynamics at the cell membrane, using MiaPaCa2 cells stably expressing GFP-DR5 and live total internal reflection (TIRF) microscopy[30]. Live imaging revealed clusters of DR5 at the membrane of cells that had been transfected with βIII-tubulin siRNA, relative to controls (**Supplementary Movie 1** and **Supplementary Figure 5D**). Importantly, DR5-GFP membrane clustering was associated with apoptotic features such as membrane blebbing (**Supplementary Movie 1** and **Supplementary Figure 5D**). To assess the movement of DR5 clusters at the cell membrane, we repeated TIRF live-cell imaging with higher temporal resolution (**Supplementary Movie 2**). We quantified DR5 dynamics and demonstrated increased diffusion coefficients across all spatial scales of movement with βIII-tubulin siRNA compared to controls (**Figure 5F**), consistent with faster movement of DR5 at the cell membrane. This suggested that βIII-tubulin silencing in PDAC cells increased DR5 membrane diffusion and increased chances of encountering other DR5 monomers leading to clustering and activation of cell death. To determine the feasibility of this hypothesis, we assessed if a βIII-tubulin cortical network is present at the cell membrane of PDAC cells. We identified βIII-tubulin containing microtubules that formed a mesh-like structure at the membrane of PDAC cells (**Figure 5G**), and βIII-tubulin knockdown almost completely abolished βIII-tubulin at the membrane (**Supplementary Figure 5E**). Architecture of membrane-associated α-tubulin was unaffected by βIII-tubulin knockdown (**Supplementary Figure 5F**). These results indicate that βIII-tubulin containing microtubules are present at the PDAC membrane and may be in a position to influence DR5 trafficking and clustering. We also observed a fraction of βIII-tubulin at the cell membrane that did not colocalise with α-tubulin and demonstrated punctate distribution (**Figure 5G**).

### Anti-tumour effects of βIII-tubulin silencing in PDAC cells are enhanced in the presence of cancer-associated fibroblasts

Based on our *in vitro* data showing that βIII-tubulin knockdown in PDAC cells enhanced TNFα-induced apoptosis, we questioned whether silencing of βIII-tubulin in PDAC cells would render them sensitive to CAF TNFα secretion. We co-cultured patient-derived CAFs with GFP-expressing PDAC cells and tracked GFP-positive PDAC cell proliferation following βIII-tubulin knockdown in tumour cells only. βIII-tubulin knockdown reduced proliferation of MiaPaCa2 cells (**Figure 6A** and **Supplementary Figure 6A**), patient-derived TKCC5, (**Supplementary Figure 6B**) and TKCC10 cells (**Supplementary Figure 6C**) when cultured in the presence of CAFs. Using the same culture model, we demonstrated that apoptosis induced by βIII-tubulin knockdown in MiaPaCa2 cells was enhanced in the presence of CAFs (**Figure 6B**). Similarly, immunofluorescence for cleaved caspase 8 in cytokeratin-positive PDAC cells in MiaPaCa2:CAF co-cultures similarly showed that βIII-tubulin knockdown in MiaPaCa2 cells cultured in the presence of CAFs significantly increased cleaved caspase 8 in PDAC cells, relative to those in the absence of CAFs (**Figure 6C-D**). TNFα neutralising antibodies blocked CAF-induced increase in caspase 8 cleavage in MiaPaCa2 cells with βIII-tubulin knockdown, indicating this effect was at least in part driven by TNFα secretions from CAFs (**Figure 6C-D**). These results were reproduced in TKCC10 PDAC cells (**Figure 6E**). Overall, these findings highlight that the pro-apoptotic and anti-tumour effects of βIII-tubulin silencing in PDAC cells are maintained and may be enhanced in the presence of CAFs.

**Figure 6:**
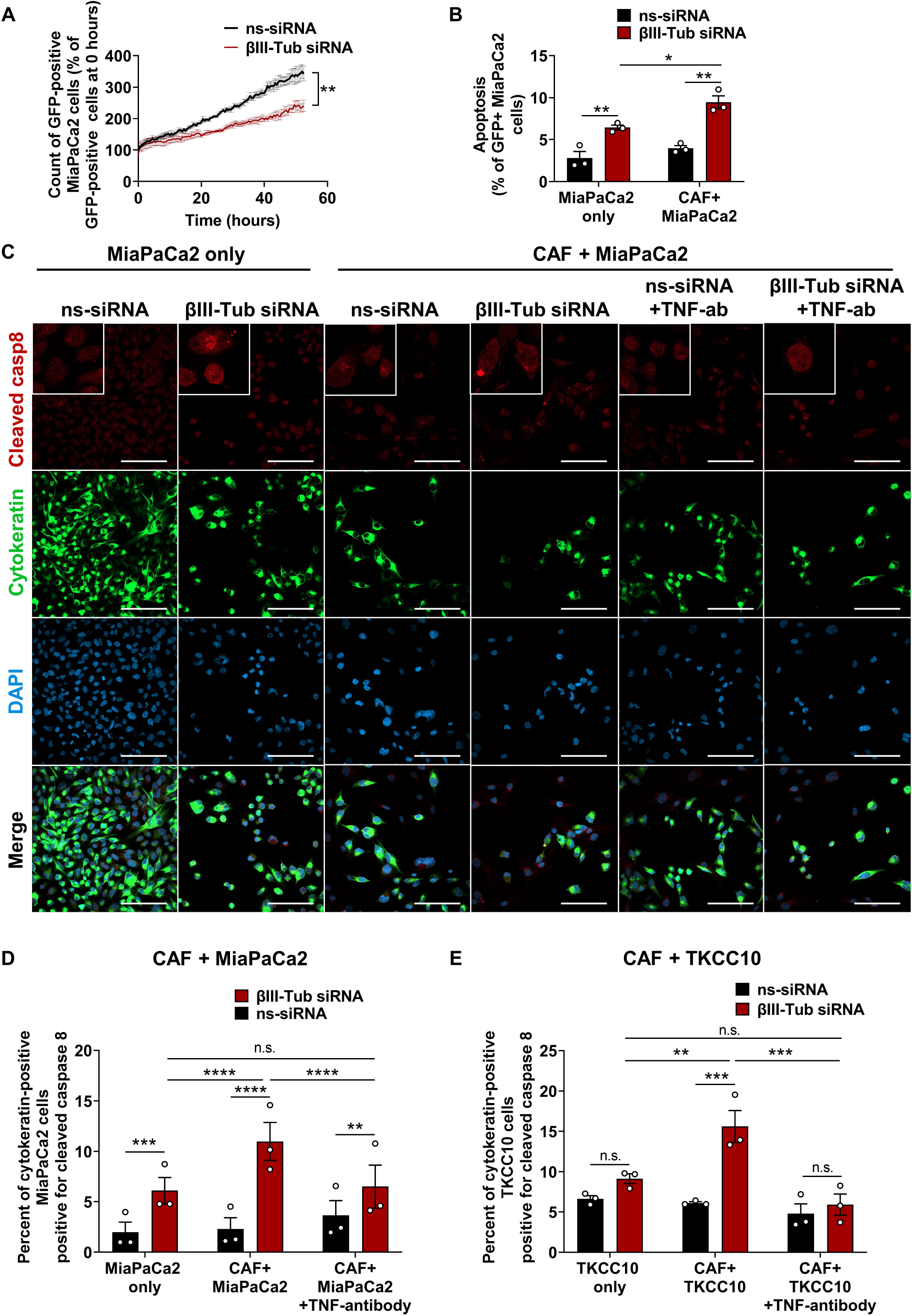
Anti-tumour effects of βIII-tubulin silencing are enhanced when PDAC cells are co-cultured with CAFs. **(A)** Live cell proliferation measured in GFP-MiaPaCa2 cells with non-silencing (ns) or βIII-tubulin (βIII-Tub) siRNA and co-cultured with cancer-associated fibroblasts (CAFs) (n=3 independent experiments with CAFs from 3 PDAC patients). **(B)** Apoptosis was measured using flow cytometry for AnnexinV/DAPI in GFP-MiaPaCa2 cells with βIII-tubulin knockdown and co-cultured with CAFs (n=3). **(C)** Immunofluorescence was used to measure the percentage of cleaved caspase 8 positive cells in cytokeratin positive PDAC cells when cultured alone or in the presence of CAFs. Cells were treated with or without TNFα-neutralising antibody (2 μg/mL). Representative images show immunofluorescence staining in MiaPaCa2 cells. All scale bars represent 100 μm. Equal number of total cells (MiaPaCa2 + CAFs) were seeded across each well. **(D-E)** QuPath software was used to measure the percentage of cleaved caspase 8 positive cells in cytokeratin positive MiaPaCa2 **(D)** or TKCC10 **(E)** cells (n=3). Bars represent mean of n≥3 independent experiments (data points shown from independent experiments) ± SEM. Asterisks indicate significance as assessed by one-way ANOVA, Bonferroni’s multiple comparisons test or two-tailed paired t-test (*p≤0.05, **p≤0.01, ***p≤0.001, ****p ≤0.0001, n.s.; non-significant).

### Combination of βIII-tubulin silencing and TRAIL treatment exerted potent anti-tumour effects in PDAC patient-derived tumour explants

Next, we examined the therapeutic potential of silencing βIII-tubulin in combination with TRAIL in a human patient-derived PDAC explant model which maintains 3D spatial multicellular organisation of PDAC tissue[27]. Star 3 nanoparticles + βIII-tubulin siRNA effectively into PDAC tumour explants and demonstrated potent βIII-tubulin protein knockdown (**Supplementary Figure 7A**-**C**).

**Figure 7:**
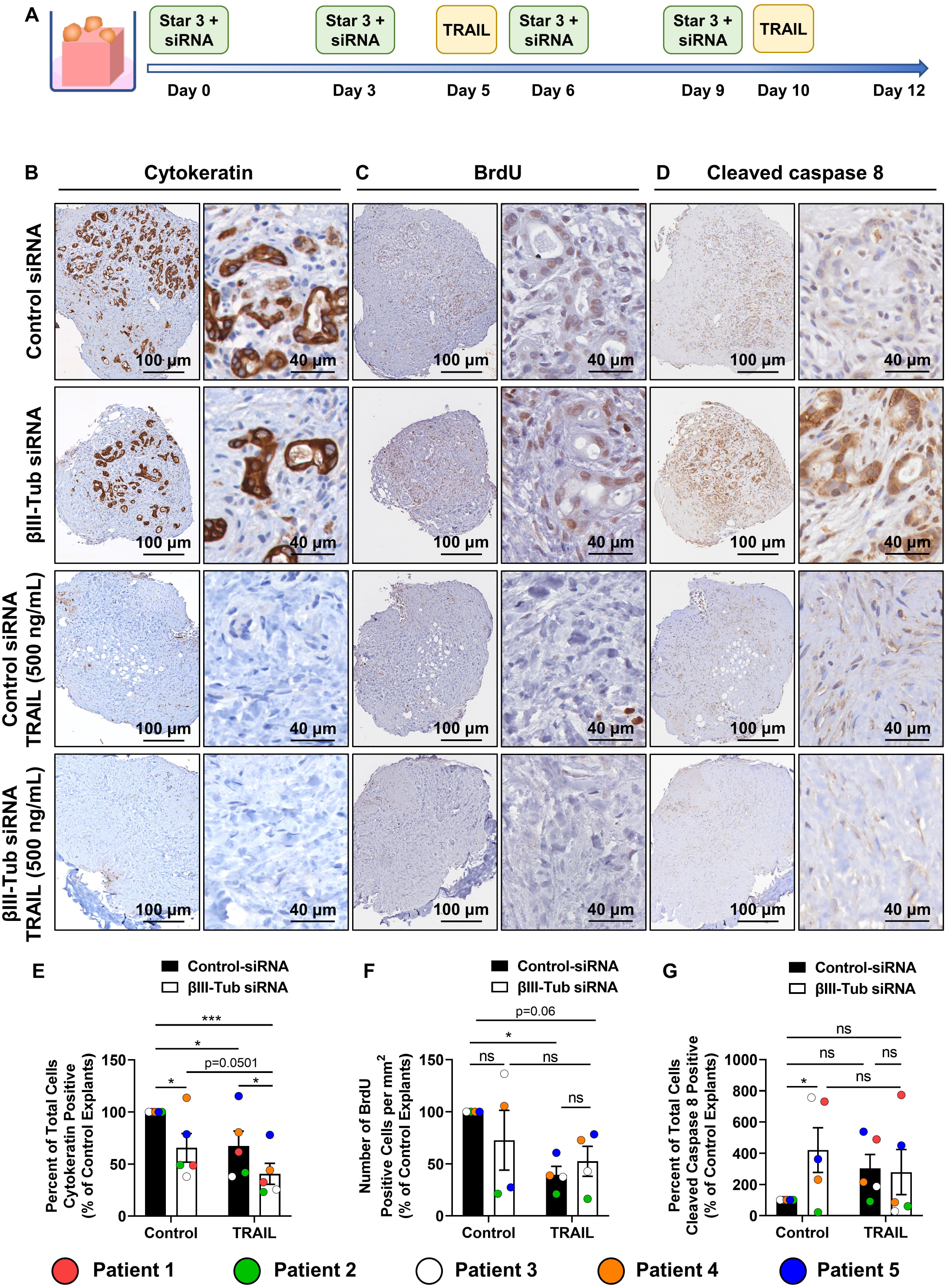
βIII-tubulin combined with TRAIL decreased tumour cell numbers in tumour explants from PDAC patients, with patient specific effects on extrinsic apoptosis and cell proliferation. **(A)** Treatment schedule was used to assess effects of βIII-tubulin (βIII-Tub) silencing combined with TRAIL in PDAC tumour explants. **(B-D)** Patient 3 representative immunohistochemistry staining of cytokeratin (tumour cell marker) **(B)**, bromodeoxyuridine (BrdU) (proliferation marker) **(C)**, and cleaved caspase 8 (extrinsic apoptosis marker) **(D)** at low and high magnification. **(E-G)** Quantification of staining from whole tumour explants was performed on QuPath for cytokeratin **(E)**, BrdU **(F)**, and cleaved caspase 8 **(G)** and data combined from n=5 patients taking the average quantification of 2-4 explants from each patient. Cell proliferation was not assessed in Patient 1. Each symbol represents the mean of 2-4 explants from each patient. Bars represent mean ± standard error of mean. Asterisks indicate significance as assessed by one-way ANOVA, Bonferroni’s multiple comparisons test (*p≤0.05, **p≤0.01, ***p≤0.001, ns: non-significant).

Explants from five PDAC patients were treated with Star 3+control-siRNA or βIII-tubulin siRNA and TRAIL over 12 days (**Figure 7A**). Positive βIII-tubulin staining in tumour cells was observed in control siRNA-treated explants from all 5 patients at endpoint (**Supplementary Figure 8**). Relative to tumour cell expression of βIII-tubulin, all 5 patients had low expression of βIII-tubulin in stromal cells (**Supplementary Figure 8**). Immunohistochemistry for cytokeratin (tumour cells), BrdU (proliferation), and cleaved caspase 8 (extrinsic apoptosis) was performed on endpoint explant tissue sections to determine therapeutic response (**Supplementary Figures 9-13** and representative images of cytokeratin, BrdU, and cleaved caspase 8 staining from Patient 3 are shown in **Figure 7B-D**).

**Figure 8.**
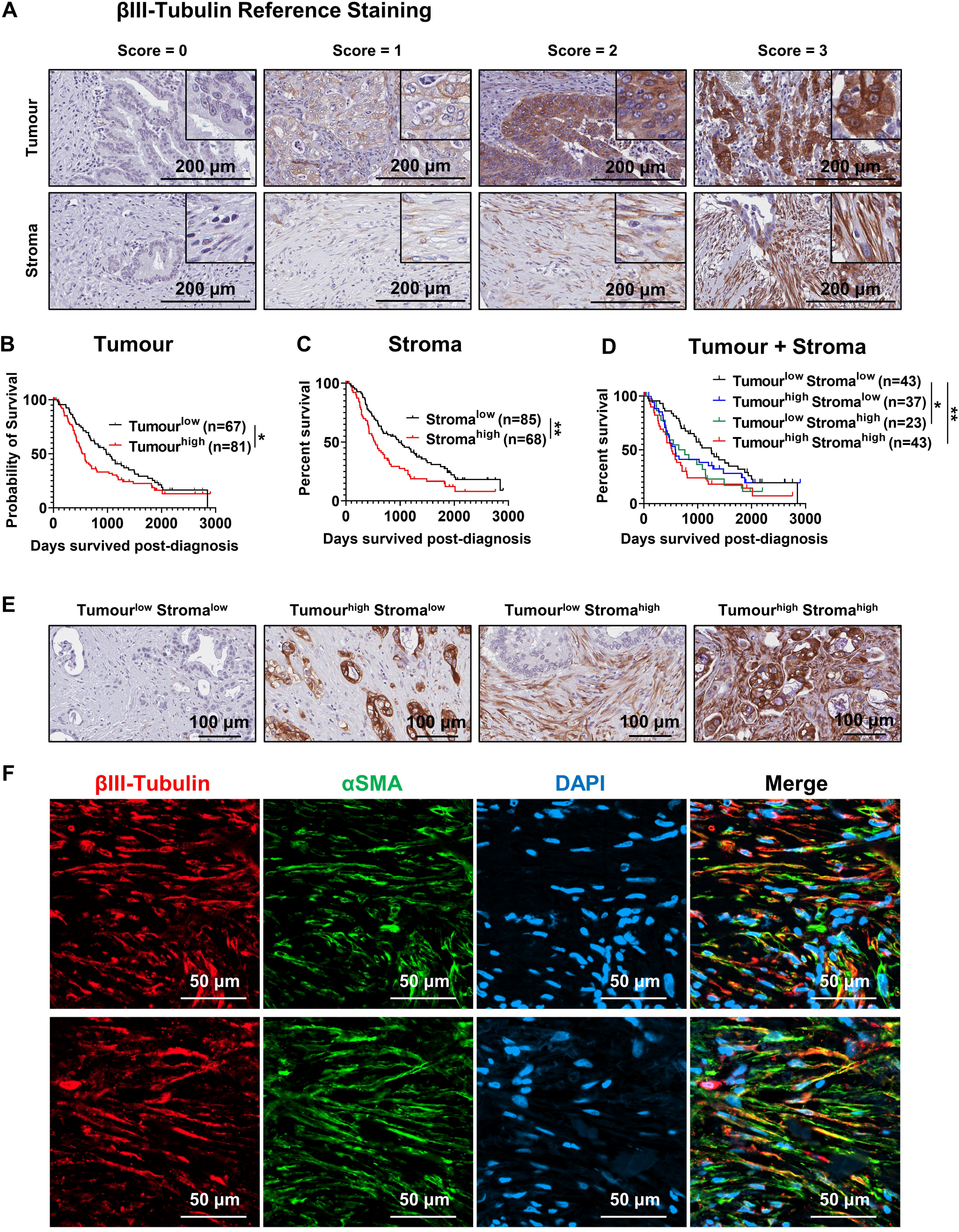
High βIII-tubulin expression correlates with PDAC patient poor overall survival. Human PDAC section obtained through the Australian Pancreatic Cancer Genome Initiative [International Cancer Genome Consortium Cohort (ICGC)] were stained for βIII-tubulin and tumour and stromal staining scored by three independent scorers. **(A)** Representative immunohistochemistry images showing four different scores of βIII-tubulin staining intensity in tumour and stromal regions. **(B-D)** Kaplan–Meier survival curves show correlation between βIII-tubulin expression in tumour cells **(B)**, stroma **(C)**, or both tumour and stroma **(D)** with overall patient survival (days post diagnosis). Total patient numbers per group are indicated in legends. Asterisks indicate significance based on univariate analysis, log-rank test (*p≤0.05; **p≤0.01). **(E)** Representative images showing examples of combined tumour and stroma score groups. **(F)** Co-immunofluorescence in 2 human PDAC samples to assess colocalisation of βIII-tubulin in α-smooth muscle actin (αSMA) positive cancer associated fibroblasts (CAFs).

Tumour cell frequency was reduced in 4/5 explants with βIII-tubulin silencing alone and TRAIL treatment alone compared to untreated control-siRNA explants (**Figure 7E**). Importantly, combination of βIII-tubulin silencing with TRAIL reduced tumour cell frequency in all the explants compared to untreated control-siRNA explants and led to the greatest reduction in tumour cell numbers compared to either treatment alone (**Figure 7E**).

Proliferating cell frequency (BrdU staining) was assessed in explants from Patients 2-5 but was not measured in Patient 1 explants given that they were not treated with BrdU substrate prior to fixation. Cell proliferation was reduced in explants from 2/4 patients treated with βIII-tubulin siRNA, 4/4 patients treated with TRAIL, and 4/4 patients treated with combination, compared to untreated control-siRNA explants (**Figure 7F**).

The frequency of cleaved caspase 8 positive cells increased in explants with βIII-tubulin silencing in 4/5 patients and increased with TRAIL treatment in 4/5 patients compared to untreated control-siRNA explants (**Figure 7G**). In combination treated explants, cleaved caspase 8-positive cell frequency increased in explants from 2/5 patients compared to untreated control-siRNA explants (**Figure 7G**).

In addition, Patient 6 explants were derived from a PDAC tumour metastasis to the stomach. Given the small size of the sample received from surgery, we only had enough tissue to perform βIII-tubulin silencing alone. In tumour explants with βIII-tubulin silencing compared to controls, there were no consistent effects on cytokeratin staining, but we observed a decrease in BrdU positive proliferative cells and an increase in cleaved caspase 8 positive cells (**Supplementary Figure 14A-C**).

We also investigated whether βIII-tubulin silencing or TRAIL treatment affected stromal CAF cells in PDAC explants. While there were no statistically significant differences in αSMA-positive CAF numbers with treatments across all 5 patients, we observed reduced CAF frequency in explants from 3/5 patients with βIII-tubulin silencing alone, and reduced CAF frequency in explants from 2/5 patients with TRAIL treatment alone or with combination treatment, compared to untreated control-siRNA explants (**Supplementary Figure 15A-B**).

### High βIII-tubulin expression in PDAC tumour cells correlated with poor overall survival and its high expression in the stroma was independently prognostic of poor overall-survival

We assessed the prognostic value of βIII-tubulin in tumour and stromal compartments of PDAC within the APGI International Cancer Genome Consortium (ICGC) PDAC cohort (**Figure 8**). Patient characteristics are described in **Supplementary Table 1**. 55% of patients had high intensity staining of βIII-tubulin in majority of PDAC tumour cells, and 44% patients had high intensity in stromal compartments. We showed that high expression of βIII-tubulin in tumour cells correlated with poor overall-survival (**Figure 8B** and **Supplementary Table 2**). Furthermore, we showed for the first time that high expression of βIII-tubulin in the stromal compartment of PDAC was independently prognostic (HR=2.163, p=0.009) of poor overall-survival (**Figure 8C** and **Supplementary Table 2**). When we combined tumour and stromal scores, we found that high expression of βIII-tubulin in both tumour and stroma was associated with the worst patient overall survival (**Figure 8D-E**). Given we showed that high stromal expression of βIII-tubulin is independently prognostic of poor overall survival, we used co-immunofluorescence in human PDAC samples to confirm that βIII-tubulin is expressed in α-smooth muscle actin (αSMA)-positive CAFs (**Figure 8F**).

## Discussion

The dismal prognosis of pancreatic cancer calls for development of novel treatment strategies. We identified βIII-tubulin, a component of the microtubule network, as a therapeutic target for PDAC[16]. βIII-tubulin has been shown to contribute to tumour progression and chemoresistance in several solid tumours[11, 16, 19, 22, 24-26]. However, the mechanism by which βIII-tubulin elicits its pro-survival effects was unknown. Moreover, there are currently no pharmacological inhibitors specific for βIII-tubulin. Here, we report for the first time that βIII-tubulin silencing is a novel activator of the extrinsic apoptotic pathway and can increase sensitivity to extrinsic apoptosis inducers. We also demonstrate proof-of-concept for application of nanoparticles carrying βIII-tubulin siRNA to selectively target and silence βIII-tubulin expression in mouse PDAC tumours and patient-derived PDAC tumour explants.

We demonstrated that both intrinsic and extrinsic apoptosis is activated by βIII-tubulin knockdown in PDAC cells. However, we showed that caspase 8 inhibition completely blocked apoptosis induced by βIII-tubulin knockdown, whereas caspase 9 inhibition had no effect, indicating βIII-tubulin specifically activates apoptosis via the extrinsic caspase 8-dependent pathway. While other studies have shown changes in survival pathways when βIII-tubulin is suppressed[19-21, 31], this is the first study to block a phenotype induced by βIII-tubulin knockdown by inhibiting a specific pathway.

Previously we showed that suppression of βIII-tubulin in PDAC cells stably expressing βIII-tubulin shRNA decreased tumour growth in mice[16]. To test effects of therapeutic βIII-tubulin suppression, we used polymeric nanoparticles (Star 3) that we demonstrated capable of delivering βIII-tubulin siRNA to orthotopic PDAC tumours[26]. These nanoparticles were recently shown to deliver βIII-tubulin siRNA to mouse orthotopic lung tumours and delayed tumour growth[32]. We demonstrated the therapeutic utility of βIII-tubulin silencing using nanoparticle-siRNA in PDAC with significantly reduced tumour volume and increased caspase 8 cleavage in tumours supporting our *in vitro* findings. The extrinsic apoptotic pathway can be activated by three ligands (TRAIL, TNFα, FasL) that bind to their own receptors and activate caspase 8-dependent apoptosis[33]. We showed that knockdown of βIII-tubulin in PDAC cells sensitised to all three ligands. Moreover, our results with TRAIL were reproduced in lung cancer cells where βIII-tubulin has also been linked to pro-tumour functions[19, 20, 24, 32]. Our findings suggest this may be a conserved βIII-tubulin function across other cancer types. This appears to be a cancer cell-specific mechanism as βIII-tubulin knockdown in PDAC patient-derived CAFs, did not increase apoptosis in the presence of TRAIL even though they express both βIII-tubulin and TRAIL receptors.

TNFα is a well-known inflammatory cytokine that binds to its receptor (TNF-R1) to induce apoptosis but is also capable of activating the NF-κB pathway to promote cell proliferation and tumour progression[34]. In the PDAC microenvironment, TNFα has been shown to be secreted by cancer-associated fibroblasts (CAFs; can make up the majority of cells in PDAC tumours) as a pro-tumour paracrine signal[35]. In co-cultures, we showed that anti-proliferative and pro-apoptotic effects induced by βIII-tubulin knockdown in PDAC cells are enhanced in the presence of CAFs at least in part via TNFα secretion from CAFs, thus potentially identifying an innovative therapeutic strategy to reprogram a microenvironmental pro-tumour signal into a death signal.

TRAIL is a chemotherapeutic capable of activating apoptosis specifically in tumour cells and demonstrated to be safe in several clinical trials[36]. Given TRAIL receptor DR5 binds to microtubules[29] and requires clustering to activate caspase 8[28], we examined the impact of βIII-tubulin knockdown on DR5 clustering. We demonstrated that βIII-tubulin knockdown triggered DR5 clustering that was enhanced in the presence of TRAIL. Knockdown of βII-tubulin, which is also upregulated in PDAC cells[16], failed to promote DR5 clustering, implying this could be specific to the βIII-tubulin isotype. We also observed that βIII-tubulin knockdown did not trigger DR4 clustering, highlighting a potential unique interaction between βIII-tubulin and DR5. Interestingly, βIII-tubulin silencing resulted in reduced protein expression of DR4 in PDAC cells, suggesting a potential compensatory downregulation of DR4 in response to DR5 activation.

We postulated that βIII-tubulin silencing triggered DR5 clustering and auto-activation, thus activating extrinsic apoptosis. We observed this with a live-cell imaging approach where we silenced βIII-tubulin and observed DR5 clustering in real time which preceded cell apoptosis. We also observed that DR5 clusters that were triggered by βIII-tubulin silencing showed high levels of lateral movement across the cell membrane, whereas in control cells, DR5 clusters were either non-existent or showed little to no movement. This suggests βIII-tubulin acts to impede movement of DR5 on the cell membrane and silencing of βIII-tubulin allows DR5 to freely diffuse to increase DR5 monomer encounters and subsequent clustering. In support of this, TIRF microscopy demonstrated that βIII-tubulin containing microtubules are present at the cell membrane of PDAC cells as a mesh-like architecture and may be in a position to influence DR5 trafficking and clustering. We also detected α-tubulin at the membrane of PDAC cells as previously reported[37], but α-tubulin membrane distribution was unaffected by βIII-tubulin knockdown. The mesh-like distribution of βIII-tubulin on the cell membrane of PDAC cells could hinder the transmembrane diffusion of DR5 by preventing the movement of its intracellular domain. When this hindrance is removed by silencing βIII-tubulin expression, we hypothesise this would allow DR5 to move freely across the cell membrane, increasing its chance of clustering with other DR5 receptors and activate cell death. This potential interaction warrants further investigation.

The ability to sensitise PDAC cells to TRAIL with βIII-tubulin suppression holds promise for PDAC therapy. TRAIL as an anti-cancer therapeutic has been investigated in clinical trials, where it was initially shown to be safe but failed due to a lack of therapeutic response due to the poor bioavailability of TRAIL protein and development of resistance[36, 38]. Here, we have described a novel way to sensitise PDAC cells to TRAIL-induced apoptosis, thus potentially overcoming the challenge of TRAIL resistance. However, to evaluate clinical potential of therapeutic targeting of βIII-tubulin and TRAIL for PDAC treatment, we wanted to test this combination in a model that mimics the complexity and heterogeneity of the human PDAC microenvironment. We used our patient derived PDAC tumour explant model which accurately maintains the complexity, multicellular architecture, and cross-cell interactions of human PDAC stroma[27].

In all patient explants tested, silencing of βIII-tubulin alone demonstrated potent anti-tumour cell effects. In 4/5 patients, silencing of βIII-tubulin reduced the number of tumour cells in the explants, and in patient explants with no reduction in tumour cell frequency, we observed increased extrinsic apoptosis. This suggests that all patient explants were sensitive to βIII-tubulin silencing alone, albeit to different degrees between patients reflecting inter-patient tumour heterogeneity. That is, more sensitive patient tumour explants would have lost tumour cells early in the 12-day culture window, whereas less sensitive tumour explants may have taken longer to succumb to the cell death induced by βIII-tubulin silencing. Similarly, all patient explants demonstrated an anti-tumour response to TRAIL. Tumour cell numbers were reduced in 4/5 patients, and in patient explants with no reduction in tumour cell numbers, we observed increased extrinsic apoptosis and decreased cell proliferation. These results are promising, especially given that TRAIL phase III clinical trials failed due to lack of clinical benefit[38]. Our results also suggest that lack of clinical benefit in TRAIL trials could be due, at least in part, to poor bioavailability of TRAIL resulting in its poor access to the tumour[38], whereas TRAIL would have had more direct access into tumour explants. Studies that aim to improve the bioavailability of TRAIL using nanoparticle drug carriers or antibody approaches are thus warranted. Combination treatment of βIII-tubulin silencing and TRAIL resulted in a more pronounced reduction in tumour cell numbers compared to either treatment alone. There were no consistent differences in caspase 8 cleavage or cell proliferation with combination treatment compared to either treatment alone, highlighting varying levels of response due to different degrees of sensitivity to treatments between each patient. We also demonstrated anti-tumour effects and induction of extrinsic apoptosis in tumour explants from a patient with metastatic PDAC, providing evidence that βIII-tubulin silencing could also be effective in patients with metastatic disease. These results highlight the promising therapeutic potential of targeting βIII-tubulin to release the brakes on extrinsic cell-death signals and raises the possibility of combining such a therapeutic strategy with TRAIL treatment.

Finally, we assessed the proportion of PDAC patients that have upregulation of βIII-tubulin and whether this correlates with patient outcome. βIII-tubulin had been previously shown to be overexpressed in surgically resectable PDAC patients[17], and in patients with metastatic PDAC, high expression of βIII-tubulin was associated with decreased progression free survival[18]. A recent study found that high βIII-tubulin protein expression correlated with poor progression-free survival in PDAC patients receiving gemcitabine and Abraxane chemotherapy[39]. Our study, in a cohort of surgically-resectable PDAC patients, strengthens these findings as this is the largest PDAC cohort in which this correlation has been demonstrated and the first time that high expression of βIII-tubulin has been correlated with decreased overall-survival in PDAC. Importantly, using multivariate analysis, we also show for the first time that high expression of βIII-tubulin in the stromal compartment of PDAC is independently prognostic of poor overall survival. When we combined the tumour and stromal score, we found that high expression of βIII-tubulin in both tumour and stroma is associated with the worst patient overall-survival. This opens a previously unexplored avenue to investigate the functional role of βIII-tubulin in stromal cells of PDAC. In this study, we demonstrated that silencing of βIII-tubulin in CAFs *in vitro* reduced their proliferation, and in patient-derived tumour explants, βIII-tubulin silencing led to reduced CAF cell frequency in 3/5 patients. However, in contrast with PDAC cells, βIII-tubulin did not appear to regulate extrinsic apoptosis and TRAIL sensitivity in CAFs, requiring future work to further probe the functional role of βIII-tubulin in PDAC CAFs. Given we observed patient specific effects of βIII-tubulin silencing on CAFs, future work should also investigate whether subtypes of PDAC CAFs are more responsive to βIII-tubulin inhibition. Recent studies have uncovered the complexity of CAF cell heterogeneity within the PDAC stroma and the important functional roles played by different CAF subtypes in PDAC[7, 40-42]. Future work in larger patient cohorts should investigate how inhibition of βIII-tubulin in PDAC cells and CAFs are influenced by different stromal subtypes and whether particular stromal signatures can predict response to βIII-tubulin targeting therapy.

Taken together, this work represents a major breakthrough in understanding βIII-tubulin biology and developing therapeutic strategies to inhibit βIII-tubulin. It has provided several new insights into the pro-survival role of βIII-tubulin in PDAC and more broadly, its role as a therapeutic target in cancer. For decades, βIII-tubulin has been established as a key cancer therapeutic target[19-21, 24, 25, 31, 43-50]. Prior to this work however, the precise pro-survival pathways controlled by βIII-tubulin in cancer cells were poorly understood. We discovered an unexpected role of βIII-tubulin as a brake on extrinsic cell death. These findings provide momentum to develop βIII-tubulin targeting therapies, especially in stromal rich cancers such as PDAC where the complex role of the tumour microenvironment can be seen as an opportunity to provide a source of extrinsic apoptotic signals to cancer cells where βIII-tubulin has been supressed. This new insight demonstrates the clinical utility of βIII-tubulin silencing using gene silencing nanomedicines as a multi-pronged attack on tumour cells and paves the way for the translation of βIII-tubulin targeting therapies for PDAC.

## Materials and Methods

All tissue culture reagents were purchased from Gibco by Life Technologies. All siRNAs were purchased from Dharmacon, Horizon Discovery or Qiagen.

### Cell culture

Human PDAC (MiaPaCa2, PANC1, and HPAF-II) and non-small cell lung carcinoma (NSCLC) (H460) cell lines were obtained from American Type Culture Collection (ATCC). MiaPaCa2 cells were grown in Dulbecco’s Modified Eagle Medium (DMEM) supplemented with 10% foetal bovine serum (FBS), 2.5% horse serum and 2 mM GlutaMAX. PANC1 cells were grown in DMEM supplemented with 10% FBS and 2 mM GlutaMAX. HPAF-II cells were grown in Eagle’s Minimum Essential Medium (MEM) supplemented with 10% FBS, 2 mM GlutaMAX and 1 mM sodium pyruvate. H460 cells were cultured in RPMI medium supplemented with 10% FBS and 2 mM GlutaMAX. Patient-derived TKCC cells were isolated from patient-derived xenografts, as previously described [51], and were a gift from A/Prof Marina Pajic from the Garvan Institute of Medical Research. TKCC5 cells were grown in DMEM mixed 1:1 with Ham’s F12 medium, supplemented with 7.5% FBS, 2 mM GlutaMAX, 15 mM HEPES, 1.2% glucose, 25 μg/mL apo-transferrin, 40 ng/mL hydrocortisone, 0.1 IU/mL insulin and 10 ng/mL human recombinant epidermal growth factor (EGF) (all from Sigma-Aldrich). TKCC10 cells were grown in a 1:1 mixture of M199 medium and Ham’s F12 medium, supplemented with 7.5% FBS, 1x MEM vitamins, 2 mM GlutaMAX, 15 mM HEPES, 0.06% glucose, 25 ng/mL apo-transferrin, 0.2 IU/mL insulin, 40 ng/mL hydrocortisone, 20 ng/mL EGF, 0.5 pg/mL Tri-iodothyronine, and 2 μg/mL O-phosphoryl ethanolamine. Patient derived CAFs were isolated from PDAC tumours by explant outgrowth culture and used within 12 passages as previously described [52, 53]. CAFs were grown in Iscove’s modified Dulbecco medium, 10% FBS, and 4 mmol/L GlutaMAX. The purity of CAFs was assessed by positive staining for glial fibrillary acidic protein (GFAP) and α-smooth muscle actin (αSMA) and negative staining for cytokeratin, as described previously [54]. All experiments using patient derived CAFs were approved by UNSW Sydney human ethics committee (approvals: HC14039, HC180973) and all experiments were performed in accordance with the relevant guidelines and regulations. All patients provided written informed consent.

All cells were maintained in 75 cm^2^ flasks at 37 °C in a humidified incubator with 5% CO2. For experiments described below, PDAC and NSCLC cells were lifted with 0.25% trypsin/0.53 mM EDTA, counted with an automated cell counter (Bio-Rad Laboratories) and re-seeded at equal densities in 6- or 96-well plates. Patient derived CAFs were lifted with 0.05% trypsin/0.53 mM EDTA. All cells were monthly tested negative for mycoplasma.

### siRNA transfection

PDAC and NSCLC cells were transfected with siRNA 24-hours post seeding (MiaPaCa2, PANC1, HPAF-II and TKCC cells: 10^5^ cells/well; H460 cells: 6×10^4^ cells/well) in a 6-well plate, as previously described [55, 56]. ON-TARGET*plus* SMARTpool siRNA specific for βIII-tubulin (Dharmacon, cat. L-020099-00) and βII-tubulin (Dharmacon, cat. L-008260-00) were used. ON-TARGET*plus* Non-Targeting Control Pool siRNA (Dharmacon, cat. D-001810-10-20), referred to as non-silencing siRNA (ns-siRNA) was used as a negative control. Results using SMARTpool siRNA were validated using an independent single-sequence siRNA targeting βIII-tubulin (Qiagen, cat. 1027418). MiaPaCa2 cells were re-seeded 1:3, 24 hours post-transfection.

### Quantitative real-time PCR

Total cellular RNA was extracted from transfected PDAC cells using the RNeasy Mini Plus Kit (Qiagen), according to the manufacturer’s instructions. RNA purity was tested on a Nanodrop spectrophotometer using spectrophotometric absorbance ratios at wavelengths of 260 nm and 280 nm (A260/A280). All RNA samples had an A260/A280 ratio of >1.8. For production of complementary DNA (cDNA), 500 ng RNA was reverse transcribed using a High-Capacity cDNA Reverse Transcription Kit (Applied Biosystems), according to the manufacturer’s instructions. mRNA levels of βIII-tubulin were then quantified with Quantifast SYBR Green PCR Kit (Qiagen) with 40 cycles on the ViiA 7 Real-Time PCR machine (Life Technologies). βIII-tubulin forward (GeneWorks, cat. 730175) and reverse (GeneWorks, cat. 730176) primers were used. 18S ribosomal RNA (rRNA) (Qiagen, cat. QT00199367) was used as a housekeeping gene to normalise βIII-tubulin mRNA levels.

### Western blot analysis

Total cellular protein was harvested from transfected PDAC and NSCLC cells 72 hours post-transfection using Cell Lysis Buffer (Cell Signalling Technology) supplemented with 1 mM phenylmethanesulfonylfluoride (PMSF). Protein extract concentrations were determined using a Pierce BCA Protein Assay Kit (ThermoFisher Scientific) according to the manufacturer’s instructions. 10 μg protein and standards were loaded onto a 10% SDS polyacrylamide gel. Protein samples were electrophoresed in a Mini-PROTEAN Tetra Cell system, then separated proteins were transferred to a 0.45 μm nitrocellulose membrane (Bio-Rad Laboratories). Membranes were incubated in blocking buffer containing 5% skim milk in PBS/0.1% Tween-20 (PBST) then incubated with their respective primary antibodies diluted in 5% skim milk in PBST at 4 °C overnight: purified mouse anti-βIII-tubulin (1:1,000) (BioLegend, cat. 801202), monoclonal mouse anti-GAPDH (1:50,000) (abcam, cat. Ab8245), monoclonal rabbit anti-DR5 (1:1000) (Cell Signalling Technology, cat. 8074), or monoclonal rabbit anti-DR4 (1:1000) (Cell Signalling Technology, cat. 42533). Membranes were then incubated for 1 hour at room temperature in the HRP-labelled polyclonal goat anti-mouse IgG (DAKO, cat. P0447) or HRP-labelled polyclonal goat anti-rabbit IgG (DAKO, cat. P0448) diluted in 5% skim milk/PBST. Protein bands were visualised using Amersham ECL Western Blotting Detection Reagent and the ImageQuant LAS4000 luminometer (GE Healthcare Life Sciences). βIII-tubulin relative band intensities were quantified using ImageStudio software (LI-COR Biosciences) and normalised to GAPDH expression.

For western blot analysis of TRAIL-receptor clustering, MiaPaCa2 cells were treated with or without TRAIL at 250 ng/mL for 2 hours, 72 hours post-transfection. Following TRAIL treatment, cells were washed in 1x PBS three times, then protein was crosslinked by adding 2 mM BS3 crosslinker into each well and incubated for 45 minutes at room temperature. Following 45-minute crosslinking, BS3 was quenched by adding 40 mM Tris-HCl at pH 7.5 for 10 minutes, then protein was extracted in cell lysis buffer supplemented with 1 mM PMSF as described above. Western blot was performed under non-denaturing conditions, and separated proteins were transferred to a polyvinylidene fluoride (PVDF) membrane for 2 hours at 100 V. Membranes were incubated in blocking buffer containing 5% skim milk in PBST then incubated with primary antibodies diluted in 5% skim milk in PBST at 4 °C overnight: (i) monoclonal rabbit anti-DR5 (1:1000) (Cell Signalling, cat. 8074), (ii) monoclonal rabbit anti-DR4 (1:1000) (Cell Signalling, cat. 42533), (ii) monoclonal mouse anti-GAPDH (1:50,000) (abcam, cat. Ab8245). Membranes were then incubated for 1 hour at room temperature in the HRP-labelled polyclonal goat anti-rabbit IgG (DAKO, cat. P0448) or HRP-labelled polyclonal goat anti-mouse IgG (DAKO, cat. P0447) diluted in 5% skim milk/PBST. Protein bands were visualised using Biorad Clarity Max ECL reagent and the iBright luminometer (ThermoFisher Scientific). Quantification of multimeric DR5 was performed on the iBright analysis software by quantifying DR5 bands between 50-250 kDa and normalising to GAPDH expression.

### Quantification of apoptosis

Apoptosis was measured using flow cytometry for Annexin V-PE (BD Biosciences) and DAPI (Sigma-Aldrich, cat. D9542), as previously described [55, 56]. Floating and adherent cells were pelleted in a centrifuge at 230g for 3 minutes, then 10^5^ cells were resuspended in 1x binding buffer containing Annexin V-PE and 100 μg/mL DAPI. Cells were incubated in the dark for 15 minutes at room temperature then analysed on a FortessaSORP flow cytometer (BD Biosciences). The distribution of cells in early, middle, to late apoptosis was quantified using FlowJo v10 software to determine the total proportions of apoptosis in each sample. An example of the gating strategy used for quantification of apoptosis is shown in **Supplementary Figure 16A**.

### Quantification of cell viability and cell proliferation

To quantify cell viability, cells were harvested and pelleted at 335g for 3 minutes. Cells were resuspended in culture medium and mixed 1:1 with trypan blue solution (ThermoFisher Scientific). Live cells were counted on an automated cell counter (Bio-Rad Laboratories).

To measure real-time cell proliferation, cells were placed in an IncuCyte® S3 Live-Cell Analysis System (Essen BioScience). Each well of a 6-well plate was scanned every hour for a total of 48 hours with a 10x objective (NA=0.3). Phase-contrast images and integrated cell analysis metrics were used to calculate cell proliferation expressed as percent confluence or total cell count using the cell-by-cell analysis platform on IncuCyte® software. An example phase contrast image of MiaPaCa2 cells on the IncuCyte® S3 is shown in **Supplementary Figure 17A** and the corresponding confluence mask (yellow) applied to the phase contrast image is shown in **Supplementary Figure 17B**.

### Drug and inhibitor treatment

Specific pharmacological inhibitors of caspase 9 (Z-LEHD-FMK; Cat. 1149-1, BioVision) and caspase 8 (Q-IETD-OPh; Cat. 1176-1, BioVision) were used to treat MiaPaCa2 (20 μM inhibitors), PANC1 (50 μM inhibitors) and H460 (50 μM inhibitors) cells, 48 hours post-transfection with ns-siRNA or βIII-tubulin siRNA. Apoptosis was measured 24 hours post-treatment with the inhibitor. The inhibitors were used at doses previously published in PDAC cells [57]. Both inhibitors were dissolved in dimethyl sulfoxide (DMSO) and DMSO vehicle was used as the 0 μM control. We confirmed that these inhibitors potently inhibit their target caspase by demonstrating that the caspase 9 inhibitor blocked paclitaxel induced caspase 9 activity (**Supplementary Figure 18A**) and the caspase 8 inhibitor blocked TRAIL induced apoptosis (**Supplementary Figure 18B**) in MiaPaCa2 cells. Caspase 9 activity in cells treated with caspase 9 inhibitor was recorded as zero given the inhibitor binds to the active site of the enzyme used in the CaspaseGlo9 assay. To test whether caspase 8 inhibition can reverse βIII-tubulin knockdown and TRAIL-induced apoptosis, cells were pre-incubated with caspase 8 inhibitor for 1 hour then treated with TRAIL (10 ng/mL in MiaPaCa2; 2.5 ng/mL in H460), 48 hours post-transfection.

To determine the effect of βIII-tubulin silencing on the sensitivity to TRAIL, cells were treated 48 hours post-transfection with human recombinant TRAIL (MiaPaCa2: 5-20 ng/mL; PANC1: 5-20 ng/mL; TKCC5: 2.5 ng/mL; TKCC10: 20 ng/mL; H460: 2.5 ng/mL) (Merck-Millipore, cat. GF092). TRAIL doses in each cell line were chosen at a concentration that resulted in at least a 50% increase in apoptosis. TRAIL was dissolved in sterile H_2_O according to the manufacturer’s instructions. Cells were harvested for measurement of apoptosis or cell viability 24 hours post-treatment with TRAIL. Similarly, cells were treated with human recombinant TNFα (abcam, cat. ab9642) 48 hours post transfection, and apoptosis measured 24 hours after treatment with TNFα (MiaPaCa2: 10 ng/mL; PANC1: 10 ng/mL; TKCC5: 20 ng/mL; TKCC10: 20 ng/mL). TNFα was dissolved in sterile H_2_O according to manufacturer’s instructions. PDAC cells were treated with human recombinant FasL (MiaPaCa2: 50 ng/mL; TKCC10: 100 ng/mL) (Sigma-Aldrich, cat. SRP3036), 48 hours post transfection, and apoptosis was measured 48 hours post-treatment with FasL in MiaPaCa2 cells, and 24 hours post-treatment with FasL in TKCC10 cells. FasL was dissolved in sterile 1x PBS containing 0.1% bovine serum albumin, as per manufacturer’s instructions.

### Measurement of caspase activity

To measure caspase activity, MiaPaCa2 cells were seeded into both clear and white-walled 96-well plates 24 hours post-transfection at 5000 cells/well, as previously described [16]. 72 hours post-transfection, CaspaseGlo assays were performed in white-walled 96-well plates to measure the activity of caspase 9 (cat. G8210) or caspase 8 (cat. G8200) (both from Promega) according to manufacturer’s instructions. Luminescence was measured using the GloMax 96 Microplate Luminometer (Promega). In parallel, Cell-Counting Kit-8 (CCK-8, Sigma-Aldrich) was used to quantify cell viability so that caspase activity could be normalised to cell number. CCK-8 substrate was added to each well of the clear-walled 96-well plate, incubated at 37 °C for 1 hour, then absorbance was measured at 450 nm on a SpectraMax 190 Microplate Reader (Molecular Devices).

### Generation of βIII-tubulin stable short hairpin RNA (shRNA)-luciferase expressing cells

MiaPaCa2 cells stably expressing pGL4.50 (Mammalian Luciferase, Promega) for luciferase expression, and βIII-tubulin shRNA or scramble shRNA were established as previously described [16].

### Nanoparticle synthesis and preparation

The synthesis of Star 3 nanoparticles was carried out as previously described using the reversible addition-fragmentation chain transfer polymerisation technique [26]. Star 3 is a miktoarm star polymer consisting of cationic poly(dimethylaminoethyl methacrylate) (PDMAEMA) arms to electrostatically self-assemble siRNA and poly[oligo(ethylene glycol) methyl ether methacrylate] (POEGMA) arms to help shield the positive charge of the nanoparticles [26]. When complexed with siRNA, Star 3 nanoparticles have a size of around 35 nm [26].

To prepare Star 3 nanoparticles and siRNA complexes for *in vivo* use, freeze-dried Star 3 nanoparticles were dissolved in ultrapure H_2_O at 2.4 mg/mL and passed through a 0.45 μm polyethersulfone (PES) filter. Star 3 + siRNA complexes were prepared by diluting siRNA in ultrapure H_2_O for a final amount of 4 mg/kg siRNA per mouse and adding to 120 μg Star 3. Star 3 + siRNA was allowed to complex for 5 minutes at room temperature and then immediately injected intravenously into the mouse.

### Orthotopic pancreatic cancer mouse model

8-week-old female BALB/c nude mice were used. All animal experiments were approved by the Animal Ethics committee, UNSW (ACEC 12/7B; 13/130B; 18/54B). 1x 10^6^ MiaPaCa2 cells (with stable expression of luciferase) were implanted into the tail of the pancreas of mice as previously described [16]. PDAC tumours were allowed to develop for 4 weeks. Mice were then randomised based on tumour size (to obtain an equal average starting tumour size per treatment group, **Figure 2B**), as determined by administration of 150 mg/kg D-luciferin and measurement of tumour luminescence on an IVIS SpectrumCT imager (PerkinElmer). Star 3 + control siRNA (antisense: 5′-GAACUUCAGGGUCAGCUUGCCG) or βIII-tubulin siRNA (antisense: 5′-GCAGUUUUCACACUCCUUCUU) was then administered intravenously at 4 mg/kg twice weekly for 4 weeks. Mice were humanely sacrificed and PDAC tumours were harvested. Primary tumour volume was measured using micro-callipers and the formula volume = (length x width x height)/2. PDAC tumour tissue was also collected for assessment of βIII-tubulin knockdown by western blot and immunohistochemistry. PDAC tumour tissue was fixed in 4% paraformaldehyde for 24 hours (Electron Microscopy Sciences, cat. 15710) for immunohistochemistry analysis. 30 mg of snap-frozen PDAC tumour tissue was homogenised in 1 mL of cold 1x RIPA buffer containing 1% protease/phosphatase inhibitor cocktails (Sigma-Aldrich, cat. P8340 and cat. P5726, respectively) using a Qiagen TissueRuptor (cat. 990890). Homogenisation was followed by sonication at 4 °C. Samples were then incubated on ice for 30 minutes, before centrifugation at 14000g for 10 minutes at 4 °C. Protein-containing supernatant was frozen (−80 °C) and used for downstream western blot as described above in section 9.4.

### Immunohistochemistry analysis of mouse orthotopic pancreatic tumour sections

Orthotopic PDAC tumours were collected at the conclusion of orthotopic models by surgical removal. Paraffin-embedded tissue sections were stained with βIII-tubulin antibody (1:50; Biolegend, cat. 801202) using a mouse-on-mouse kit (Vector Laboratories, cat. BMK-2202) according to the manufacturer’s instructions. Cleaved caspase 8 immunohistochemistry staining was performed using cleaved caspase 8 primary antibody (1:100; Cell Signalling Technology, cat. #9496) and goat anti-rabbit biotinylated secondary (1:200; Vector Laboratories, cat. BA-1000). Briefly, tissue sections were deparaffinised at 60 °C for 30 minutes then rehydrated through consecutive washes in xylene, ethanol, and water. Antigen retrieval was performed by microwaving slides for 4 minutes in 10 mM citrate buffer + 0.05% Tween-20 at pH6.0, followed by a 15-minute incubation at 104 °C. Non-specific peroxidase activity was blocked with 1% hydrogen peroxide + 1% methanol for 10 minutes at room temperature. After blocking in 10% goat serum, tissue samples were stained with cleaved caspase 8 primary antibody diluted 1:100 and incubated overnight at 4 °C. Biotinylated anti-rabbit secondary antibody (1:200) was followed by incubation with Vectastain® ABC kit (Vector laboratories). 3,3’ diaminobenxidine (DAB) was used as the substrate, and tissues were counterstained with hematoxylin. All stained tissue sections were scanned on a Vectra Polaris (PerkinElmer) slide scanner using a 40x/0.75 NA objective. Quantification of cleaved caspase 8 staining was performed on QuPath v0.3.2 [58] to count the percent of total cells cleaved caspase 8 positive in whole scanned tumour sections. Isotype control antibodies were used at the same concentration as primary antibodies (Mouse IgG2A for βIII-tubulin stain; Rabbit IgG for cleaved caspase 8 stain; **Supplementary Figure 19**).

### Clonogenic assays

24 hours post-transfection, MiaPaCa2 cells were seeded at 300 cells/well, and HPAF-II cells at 500 cells/well in 6-well tissue culture plates, as previously described [16]. 24 hours post-seeding, cells were incubated with TRAIL (MiaPaCa2: 5 and 10 ng/mL; HPAF-II: 50 and 200 ng/mL) for a total of 72 hours. Colonies were stained with crystal violet 9 days post-seeding for MiaPaCa2 cells and 14 days post-seeding for HPAF-II cells, imaged with epi-illumination on an ImageQuant LAS4000 (GE Healthcare Life Sciences), and counted using the ImageQuantTL software (GE Healthcare Life Sciences).

### Anoikis assays

Anoikis assays were performed as previously described [16]. Briefly, two coats of poly 2-hyroxyethyl methacrylate (Poly-HEMA) (12 mg/mL, dissolved in 95% ethanol) were added to 6-well tissue culture plates and left to dry overnight at room temperature. MiaPaCa2 cells were seeded into 6-well tissue culture plates at 100,000 cells/well, then transfected with ns-siRNA or βIII-tubulin siRNA. 24 hours post-transfection, MiaPaCa2 cells were then re-seeded into the Poly-HEMA coated plates. 48 hours post-transfection, cells were cultured with or without TRAIL (10 ng/mL) for a further 9 hours then apoptosis was measured by annexin V/DAPI staining and flow cytometry, as described above (Section 9.5).

### TRAIL receptor immunofluorescence

MiaPaCa2 and PANC1 cells were re-seeded into 8-well chamber slides (Ibidi, cat. 80826) at 10,000 cells/well, 24 hours post transfection. 72 hours post-transfection, cells were treated with TRAIL (250ng/mL) for 2 hours, then fixed in 4% paraformaldehyde for 10 minutes. The 2-hour TRAIL treatment was chosen as an early timepoint to allow us to observe early changes in DR5 prior to the cell becoming apoptotic. Immunofluorescence staining was performed as previously described [16], using primary antibodies for DR5 (1:100; Cell Signalling Technologies, cat. 8074) and α-tubulin (1:500; Sigma-Aldrich, cat. T9026). Secondary antibodies used were goat anti-rabbit AF488 (1:500; Molecular Probes, cat. A-11008) and goat anti-mouse AF647 (1:500; abcam, cat. ab150115). Stained cells were mounted with Prolong Gold anti-fade mounting media with DAPI to visualise cell nuclei. Images were taken on a Zeiss LSM800 confocal microscope. For quantification of DR5 cluster size, z-stack images (7 slices at 3 μm) were obtained using a 40x/1.3 NA objective to capture an average of 20 cells per field of view, and 3-5 representative images were captured for each sample. Maximum intensity projections of the z-stack images were processed using Zeiss Zen Black 2.3 software, and DR5 cluster size was measured using the Analyse Particle function on ImageJ v1.52a (National Institutes of Health, Bethesda, Maryland, USA). DR5 cluster size was normalised to cell number by counting the number of DAPI positive cells. MiaPaCa2 cells were also stained for DR4 (1:100; Cell Signalling Technology, cat. 42533) as described above. DR4 mean fluorescence intensity per cell was quantified using ImageJ v1.52a (National Institutes of Health, Bethesda, Maryland, USA).

### Live imaging of GFP-tagged death receptor 5 (DR5) in pancreatic ductal adenocarcinoma cells

For lentiviral expression of mGFP-tagged DR5, MiaPaCa2 cells were seeded at 10,000 cells/well in a 24-well plate. 24 hours later, medium was replaced with 50 μL pre-made lentiviral particles (TNFRSF10B, Origene, Cat. RC201588L4V) in 450 μL complete culture medium, supplemented with 8 μg/mL polybrene. 24 hours later, medium was replaced with complete culture medium. 72 hours post-transfection, transduced cells were cultured in complete culture medium containing 1 μg/mL puromycin for 2 weeks to select transduced cells. GFP-positive cells were sorted on a BD FACS Aria II. Sorted cells were transfected with control non-silencing or βIII-tubulin siRNA and then re-seeded into 8-well glass bottom chamber slides (#1.5, 0.170 mm thickness, Ibidi, cat. 80826). At 48 hours post-transfection, cells were imaged on a Zeiss Elyra 7 Lattice SIM^2^ microscope using a 63x/1.46 objective with total internal reflection (TIRF) imaging to visualise the thin layer (<150 nm) containing the cell membrane (TIRF mirror angle set to 66.25°). 488 nm laser line was used to excite GFP and BP 495-550 emission filter for collection. To observe whether DR5 clustering precedes cell death, cells were imaged for 3.5 hours every 40 seconds, while being maintained in a humidified chamber at 37 °C and 5% CO2. To analyse the dynamics and diffusion of DR5-GFP, cells were imaged with 50 ms exposure on PCO edge sCMOS (pixel size 0.097 μm), using the same TIRF settings, and 1000 frames were acquired per field of view. DR5-GFP diffusion was quantified using k-space image correlation spectroscopy (kICS), as previously described [59-61]. Briefly, image time series were loaded in a custom-built script in MATLAB (MathWorks, Natick, MA). First, images were corrected for potential spatial drift using *imregtform* function in MATLAB. Next, pixels outside of the cellular areas were padded with the mean intensity of pixels within cells, as this ensures that they do not contribute to the kICS correlation function. Prior to application of kICS correlation function (CF) calculation, the padded image series were spatially Hann windowed, to remove the high spatial frequencies ‘leakage’ in kICS CF, coming from the image cell edges. kICS CF was calculated as previously described [59-61] and azimuthally averaged at every temporal lag. The resulting kICS Cf was fitted with two dynamic components as detailed previously [59-61] and temporally (tau lag) varying amplitudes and decays of dynamic components assessed to extract the diffusion coefficients at micro and macro spatial scales. The effective diffusion on largest spatial scale is extracted from long tau slope of macro component and represents the diffusion of DR5-GFP over whole cell surface area.

### Total internal reflection (TIRF) microscopy of microtubules on the cell membrane of pancreatic ductal adenocarcinoma cells

To observe the distribution of microtubules on the cell membrane of PDAC cells, MiaPaCa2, PANC1, and TKCC10 cells were transfected with control non-silencing or βIII-tubulin siRNA and then re-seeded into 8-well chamber slides (#1.5, 0.170 mm thickness, Ibidi, cat. 80826). 48 hours post-transfection, cells were fixed in 4% paraformaldehyde for 10 minutes. Immunofluorescence staining was performed as previously described [16], using primary antibodies for mouse anti-human βIII-tubulin (1:500; Biolegend, cat. 801202), rabbit anti-human βIII-tubulin (1:500; Covance, cat. MRB-435P) and α-tubulin (1:500; Sigma-Aldrich, cat. T9026). Primary antibodies were tagged using either goat anti-mouse AF647 (1:500; abcam, cat. ab150115), goat anti-rabbit AF647 (1:500; abcam, cat. Ab150079), or goat anti-mouse AF488 (1:500; ThermoFisher, cat. A11001) secondary antibodies. AF647 was excited by a 642 nm laser line and collected through LP655 filter, and AF488 was excited by a 488 nm laser line and collected through BP495-575 filter on Andor iXon 897 EMCCD camera with 33 ms exposure with 16 frames averaging and pixel size of 0.1 μm. Cells were kept in 1x PBS and imaged on a Zeiss Elyra PALM/SIM Super-resolution Microscope using a 100x/1.46 objective and a TIRF mirror angle of 62.55°.

### Cancer-associated fibroblast and pancreatic ductal adenocarcinoma cell co-culture

PDAC cells (MiaPaCa2, TKCC5 and TKCC10) with stable expression of GFP were transfected with control non-silencing or βIII-tubulin siRNA. After 24 hours, transfected PDAC cells were co-seeded into 12 well plates with patient derived PDAC CAFs at a 3:1 ratio of CAF:PDAC cells (54,000 CAFs + 18,000 PDAC cells). This ratio of CAF:PDAC cells was chosen to be consistent with previous co-culture work by our lab [55]. Cells were allowed to adhere for 24 hours, then placed in an IncuCyte® S3 Live-Cell Analysis System (Essen BioScience) with phase contrast and green fluorescence images taken every 30 minutes for a further 48 hours. The IncuCyte® cell-by-cell analysis software was used to quantify the number of GFP-positive cancer cells.

To measure apoptosis, GFP-labelled PDAC cells were transfected with non-silencing siRNA or βIII-tubulin siRNA and then co-seeded into 6 well plates with patient-derived CAF cells at a 3:1 ratio of CAF:PDAC cells (150,000 CAFs + 50,000 PDAC cells). At 72 hours post transfection, cells were harvested for detection of apoptosis with Annexin V/DAPI staining on a FortessaSORP flow cytometer. Cells were gated for GFP-positive cells to quantify apoptosis in tumour cells only (gating strategy shown in **Supplementary Figure 16B**).

To observe caspase 8 cleavage in a coculture setting, PDAC cells were transfected with non-silencing siRNA or βIII-tubulin siRNA and then co-seeded into 8-well chamber slides with patient-derived CAF cells at a 3:1 ratio of CAF:PDAC cells (15,000 CAFs + 5,000 PDAC cells per well). At 48 hours post-transfection, cells were treated with or without 2 μg/mL TNFα neutralising antibody (R&D, cat. MAB210). Cells were fixed in 4% paraformaldehyde at 72 hours post transfection. Immunofluorescence staining was performed as described above with antibodies for cleaved caspase 8 (rabbit, 1:100; Cell Signalling Technology, cat. #9496), AlexaFluor-488 conjugated cytokeratin (1:100, BioLegend, cat. 628608), and secondary antibody goat anti-rabbit AF647 (1:500, abcam, cat. Ab150079). Stained cells were mounted with Prolong Gold anti-fade mounting media with DAPI to visualise cell nuclei. Images were taken on a Zeiss LSM800 confocal microscope using a 40x/1.3 NA objective. AF647 was excited with a 640 nm laser and captured using detection wavelength 450-700 nm. AF488 was excited with a 488 nm laser and captured using detection wavelength 490-620 nm. DAPI was excited with a 405 nm laser and captured using detection wavelength 400-605 nm. For quantification of caspase 8 cleavage, QuPath v0.3.2 [58]was used to count the percentage of cytokeratin positive PDAC cells that were positive for cleaved caspase 8.

### Patient derived pancreatic ductal adenocarcinoma whole-tissue explant culture

Patient surgical PDAC tumour samples were obtained from Prince of Wales Hospital or Prince of Wales Private Hospital (Randwick, New South Wales, Australia). All patients provided written informed consent through the Health Precincts Biobank, all work was approved by UNSW human ethics (HC180973), and all experiments were performed in accordance with the relevant regulations. Patient-derived explants (1-2 mm diameter) were manually cut from fresh surgical samples and cultured for 12 days with daily medium changes, as described previously [27]. Star 3 nanoparticles (60 μg) were complexed to control or βIII-tubulin siRNA (20 μg) and added to the medium reservoir as described previously [27, 55] on days 0, 3, 6, 9. Star 3 + siRNA dosing was calculated to match the blood concentration of Star 3 + siRNA used *in vivo* (2.4 mg/kg) assuming an average 20 g mouse has a blood volume of 1.2 mL. Star 3 + siRNA treatments were performed on day 0 and repeated every 72 hours to ensure efficient protein knockdown throughout the 12-day culture. On days 5 and 10, tumour explants were treated with or without 500 ng/mL human recombinant TRAIL. TRAIL concentration was 40-fold lower than the maximum serum concentration of dulanermin (clinical TRAIL drug) measured in clinical pharmacokinetic studies at a dose of 4 mg/kg [62]. Depending on the size of the surgical sample received, 2 to 4 explants from different regions of the sample were cultured per treatment group. Patient-derived explants were treated with 10 μM BrdU substrate (BD Biosciences, catalogue no. 550891) for 24 hours prior to fixation (day 12), as described previously to assess proliferation [27]. H&E-stained sections of tumour explants are shown in **Supplementary Figure 20**.

### Immunohistochemistry of human pancreatic ductal adenocarcinoma tissue sections

Immunohistochemistry on paraformaldehyde-fixed and paraffin-embedded PDAC tumour explant sections was performed as previously described [27]. Tissue sections were stained for cleaved caspase 8 (Cell Signalling, Cat. 9748S, 1:100 dilution overnight at 4 °C), bromodeoxyuridine (BrdU; explants treated with BrdU substrate for 24 hours prior to fixation as described in Section 6.3.1) (DAKO, Cat. M0744; 1:50 overnight at 4 °C), cytokeratin (DAKO, Cat. M3515; 1:100 at 4 °C), ki67 (ThermoFisher, Cat. RM-9106; 1:50 overnight at 4 °C), βIII-tubulin (Biolegend, Cat. 801202; 1:100 overnight at 4 °C) and α-smooth muscle actin (αSMA) (Sigma, Cat. A5228; 1:1000 1 hour room temperature). Isotype control antibodies (mouse IgG2A, mouse IgG1A, and rabbit IgG) were used as negative controls (representative images shown in **Supplementary Figure 19**). All stained slides were scanned on Vectra Polaris (PerkinElmer) (40x/0.75 NA objective) or VS200 (Olympus) (40x objective) slide scanners. Quantification of staining was performed using the positive cell detection function on QuPath.

### Correlation of βIII-tubulin expression in human pancreatic ductal adenocarcinoma samples with overall survival

Staining for βIII-tubulin in human PDAC tissue microarrays (TMA) was performed in the same way as described in Section 9.20. Formalin-fixed and paraffin-embedded PDAC TMAs were obtained through the Australian Pancreatic Cancer Genome Initiative as part of the International Cancer Genome Consortium Cohort. All work was approved by UNSW human ethics (HC180973). All patients provided written informed consent, and patient demographics are included in **Supplementary Table 1**. Stained TMA slides were scanned on an AperioXT (Leica Biosystems) slide scanner. Staining intensity and percentage of stained cells in tumour and stromal compartments were scored by 3 independent scorers on a 4-point scale (0,1,2,3). The following index was then used to calculate the overall score for tumour and stromal compartments, per scorer, based on the percentage of cells stained: an overall score of 0 if 100% of cells were negative; an overall score of 1 if greater than 50% of cells were scored as a 0 or 1; an overall score of 2 if there was a 50%:50% ratio of cells with a score of 0 or 1 to a score of 2 or 3, or 50% of cells were scored as a 2 or 3; or an overall score of 3 if more than 50% of cells were scored as a 2 or 3. This scoring system that includes staining intensity and percent of cells stained has been described previously [18]. A consensus score was then determined for each core. For each set of 3 cores per patient, the highest tumour and stroma scores were used. Overall scores of 0-1 were classified as low, and overall scores of 2-3 were classified as high. Scores were then correlated with overall survival using a Kaplan Meier Survival Curve (see statistical analyses). Patients deceased due to other causes/still alive were censored. Non-PDAC tumours and patients that lacked 3 full cores were excluded (patient cohort details summarised in **Supplementary Table 1**).

### Immunofluorescence for βIII-tubulin and α-smooth muscle actin colocalisation

Human surgical PDAC tissue was obtained fresh Prince of Wales Hospitals through the Health Precincts Biobank as described above. Fresh PDAC tissue was fixed in 4% paraformaldehyde for 24 hours and paraffin embedded. Antigen retrieval on tissue sections was performed as previously described [27]. Tissue sections were stained with monoclonal mouse anti-βIII-tubulin primary antibody (Biolegend, Cat. 801202; 1:100 overnight at 4 °C) and goat anti-mouse AF647 (abcam, cat. ab150115; 1:500 for 1 hour at room temperature) secondary antibody was used. Tissue sections were then subsequently stained with Cy3-conjugated αSMA antibody (Sigma-Aldrich, cat. C6198; 1:200 for 1 hour at room temperature) then mounted with Prolong Gold anti-fade mounting media with DAPI to visualise cell nuclei. Staining was imaged on a Zeiss LSM900 confocal microscope using a 40x/1.3 NA objective). AF647 was excited with a 640 nm laser and captured using detection wavelength 645-700 nm. Cy3 was excited with a 561 nm laser and captured using detection wavelength 535-617 nm. DAPI was excited with a 405 nm laser and captured using detection wavelength 400-605 nm.

### Validation of βIII-tubulin antibody

The βIII-tubulin antibody used in this study had been previously shown to be specific to βIII-tubulin [16]. Our lab previously demonstrated using western blot that knockdown of βII-tubulin did not affect βIII-tubulin protein levels, indicating that the βIII-tubulin antibody does not bind to βII-tubulin, which is one of the major isotypes expressed in cells. We also showed positive staining of βIII-tubulin in human brain tissue, and negative staining in normal pancreas tissue, consistent with expression in the human protein atlas (**Supplementary Figure 21**). Specificity was also shown by demonstrating protein knockdown via western blot using both siRNA and shRNA (**Supplementary Figure 1** and **Supplementary Figure 2E**).

### Statistical analysis

Statistical analyses were performed using GraphPad Prism 9 (GraphPad Software). All data were presented as mean ± standard error of mean (SEM). For experiments with n≥3 independent replicates, two-tailed student t-test or one-way ANOVA followed by non-parametric Bonferroni’s multiple comparisons test were performed to calculate statistical significance, which was assigned to p<0.05. Comparisons of univariate time to event (survival) were performed using the log-rank test and hazard ratios calculated from the Cox proportional hazards (PH) model. Multivariate associations between variables and time to event were contained from PH regression and survival curves calculated using the method of Kaplan-Meier (KM). Where tumour and stroma scores correlated with outcome, baseline variables associated with predicting scores were examined by multivariate logistic regression. Tumour grade was excluded from multivariate analyses as it did not correlate with overall survival in our cohort due to the low percentage of grade 3-4 tumours (13% of cohort). Survival analyses were performed using Analysis of Censored and Correlated Data (ACCoRD; RRID:SCR_009015) V6.4 Boffin. A p-value <0.05 was considered statistically significant.

## Acknowledgements

Biospecimens and data used for tumour explants were obtained from the Health Precincts Biobank, UNSW Biorepository, UNSW Sydney, Australia. We sincerely thank the patients who consented to donate their tumour samples for research. We would like to thank Dr. Carmel Quinn and Dr. Anusha Hettiaratchi of the Health Precincts Biobank for their support in managing clinical samples and patient consent. We would also like to acknowledge our community consumers Gino Iori, Michelle Daly, and Claire Harvey for their invaluable input on the project and grant applications. Biospecimens and clinical data for prognostic studies were provided by the Avner APGI Bioresource (**www.pancreaticcancer.net.au**), which is supported by PanKind, The Australian Pancreatic Cancer Foundation (**www.pankind.org.au**). We acknowledge the Flow Cytometry, Katharina Gaus Light Microscopy Facility, and Biological Resource Imaging Facility within the Mark Wainwright Analytical Centre at UNSW Sydney for their technical support.

This work was made possible by the following major funding sources: NHMRC Ideas Grant (Phillips, Sharbeen, APP2002707), NHMRC project grant (Phillips, McCarroll, Goldstein, APP1144108), Tour de Cure Senior Research Grant (Phillips, McCarroll, Goldstein, RSP-235-2020), Tour de Cure Pioneering Research Grant (Sharbeen, Phillips, Goldstein, RSP-255-2020), Cancer Institute NSW Translational Program Grant (Phillips, Goldstein, 2020/TPG2100), Tour de Cure PhD Support Scholarship (Kokkinos, Phillips, Goldstein, RSP-011-18/19), and Cancer Research UK Institute Award (Morton, A29996). We acknowledge the generous philanthropic support from Mr Paul Dainty, Dr Marjorie O’Neil, Dr Keri Spooner. The following sources supported author contributions and research: Australian Government Research Training Program Scholarship and UNSW Sydney Scientia PhD Scholarship (Kokkinos), University Postgraduate Award (Pitiyarachchi, UNSW Sydney, O.P.), Cancer-Institute NSW CDF (Sharbeen, CDF181166), Maridulu Budyari Gumal Sydney Partnership for Health, Education, Research and Enterprise [SPHERE] Cancer Clinical Academic Group Senior Research Fellowship (Funded by Cancer Institute NSW Translational Cancer Research Capacity Building Grant, 2021/CBG0003, Sharbeen), Avner Pancreatic Cancer Foundation Innovation Grant (Phillips, McCarroll, Goldstein, Davis and Sharbeen, APCF0050618), Cancer-Institute NSW ECF (Sharbeen, 13/ECF/1–to08), Olivia Lambert Foundation (McCarroll), and Cancer Australia (Phillips, McCarroll and Goldstein, APP1126736).

## Author contributions

Conceptualisation, J.K, G.S, D.G, J.A.M, P.A.P; Methodology, J.K, G.S, E.P, D.G, R.M.W, J.A.M, P.A.P; Investigation, J.K, G.S, E.P, D.G, J.A.M, P.A.P, R.M.C.I, J.Y, C.B, M.G, V.G, M.E.D, G.S, O.S.M.A, C.K, E.G, A.M; Resources, K.S.H, M.P, M.E, A.J, A.J.G, APGI; Writing – Original Draft, J.K, G.S, J.A.M, P.A.P; Writing – Review & Editing, J.K, G.S, J.A.M, P.A.P, D.G, R.M.C.I, K.S.H, V.G, O.P, M.E.D, M.P, M.E, J.P.M, M.K; Supervision, J.A.M, P.A.P, Funding Acquisition, J.K, G.S, D.G, J.A.M, P.A.P.

## Supplementary information

**Supplementary Table 1. Australian Pancreatic Cancer Genome Initiative (APGI) International Cancer Genome Cohort (ICGC) patient characteristics for βIII-tubulin survival analyses**.

Human PDAC tissue microarrays were obtained through the APGI. Patient cohort characteristics are described above. TNM staging refers to tumour size (T), lymph node involvement (N), and metastasis (M).

**Supplementary Table 2. βIII-tubulin multivariate survival analysis parameters**.

Comparisons of univariate time to event (survival) were performed using the log-rank test and hazard ratios (HR) and confidence intervals (CI) calculated from the Cox proportional hazards (PH) model. Multivariate associations between variables and time to event were contained from PH regression and survival curves calculated using the method of Kaplan-Meier (KM). Where tumour and stroma scores correlated with outcome, baseline variables associated with predicting scores were examined by multivariate logistic regression. A p-value ≤0.05 was considered statistically significant.

**Supplementary Figure 1. Confirmation of βIII-tubulin knockdown in PDAC and NSCLC cells. (A-B)** βIII-tubulin (βIII-Tub) mRNA expression in **(A)** MiaPaCa2 (n=3 independent experiments) and **(B)** PANC1 cells (n=3 independent experiments) was significantly reduced 72 hours post-transfection with βIII-tubulin siRNA, compared to non-silencing controls, as assessed by quantitative real-time PCR. Samples were standardised to 18S RNA. **(C-H)** Western blot analysis (representative blots shown) with densitometry graphs below showing reduced βIII-tubulin protein levels, 72 hours following transfection with βIII-tubulin siRNA in MiaPaCa2 (n=4) **(C)**, PANC1 (n=3) **(D)**, TKCC5 (n=3) **(E)**, TKCC10 (n=3) **(F)**, HPAF-II (n=4) **(G)**, and H460 (n=4) **(H)** cells. Densitometry analysis (graphs) was performed by measuring βIII-tubulin protein expression normalised to GAPDH expression. Bars represent mean of n≥3 independent experiments (individual data points shown from independent experiments) ± standard error of mean. Asterisks indicate significance as assessed by two-tailed paired t-tests comparing mean of βIII-Tub siRNA to mean of ns-siRNA (*p≤0.05, **p≤0.01, ***p≤0.001).

**Supplementary Figure 2: βIII-Tubulin silencing in PDAC cells increased sensitivity to TRAIL. (A)** A western blot was performed to compare the relative expression of TRAIL receptors DR4 and DR5 in PDAC and NSCLC cell lines used in this study. Protein was loaded onto two separate gels to allow imaging of both DR4 and DR5. Membranes were then re-probed for GAPDH as a loading control. **(B-D)** Knockdown of βIII-tubulin combined with TRAIL decreased cell viability in MiaPaCa2 (n=4) **(B)**, PANC1 (n=3) **(C)**, and TKCC10 (n=3) **(D)** cells. Cells were treated with TRAIL 48 hours post-transfection and then cell viability measured using a trypan blue exclusion assay 96 hours post transfection. **(E)** MiaPaCa2 cells with stable expression of βIII-tubulin (βIII-tub) shRNA demonstrated potent knockdown of βIII-tubulin protein (western blot). Knockdown of βIII-tubulin using an independent single-sequence siRNA (Qiagen) demonstrated potent protein silencing comparable to knockdown with SMARTpool siRNA (Dharmacon). **(F)** Knockdown of βIII-tubulin using single sequence siRNA in MiaPaCa2 cells (n=3) increased apoptosis in the presence of TRAIL (10 ng/mL). Apoptosis was assessed by flow cytometry for Annexin V/DAPI 72 hours post transfection. **(G)** PDAC cells were seeded at low density post-transfection with ns-siRNA or βIII-tubulin siRNA and were cultured with TRAIL for 72 hours (n=4). Colonies were stained with crystal violet (representatives shown) and counted 9 days post-seeding in MiaPaCa2 cells and 14 days post-seeding in HPAF-II cells. **(H)** βIII-tubulin knockdown combined with TRAIL in NSCLC H460 cells reduced cell proliferation to a greater extent than either treatment alone, measured on the IncuCyte S3 Live Cell Analysis system (n=3). Cells were treated with TRAIL (50 ng/mL) 24 hours post-transfection, and images taken every hour. Bars represent mean of n≥3 independent experiments (individual data points shown from independent experiments) ± standard error of mean. Asterisks indicate significance as assessed by One-Way ANOVA (*p≤0.05, **p≤0.01, ***p≤0.001, ****p≤0.0001).

**Supplementary Figure 3. βIII-Tubulin silencing in patient-derived PDAC CAF cells had no effect on TRAIL sensitivity. (A)** A western blot was performed to compare the relative expression of βIII-tubulin in 5 patient derived CAF lines with 4 commercial PDAC cell lines. Results show comparable expression of βIII-tubulin in CAF and PDAC cells. GAPDH was used as a loading control. **(B)** PDAC CAFs from n=5 patients were treated with TRAIL 48 hours post-transfection with ns-siRNA or βIII-tubulin siRNA. Apoptosis was measured via flow cytometry for annexin V/DAPI at 72 hours post-transfection. **(C)** CAF cell proliferation was reduced with βIII-tubulin knockdown, but there was no further reduction in cell proliferation with the addition of TRAIL. Cells were treated with TRAIL (100 ng/mL) 24 hours post-transfection, and images taken every hour on the IncuCyte S3 (n=4). **(D)** Western blot in MiaPaCa2 cells and CAFs from n=4 PDAC patients revealed varying expression levels of DR4 and DR5. Protein was loaded onto two separate gels to allow imaging of both DR4 and DR5. Membranes were then re-probed for GAPDH as a loading control. Bars represent mean of n≥3 independent experiments (individual data points shown from independent experiments) ± standard error of mean. Asterisks indicate significance as assessed by One-Way ANOVA (*p≤0.05, n.s.; non-significant).

**Supplementary Figure 4: βIII-Tubulin silencing in MiaPaCa2 cells triggered DR5 clustering. (A-B)** Immunofluorescence staining for DR5 and α-tubulin revealed that βIII-tubulin (βIII-Tub) knockdown triggered the formation of large membrane clusters of DR5 (red arrows) in MiaPaCa2 (representative of n=4 independent experiments) **(A)** and PANC1 (representative of n=4 independent experiments) **(B)** cells. Cells were fixed and stained 72 hours post transfection. MiaPaCa2 merged immunofluorescence images are also shown in **Figure 5A** but are shown here as well with each individual fluorescence channel as well. **(C)** βII-tubulin knockdown in MiaPaCa2 cells did not trigger DR5 clustering compared to βIII-tubulin knockdown. Cells were fixed and stained 72 hours post transfection. All scale bars represent 20 μm.

**Supplementary Figure 5: βIII-Tubulin regulates DR5 dynamics in PDAC cells. (A)** Quantification of multimeric DR5 clusters in MiaPaCa2 cells with βIII-tubulin knockdown treated with or without TRAIL (n=5). Quantification was performed on iBright analysis software by quantifying multimeric DR5 protein bands between 50-250 kDa and normalising to GAPDH expression. Bars represent mean of n=5 independent experiments (individual data points shown from independent experiments) ± standard error of mean. Asterisks indicate significance as assessed by One-Way ANOVA (*p≤0.05, **p≤0.01, n.s.; non-significant). **(B)** Western blot for DR4 was performed under non-reducing conditions using protein extracted from MiaPaCa2 cells at 72 hours post transfection. βIII-tubulin silencing did not induce high-molecular weight multimeric clusters of DR4. Membranes were re-probed for GAPDH. **(C)** Immunofluorescence for DR4 in MiaPaCa2 cells showed that βIII-tubulin silencing or TRAIL treatment did not trigger formation of DR4 clusters. βIII-tubulin silencing led to a reduction in DR4 mean fluorescence intensity per cell (n=3 independent experiments). Mean fluorescence intensity per cell was quantified using ImageJ software. **(D)** Live imaging of GFP-tagged DR5 on the cell membrane of MiaPaCa2 cells using total internal reflection (TIRF) microscopy revealed that βIII-tubulin silencing induced DR5 membrane clustering. Cells were imaged 48 hours post transfection. Cells with βIII-tubulin knockdown that had visible DR5 membrane clustering showed characteristic features of apoptosis such as membrane blebbing and fragmentation from 180 minutes of imaging. See **Supplementary Movie 1** for video of live cell imaging. **(E)** Immunofluorescence and TIRF microscopy at 48 hours post transfection revealed a mesh-like architecture of βIII-tubulin on the cell membrane of PDAC cells that was disrupted with βIII-tubulin silencing. **(F)** α-tubulin showed a similar mesh-like structure on the cell membranes of PDAC cells but was unaffected by βIII-tubulin knockdown. All scale bars represent 20 μm.

**Supplementary Figure 6: Anti-proliferative effects of βIII-tubulin silencing when pancreatic ductal adenocarcinoma (PDAC) cells are co-cultured with cancer associated fibroblasts (CAFs)**. Cell proliferation was reduced in PDAC cells with βIII-tubulin (βIII-Tub) knockdown when co-cultured with CAFs, as measured on an IncuCyte® S3 Live-Cell Analysis System. **(A)** Representative images show GFP-labelled PDAC cells surrounded by CAFs visible via phase contrast. **(B-C)** GFP-labelled TKCC5 (n=3 independent experiments) **(B)** and TKCC10 (n=3 independent experiments) **(C)** cells were transfected with non-silencing siRNA (ns-siRNA) or βIII-tubulin siRNA and co-cultured in the presence of primary patient-derived CAFs. The integrated IncuCyte software was used to count the number of GFP-positive PDAC cells. Line represents mean ± standard error of mean. n=3 indicates 3 independent experiments using CAFs isolated from 3 different PDAC patients. Asterisks indicate significance as assessed by two-tailed paired t-test (*p≤0.05).

**Supplementary Figure 7: Star 3 nanoparticle delivery of βIII-tubulin siRNA led to a potent reduction in βIII-tubulin expression in patient-derived pancreatic ductal adenocarcinoma (PDAC) explants. (A)** Human PDAC explants from a single patient were harvested at day 6 after treatment with Star 3+control siRNA or Star 3+βIII-tubulin siRNA on day 0 and day 3, and TRAIL (500 ng/mL) treatment on day 4. Representative images show H&E staining and immunohistochemistry for βIII-tubulin. **(B-C)** Quantification of βIII-tubulin immunohistochemistry using QuPath demonstrated a reduction in the overall intensity of βIII-tubulin expression **(B)** and a reduction in the number of cells positive for βIII-tubulin **(C)** in explants treated with βIII-tubulin siRNA.

**Supplementary Figure 8: Patient derived tumour explants from 5 pancreatic ductal adenocarcinoma (PDAC) patients demonstrate high tumour expression of βIII-tubulin**. Immunohistochemistry staining for βIII-tubulin revealed high tumour cell expression of βIII-tubulin in control-siRNA treated PDAC explants.

**Supplementary Figure 9: βIII-tubulin silencing combined with TRAIL in Patient 1 decreased tumour cell number and increased extrinsic apoptosis in pancreatic ductal adenocarcinoma (PDAC) patient-derived explants**. Pancreatic ductal adenocarcinoma (PDAC) tumour explants derived from Patient 1 were treated with Star 3 + βIII-tubulin (βIII-Tub) siRNA on days 0, 3, 6, and 9, and TRAIL (500 ng/mL) on days 5 and 10, with 2 explants per treatment. **(A)** Representative immunohistochemistry staining of serial sections for cytokeratin (tumour cell marker) and cleaved caspase 8 (extrinsic apoptosis marker) at low and high magnification. **(B-C)** Quantification of whole tumour explants was performed on QuPath for immunohistochemistry staining of cytokeratin **(B)** and cleaved caspase 8 **(C)**. Results show a reduction in tumour cell number with βIII-tubulin silencing combined with TRAIL compared to control explants. Caspase 8 cleavage was also increased in all treatment groups compared to control-siRNA untreated explants. Symbols represent quantification of individual whole tumour explant sections. Bars represent mean ± standard error of mean.

**Supplementary Figure 10: βIII-tubulin silencing combined with TRAIL in Patient 2 decreased tumour cell number and decreased cell proliferation in pancreatic ductal adenocarcinoma (PDAC) patient-derived explants**. Pancreatic ductal adenocarcinoma (PDAC) tumour explants derived from Patient 2 were treated with Star 3 + βIII-tubulin (βIII-Tub) siRNA on days 0, 3, 6, and 9, and TRAIL (500 ng/mL) on days 5 and 10, with 2 explants per treatment. **(A)** Representative immunohistochemistry staining of serial sections for cytokeratin (tumour cell marker), bromodeoxyuridine (BrdU) (proliferation marker), and cleaved caspase 8 (extrinsic apoptosis marker) at low and high magnification. **(B-D)** Quantification of whole tumour explants was performed on QuPath for immunohistochemistry staining of cytokeratin **(B)**, BrdU **(C)**, and cleaved caspase 8 **(D)**. Results show a reduction in tumour cell number in tumour explants with βIII-tubulin silencing combined with TRAIL compared to control explants. Cell proliferation was also reduced in all treatment groups compared to control-siRNA untreated explants. Symbols represent quantification of individual whole tumour explant sections. Bars represent mean ± standard error of mean.

**Supplementary Figure 11: βIII-tubulin silencing combined with TRAIL in Patient 3 decreased tumour cell number and decreased cell proliferation in pancreatic ductal adenocarcinoma (PDAC) patient-derived explants**. Pancreatic ductal adenocarcinoma (PDAC) tumour explants derived from Patient 3 were treated with Star 3 + βIII-tubulin (βIII-Tub) siRNA on days 0, 3, 6, and 9, and TRAIL (500 ng/mL) on days 5 and 10, with 4 explants per treatment. **(A)** Representative immunohistochemistry staining of serial sections for cytokeratin (tumour cell marker), bromodeoxyuridine (BrdU) (proliferation marker), and cleaved caspase 8 (extrinsic apoptosis marker) at low and high magnification. Representative images are also shown in **Figure 7** and are reproduced here showing quantification of each stain from Patient 3. **(B-D)** Quantification of whole tumour explants was performed on QuPath for immunohistochemistry staining of cytokeratin **(B)**, BrdU **(C)**, and cleaved caspase 8 **(D)**. Results show a reduction in tumour cell number in tumour explants with βIII-tubulin silencing alone and TRAIL treatment alone compared to control-siRNA untreated explants with no further decrease in combination treated explants. BrdU staining was reduced in all TRAIL treated explants, and caspase 8 cleavage was increased in tumour explants with βIII-tubulin silencing alone compared to control-siRNA untreated explants. Symbols represent quantification of individual whole tumour explant sections. Bars represent mean ± standard error of mean.

**Supplementary Figure 12: βIII-tubulin silencing combined with TRAIL in Patient 4 decreased tumour cell number in pancreatic ductal adenocarcinoma (PDAC) patient-derived explants**. Pancreatic ductal adenocarcinoma (PDAC) tumour explants derived from Patient 4 were treated with Star 3 + βIII-tubulin (βIII-Tub) siRNA on days 0, 3, 6, and 9, and TRAIL (500 ng/mL) on days 5 and 10, with 2 explants per treatment. **(A)** Representative immunohistochemistry staining of serial sections for cytokeratin (tumour cell marker), bromodeoxyuridine (BrdU) (proliferation marker), and cleaved caspase 8 (extrinsic apoptosis marker) at low and high magnification. **(B-D)** Quantification of whole tumour explants was performed on QuPath for immunohistochemistry staining of cytokeratin **(B)**, BrdU **(C)**, and cleaved caspase 8 **(D)**. Results show a reduction in tumour cell number in tumour explants with βIII-tubulin silencing combined with TRAIL compared to control-siRNA untreated explants. Symbols represent quantification of individual whole tumour explant sections. Bars represent mean ± standard error of mean.

**Supplementary Figure 13: βIII-tubulin silencing combined with TRAIL in Patient 5 increased extrinsic apoptosis in pancreatic ductal adenocarcinoma (PDAC) patient-derived explants**. Pancreatic ductal adenocarcinoma (PDAC) tumour explants derived from Patient 5 were treated with Star 3 + βIII-tubulin (βIII-Tub) siRNA on days 0, 3, 6, and 9, and TRAIL (500 ng/mL) on days 5 and 10, with 3 explants per treatment. **(A)** Representative immunohistochemistry staining of serial sections for cytokeratin (tumour cell marker), bromodeoxyuridine (BrdU) (proliferation marker), and cleaved caspase 8 (extrinsic apoptosis marker) at low and high magnification. **(B-D)** Quantification of whole tumour explants was performed on QuPath for immunohistochemistry staining of cytokeratin **(B)**, BrdU **(C)**, and cleaved caspase 8 **(D)**. Results show an increase in caspase 8 cleavage in tumour explants with βIII-tubulin silencing alone and TRAIL treatment alone compared to control-siRNA untreated explants. There was no further increase in caspase 8 cleavage with combination treatment compared to individual treatments. Symbols represent quantification of individual whole tumour explant sections. Bars represent mean ± standard error of mean.

**Supplementary Figure 14: βIII-tubulin silencing in Patient 6 decreased cell proliferation and increased extrinsic apoptosis in tumour explants derived from a stomach metastasis of a patient with pancreatic ductal adenocarcinoma (PDAC)**. Metastatic PDAC tumour explants derived from Patient 6 were treated with Star 3 + βIII-tubulin (βIII-Tub) siRNA on days 0, 3, 6, and 9, with 3 explants per treatment. **(A-C)** Representative images and quantification of immunohistochemistry staining of serial sections for cytokeratin (tumour cell marker) **(A)**, bromodeoxyuridine (BrdU) (proliferation marker) **(B)**, and cleaved caspase 8 (extrinsic apoptosis marker) **(C)** at low and high magnification. Quantification of whole tumour explants was performed on QuPath Results show a decrease in cell proliferation and an increase in caspase 8 cleavage in tumour explants with βIII-tubulin silencing compared to control-siRNA explants. Symbols represent quantification of individual whole tumour explant sections. Bars represent mean ± standard error of mean.

**Supplementary Figure 15: βIII-tubulin silencing and TRAIL treatment had patient-specific effects on cancer-associated fibroblast (CAF) cell number in pancreatic ductal adenocarcinoma (PDAC) tumour explants. (A)** Representative immunohistochemistry staining of CAF marker α-smooth muscle actin (αSMA) in PDAC tumour explants from patients 1-5. **(B)** Quantification of whole tumour explants was performed on QuPath for immunohistochemistry staining of αSMA and data combined from n=5 pateints taking the average quantification of 2-4 explants from each patient. Bars represent mean from n=5 patients (individual data points shown from each patient) ± standard error of mean. Asterisks indicate significance as assessed by One-Way ANOVA (n.s.; non-significant).

**Supplementary Figure 16: Gating strategy for apoptosis assays**. Apoptosis was measured via flow cytometry for Annexin V/DAPI using the FortessaSORP flow cytometer. **(A-B)** Panels show representative flow cytometry plots with gating strategies used for standard apoptosis assays **(A)** or co-culture apoptosis assays where apoptosis was measured in GFP positive PDAC cells **(B)**. Total apoptosis was assigned the sum of Q1+Q2+Q3 quadrants of Annexin V vs DAPI plots.

**Supplementary Figure 17: Analysis of cell proliferation on IncuCyte® S3 using confluence metrics**. Real-time cell proliferation was measured using IncuCyte® S3 Live Cell Analysis System. **(A)** Representative image shows phase contrast image of MiaPaCa2 cells. **(B)** Integrated confluence metrics on the IncuCyte® software was used to calculate the percent confluence of each sample at 1-hour intervals. Representative image shows confluence mask (yellow) applied by IncuCyte® software to calculate the percent confluence. At least 9 fields of view were captured from each well of a 12-well plate, and 16 fields of view from each well of a 6-well plate.

**Supplementary Figure 18. Validation of caspase inhibitors. (A)** Caspase 9 (casp9) inhibitor (Z-LEHD-FMK) potently blocks paclitaxel induced caspase 9 activity. MiaPaCa2 cells were pre-incubated with 20 μM caspase 9 inhibitor for 1 hour then treated with 10 nM paclitaxel for 24 hours (n=1 independent experiment). Caspase-9 activity was measured using CaspaseeGlo-9 luminescent assay and normalised to cell number using cell counting kit-8 (CCK-8) assay. **(B)** Caspase 8 (casp8) inhibitor (Q-IETD-OPh) blocks TRAIL induced apoptosis. MiaPaCa2 cells were pre-incubated with 20 μM caspase 8 inhibitor for 1 hour then treated with 10 ng/mL TRAIL for 24 hours (n=1). Apoptosis was measured using flow cytometry for Annexin V and DAPI.

**Supplementary Figure 19. Immunohistochemistry isotype controls. Representative images of each isotype control antibody used for immunohistochemistry**. Mouse IgG1A was used for cytokeratin, cleaved caspase 8 (mouse), and BrdU. Mouse IgG2A was used for α-Smooth Muscle Actin and βIII-tubulin. Rabbit IgG was used for ki67 and cleaved caspase 8 (rabbit).

**Supplementary Figure 20. H&E staining of pancreatic ductal adenocarcinoma (PDAC) tumour explants from patients 1-6 used for βIII-tubulin experiments**. Representative images show 1 explant from each treatment group per patient.

**Supplementary Figure 21. Validation of βIII-tubulin antibody**. Positive staining of βIII-tubulin (βIII-Tub) in human brain tissue, and negative staining in normal pancreas tissue, consistent with expression in the human protein atlas.

**Supplementary Movie 1. βIII-tubulin silencing triggered death receptor 5 (DR5) clustering in MiaPaCa2 cells prior to induction of cell death**. Live imaging of GFP-tagged DR5 on the cell membrane of MiaPaCa2 cells using total internal reflection (TIRF) microscopy revealed that βIII-tubulin silencing induced DR5 membrane clustering. Cells were imaged 48 hours post transfection. Cells with βIII-tubulin knockdown that had visible DR5 membrane clustering showed characteristic features of apoptosis such as membrane blebbing and fragmentation from 180 minutes of imaging. Cells were imaged for 3.5 hours every 40 seconds, while being maintained in a humidified chamber at 37 °C and 5% CO2.

**Supplementary Movie 2. βIII-tubulin silencing triggered death receptor 5 (DR5) clustering in MiaPaCa2 cells prior to induction of cell death**. Live imaging of GFP-tagged DR5 in MiaPaCa2 cells using total internal reflection (TIRF) microscopy revealed that βIII-tubulin silencing induced highly dynamic clusters of DR5 at the cell membrane. Cells were imaged 48 hours post transfection with 50 ms exposure on PCO edge sCMOS (pixel size 0.097 μm), using TIRF settings, and 1000 frames were acquired per field of view.

