## Supplementary Tables and Figures for "βIII-Tubulin is a Brake on Extrinsic Cell-Death in Pancreatic Cancer"

| Age at diagnosis | Number of patients |
|---|---|
| ≥50 | 146 |
| <50 | 9 |
| **Gender** | |
| Male | 78 |
| Female | 77 |
| **Ethnicity** | |
| Asian | 10 |
| Asian, White/Caucasian | 1 |
| Black/African | 2 |
| Pacific Islander | 1 |
| White/Caucasian | 141 |
| **Smoker** | |
| Ever | 80 |
| Never | 72 |
| Not reported | 3 |
| **Alcohol consumption** | |
| Ever | 88 |
| Never | 63 |
| Not reported | 4 |
| **Margin Status** | |
| R0 | 108 |
| R1 | 42 |
| R2 | 3 |
| RX | 2 |
| **Macroscopic tumour location** | |
| Ampulla | 1 |
| Body | 10 |
| Head | 116 |
| Head (Uncinate) | 8 |
| Tail | 17 |
| Not reported | 3 |

| Overall Stage | Number of patients |
|---|---|
| IA | 2 |
| IB | 4 |
| IIA | 31 |
| IIB | 109 |
| III | 1 |
| IV | 6 |
| Not reported | 2 |
| **TNM Staging** | |
| T1 | 2 |
| T2 | 7 |
| T3 | 142 |
| T4 | 1 |
| TX | 3 |
| N0 | 37 |
| N1 | 78 |
| N1a | 4 |
| N1b | 32 |
| NX | 4 |
| M0 | 3 |
| M1 | 6 |
| MX | 144 |
| **Perineural invasion** | |
| Yes | 127 |
| No | 22 |
| Not reported | 6 |
| **Vascular invasion** | |
| Yes | 94 |
| No | 55 |
| Not reported | 6 |
| **Recurrence at liver** | |
| Yes | 50 |
| No | 58 |
| No recurrence | 47 |

**Supplementary Table 1. Australian Pancreatic Cancer Genome Initiative (APGI) International Cancer Genome Cohort (ICGC) patient characteristics for βIII-tubulin survival analyses.**

| Univariate analysis of parameters used in multivariate analysis | | | | |
|---|---|---|---|---|
| **Parameter** | **HR** | **95%** | **CI** | **p-value** |
| βIII-Tubulin Tumour score | 1.526 | 1.052 | 2.206 | 0.026 |
| βIII-Tubulin Stroma score | 1.766 | 1.227 | 2.541 | 0.002 |
| Gender | 0.651 | 0.454 | 0.934 | 0.020 |
| Age at Diagnosis (years) | 0.975 | 0.453 | 2.102 | 0.949 |
| Smoker | 1.292 | 0.896 | 1.863 | 0.170 |
| Alcohol consumption | 1.129 | 0.779 | 1.636 | 0.520 |
| Margin Status | 1.624 | 1.106 | 2.385 | 0.013 |
| Lymph Nodes Involved | 1.453 | 0.928 | 2.274 | 0.102 |
| Perineural Invasion | 1.727 | 0.986 | 3.025 | 0.056 |
| Vascular Invasion | 1.779 | 1.193 | 2.655 | 0.005 |
| Overall Stage AJCC | 1.691 | 1.072 | 2.665 | 0.024 |
| Macroscopic Tumour Location | 1.103 | 0.598 | 2.035 | 0.754 |
| **Multivariate results (best subsets)** | **HR** | **95%** | **CI** | **p-value** |
| βIII-Tubulin Stroma score | 2.163 | 1.208 | 3.874 | 0.009 |
| Margin Status | 2.043 | 1.345 | 3.103 | 0.001 |

**Supplementary Table 2. βIII-tubulin multivariate survival analysis parameters.**

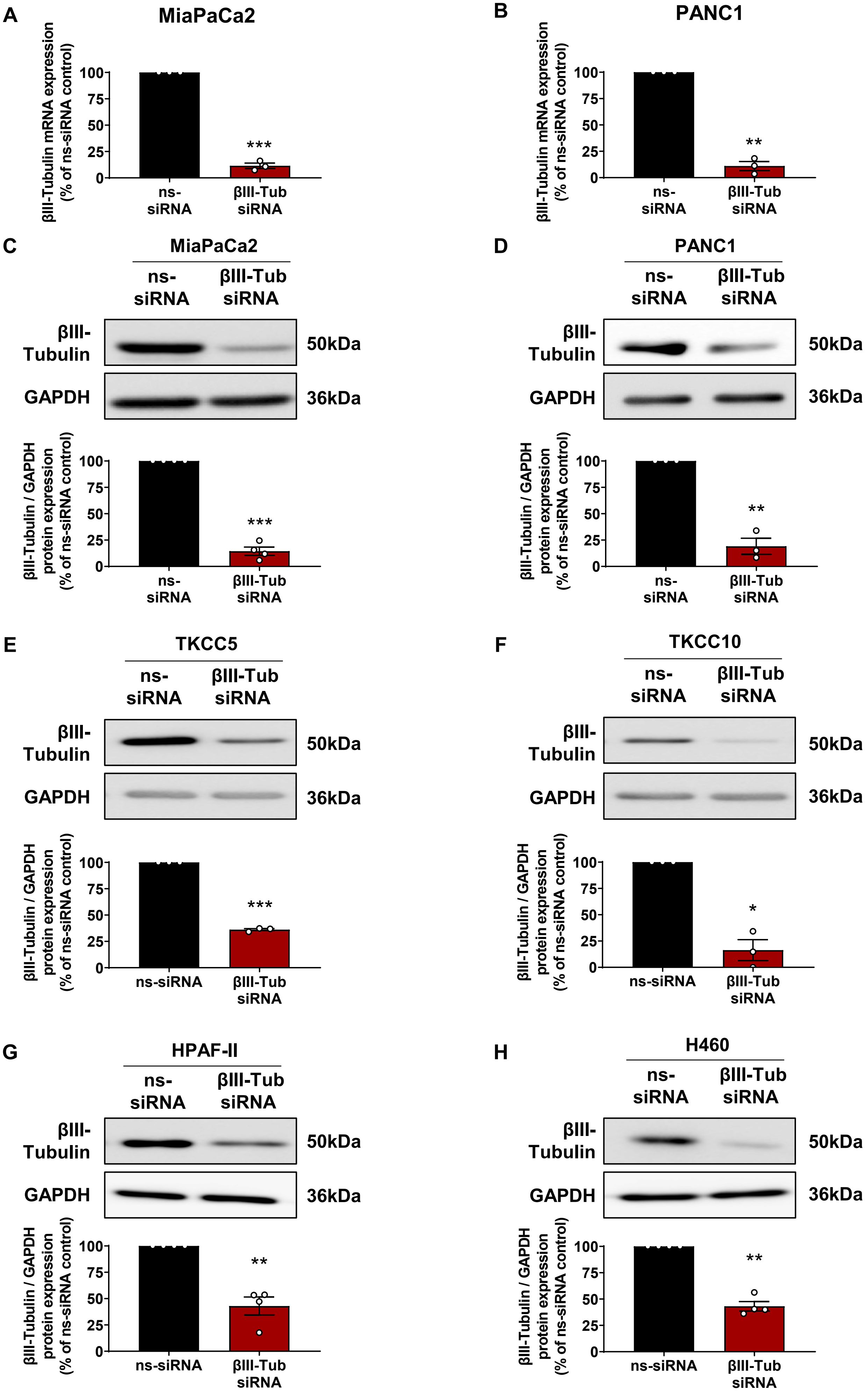


**Supplementary Figure 1: Confirmation of βIII-Tubulin knockdown in PDAC and NSCLC cells.**

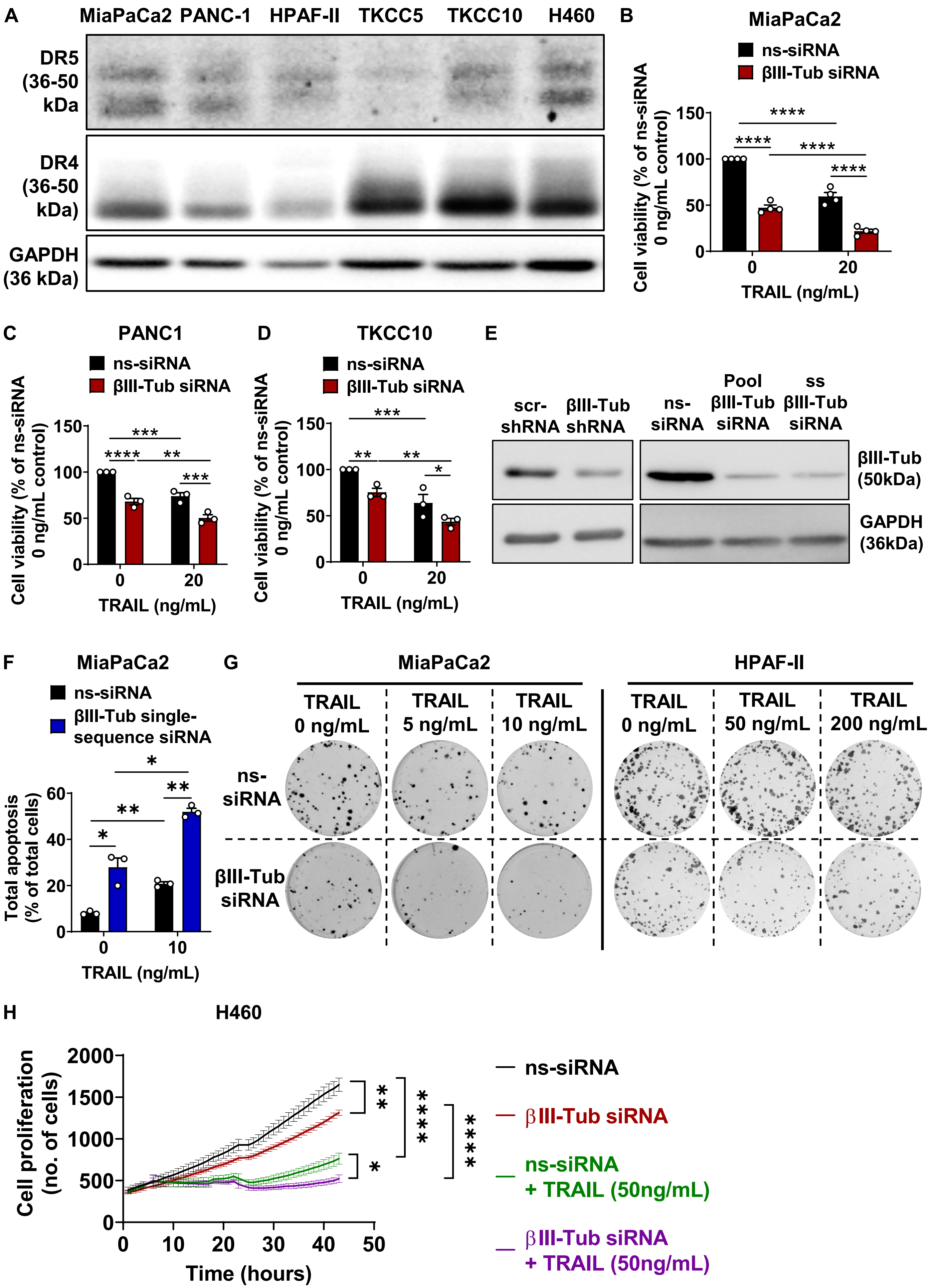


**Supplementary Figure 2: βIII-Tubulin silencing in PDAC and NSCLC cells increased sensitivity to TRAIL.**

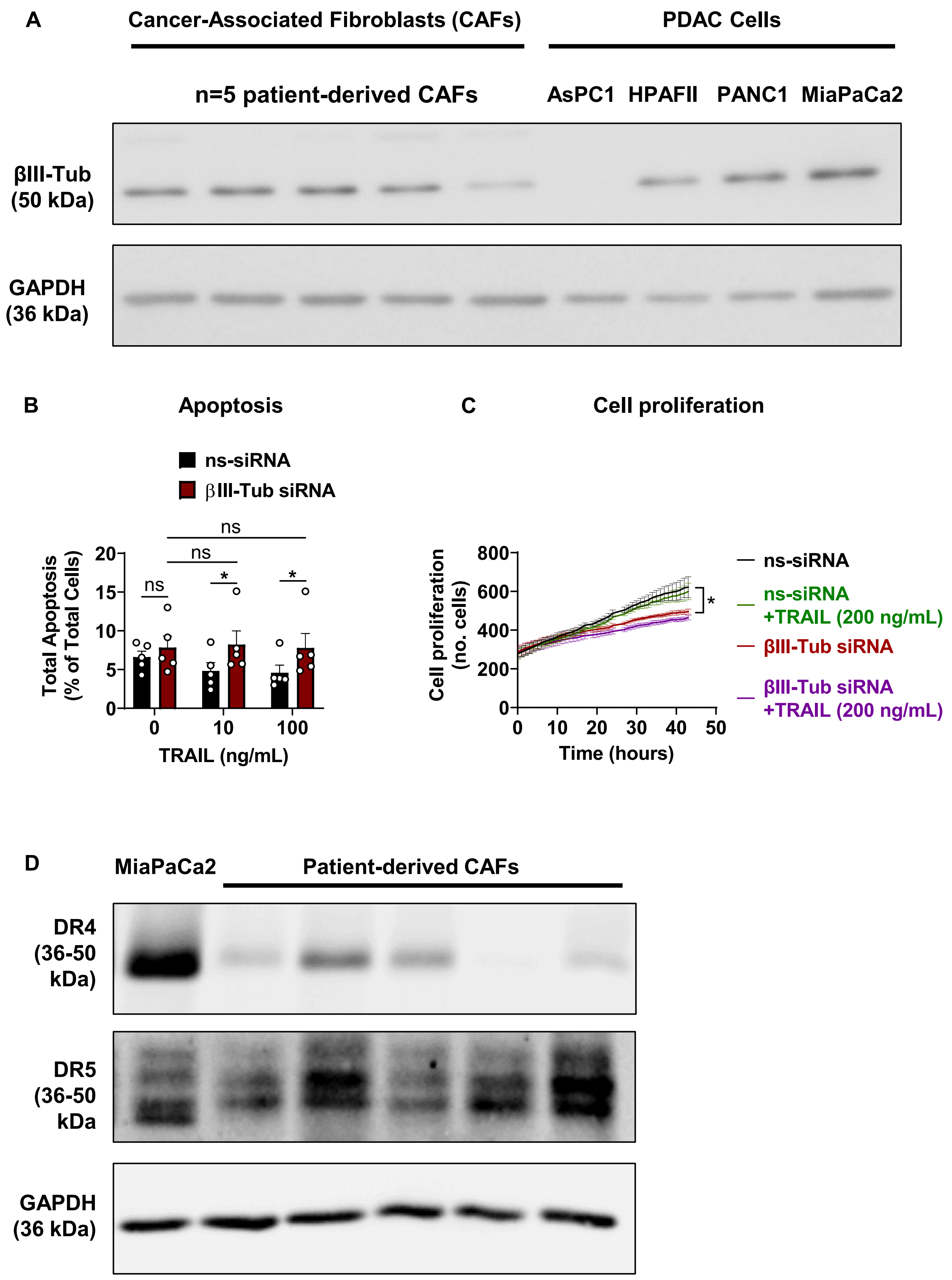


**Supplementary Figure 3: βIII-Tubulin silencing in patient-derived PDAC CAF cells had no effect on TRAIL sensitivity.**

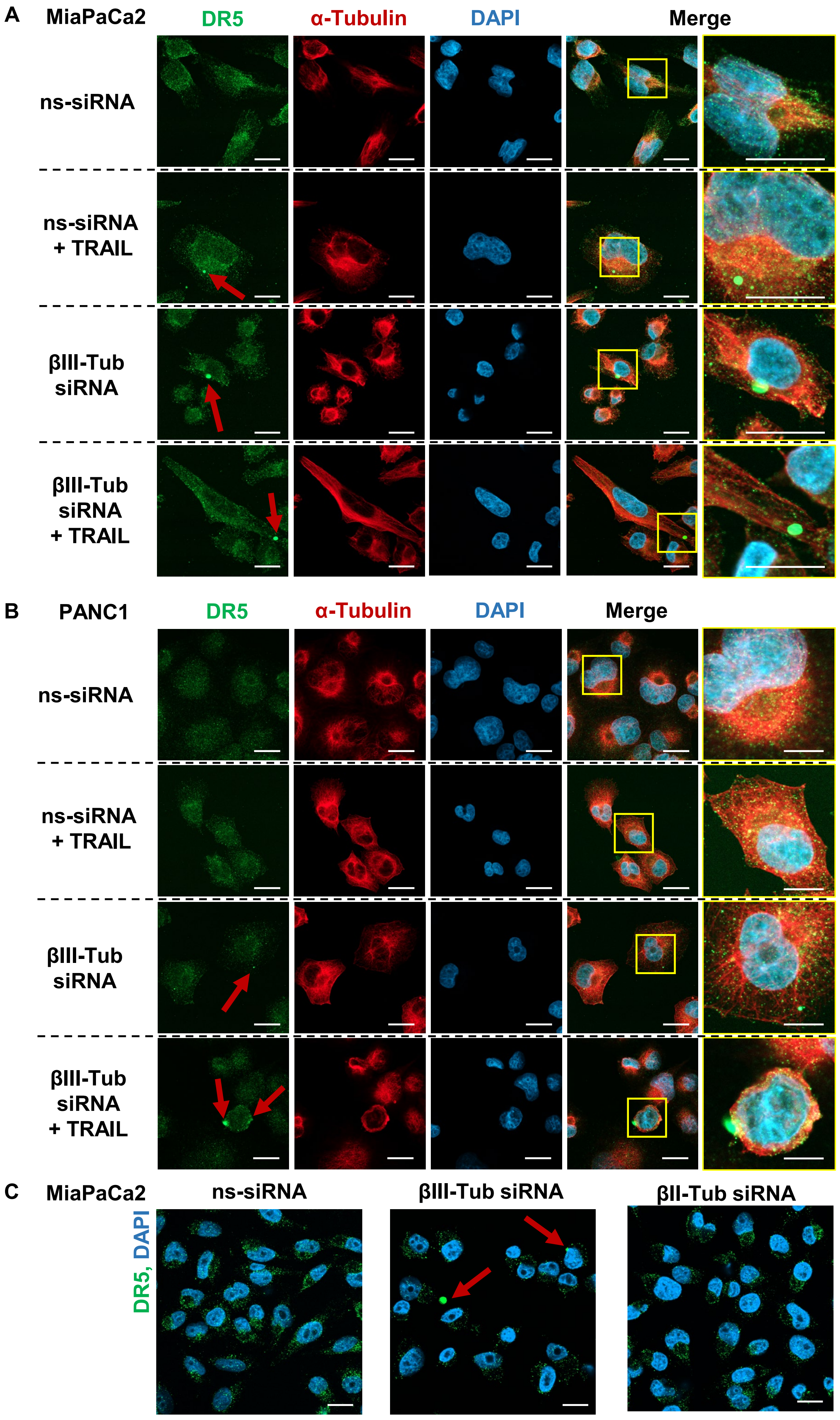


**Supplementary Figure 4: βIII-Tubulin silencing in PDAC cells triggered DR5 clustering.**

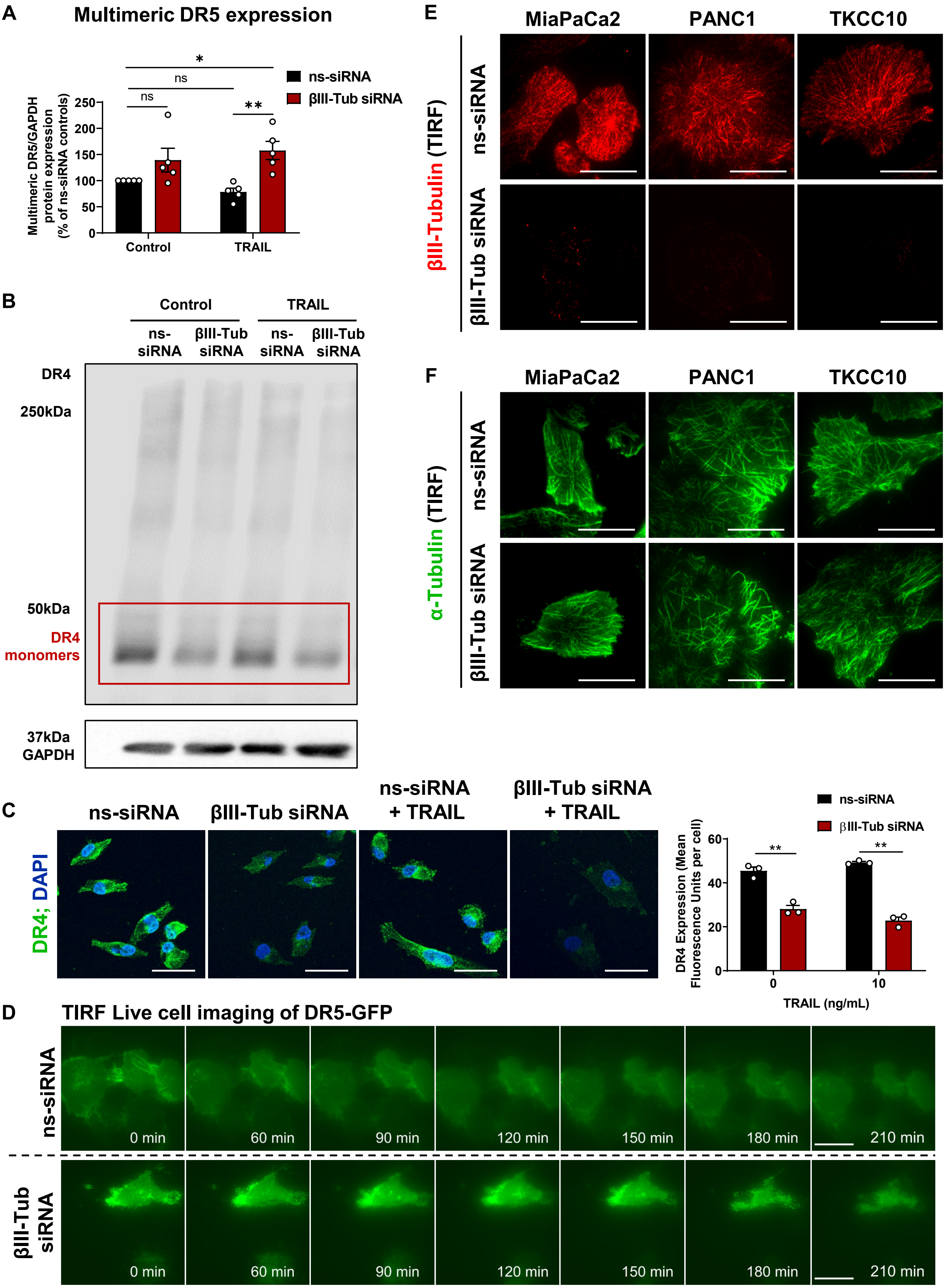


**Supplementary Figure 5. βIII-Tubulin regulates DR5 dynamics in PDAC cells.**

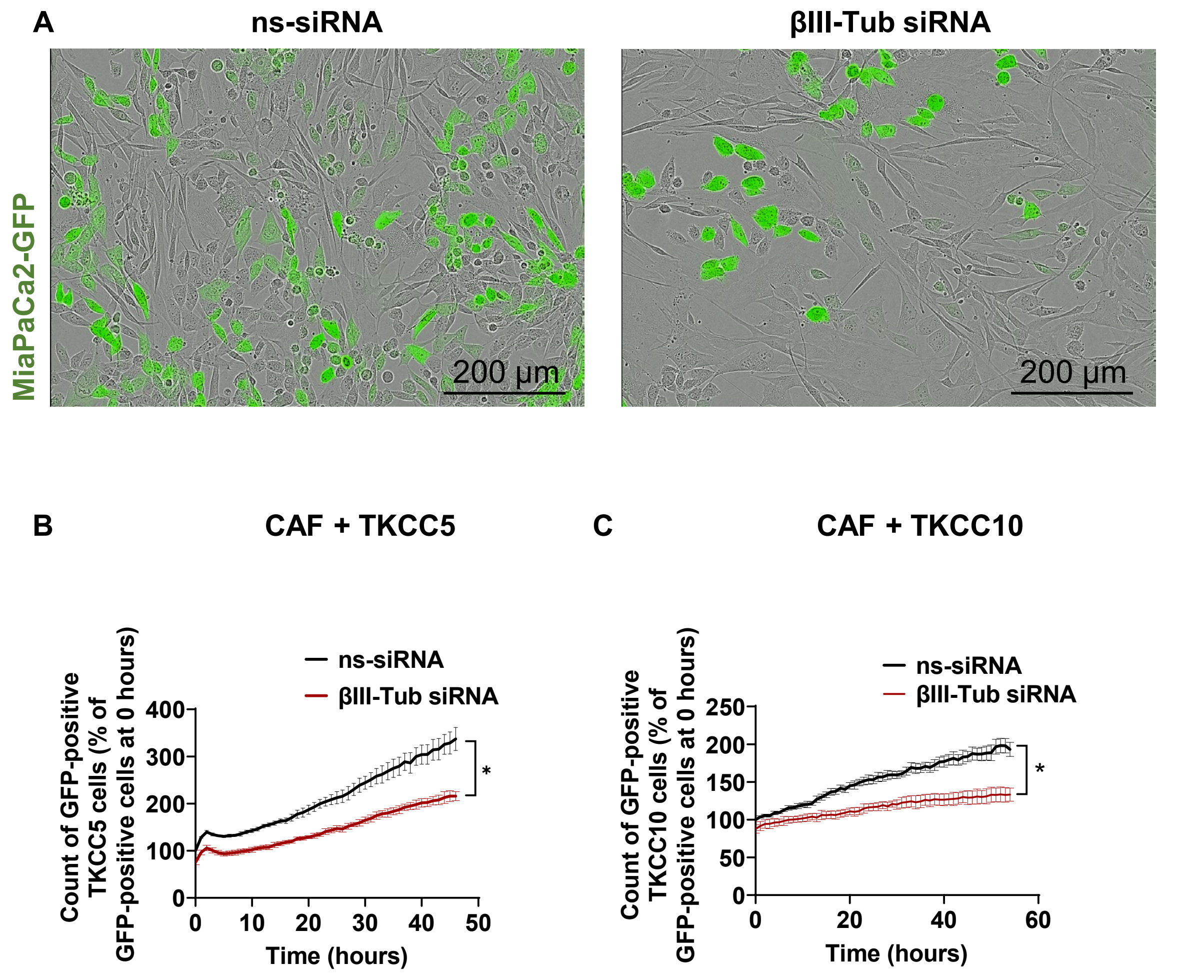


**Supplementary Figure 6: Anti-proliferative effects of βIII-tubulin silencing when pancreatic ductal adenocarcinoma (PDAC) cells are co-cultured with cancer associated fibroblasts (CAFs).**

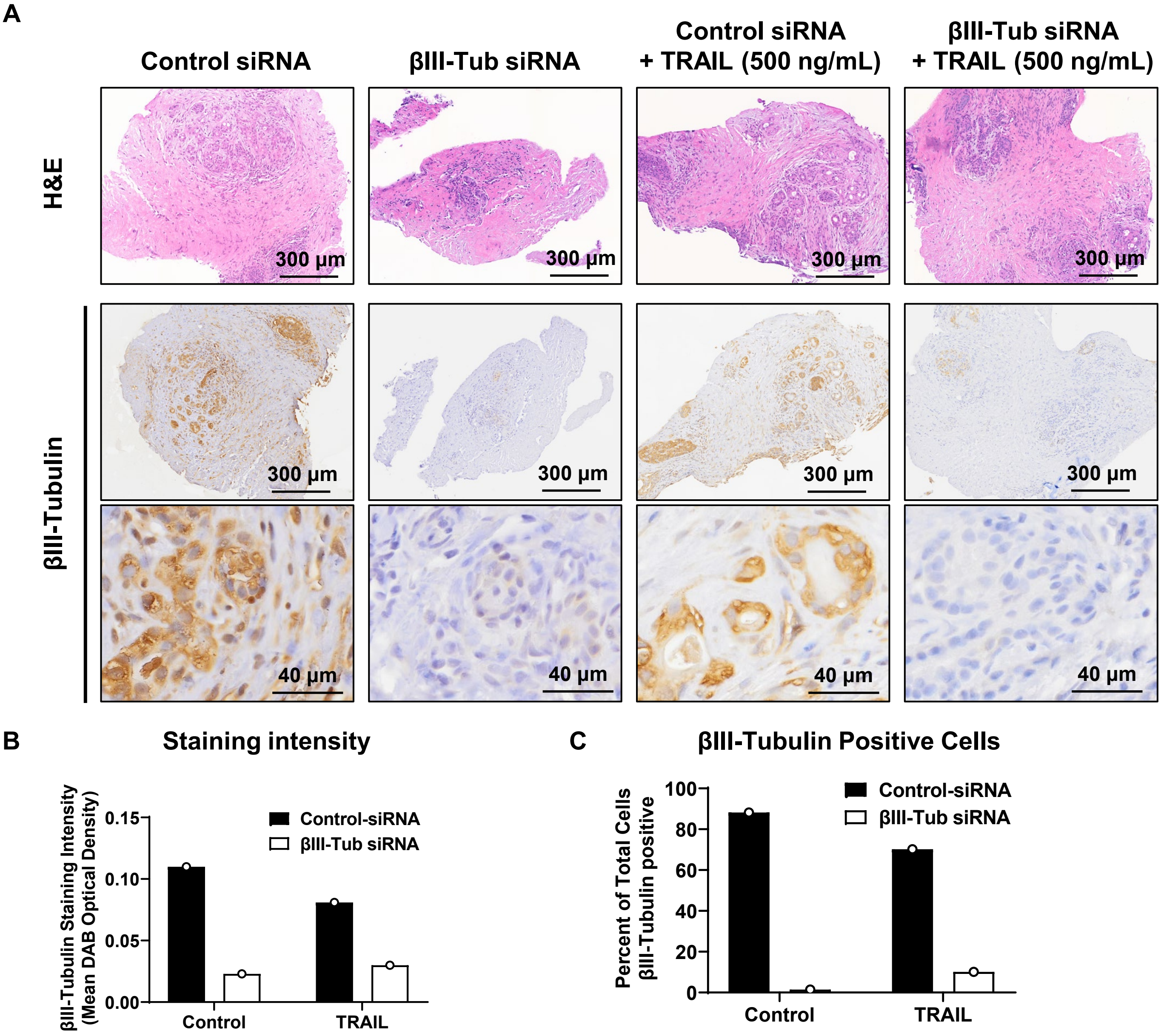


**Supplementary Figure 7: Star 3 nanoparticle delivery of βIII-tubulin siRNA led to a potent reduction in βIII-tubulin expression in patient-derived pancreatic ductal adenocarcinoma (PDAC) explants.**

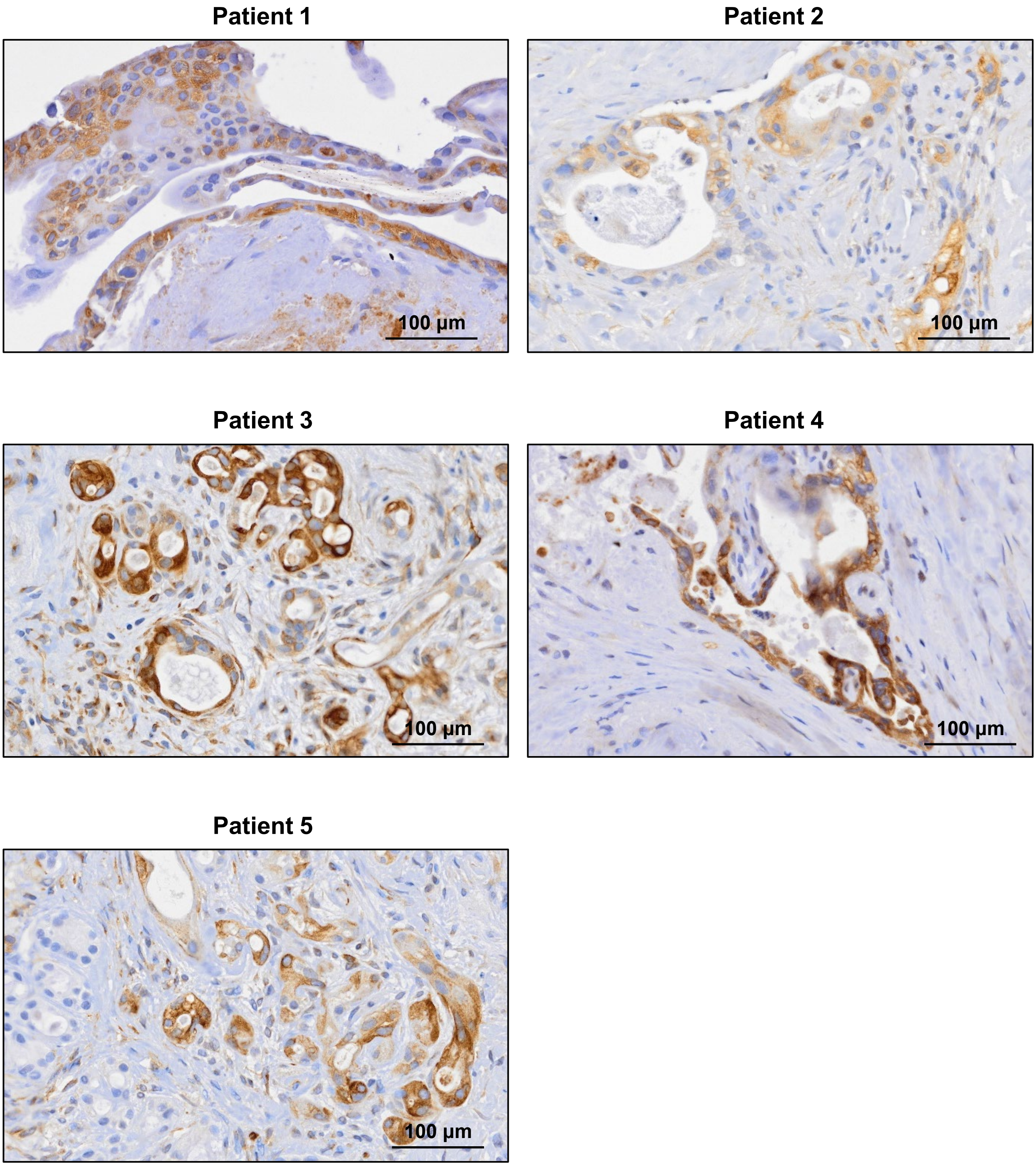


**Supplementary Figure 8: Patient derived tumour explants from 5 pancreatic ductal adenocarcinoma (PDAC) patients demonstrate high tumour expression of βIII-tubulin.**

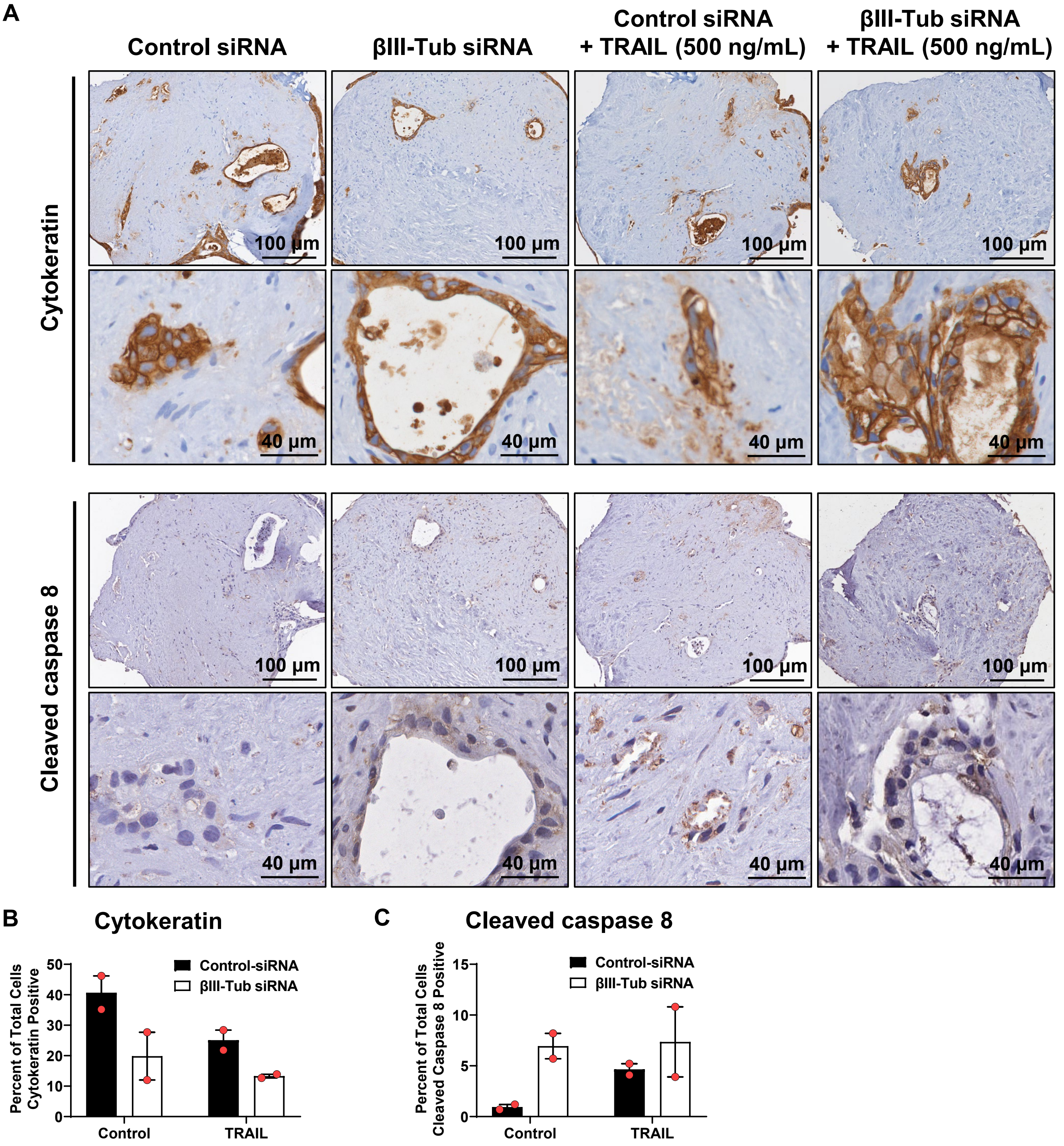


**Supplementary Figure 9: βIII-tubulin silencing combined with TRAIL in Patient 1 decreased tumour cell number and increased extrinsic apoptosis in pancreatic ductal adenocarcinoma (PDAC) patient-derived explants.**

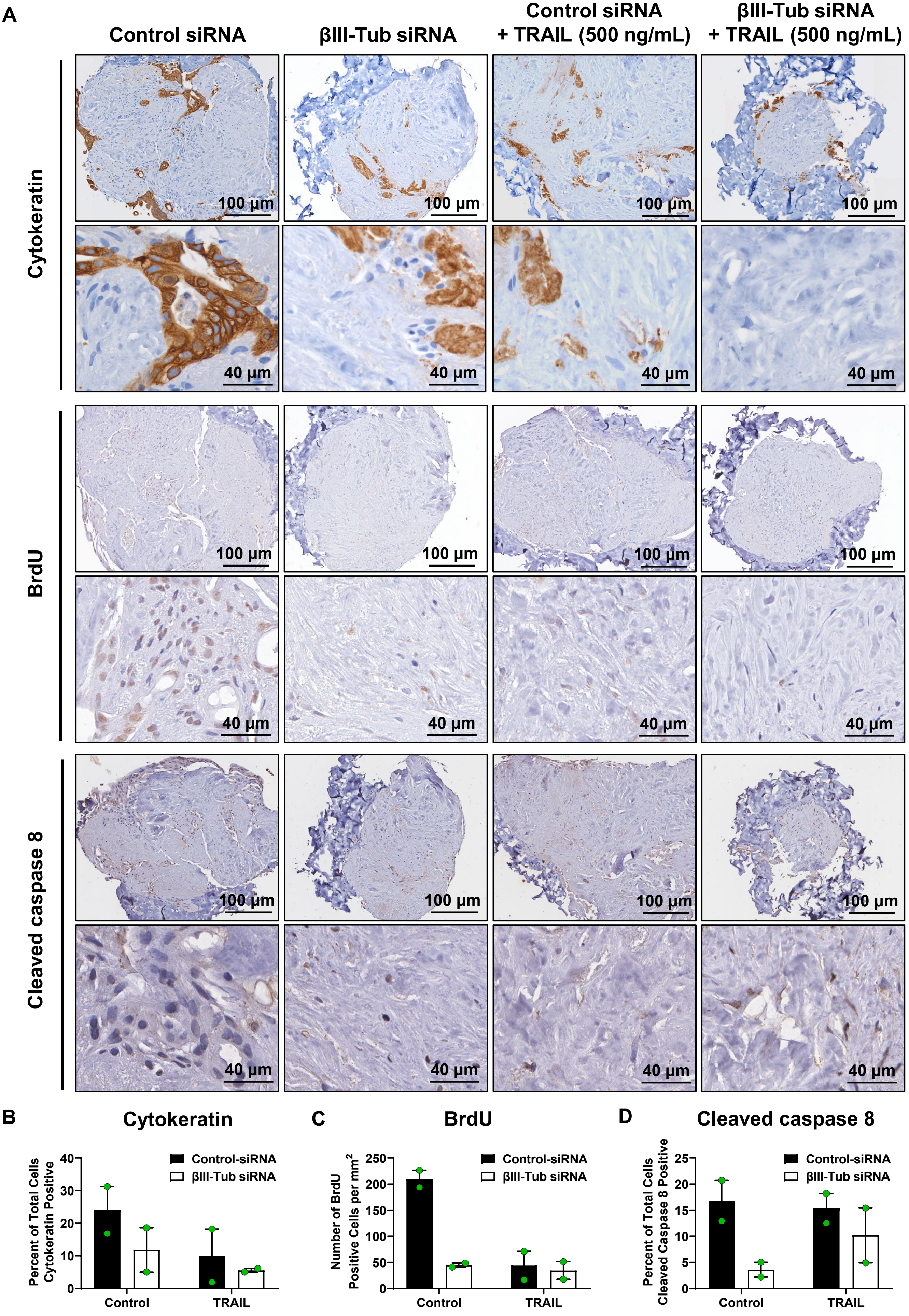


**Supplementary Figure 10: βIII-tubulin silencing combined with TRAIL in Patient 2 decreased tumour cell number and decreased cell proliferation in pancreatic ductal adenocarcinoma (PDAC) patient-derived explants.**

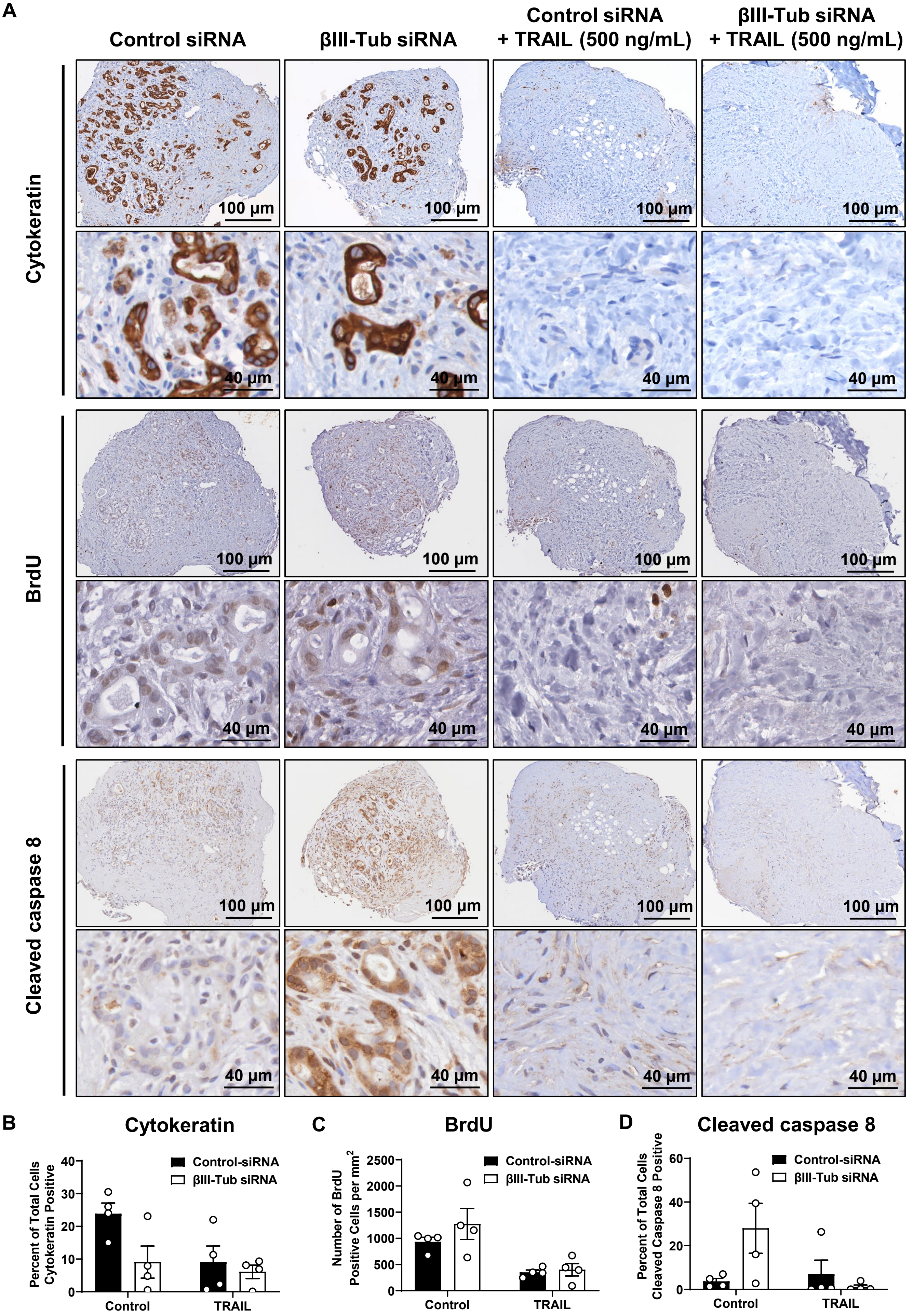


**Supplementary Figure 11: βIII-tubulin silencing combined with TRAIL in Patient 3 decreased tumour cell number and decreased cell proliferation in pancreatic ductal adenocarcinoma (PDAC) patient-derived explants.**

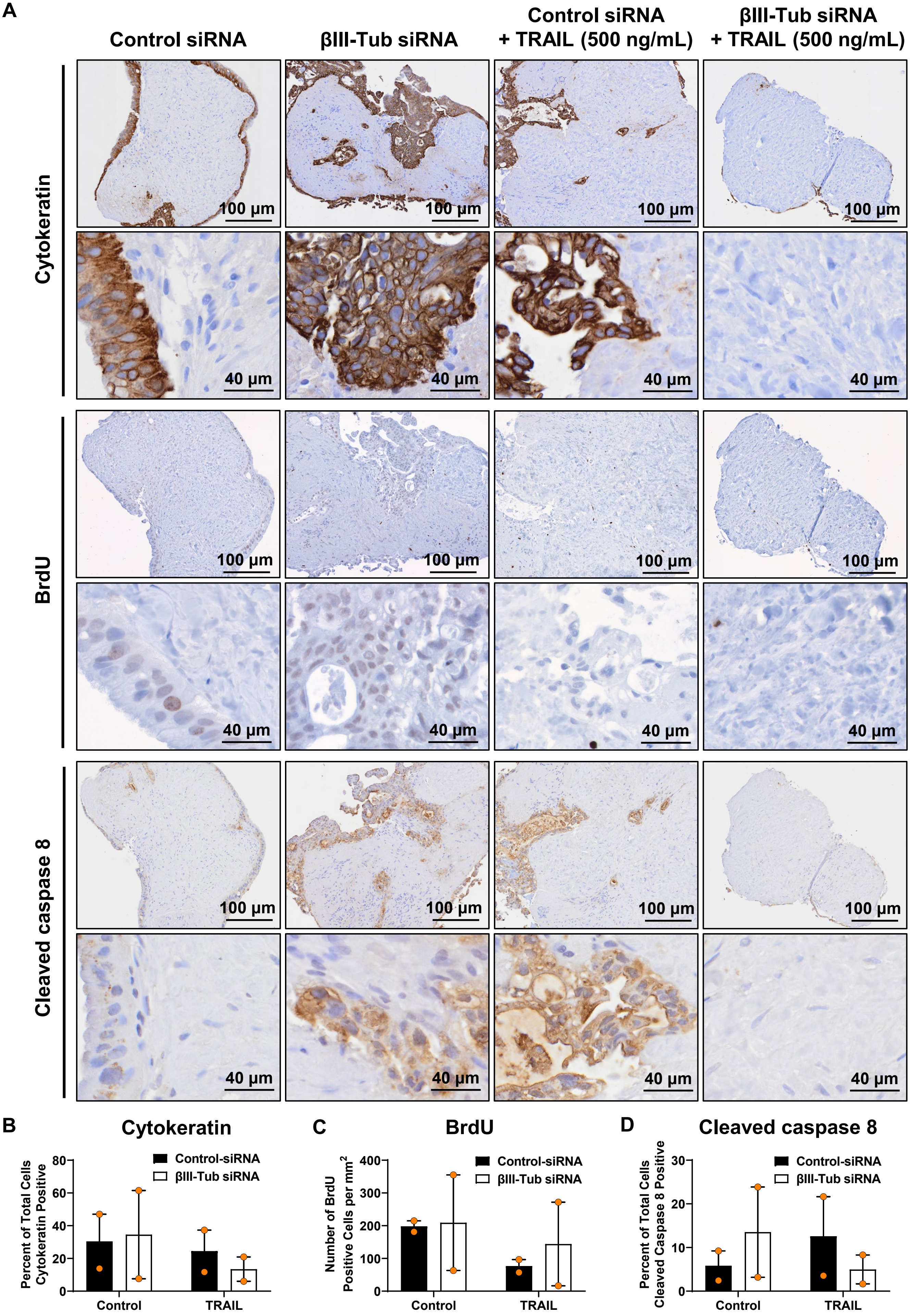


**Supplementary Figure 12: βIII-tubulin silencing combined with TRAIL in Patient 4 decreased tumour cell number in pancreatic ductal adenocarcinoma (PDAC) patient-derived explants.**

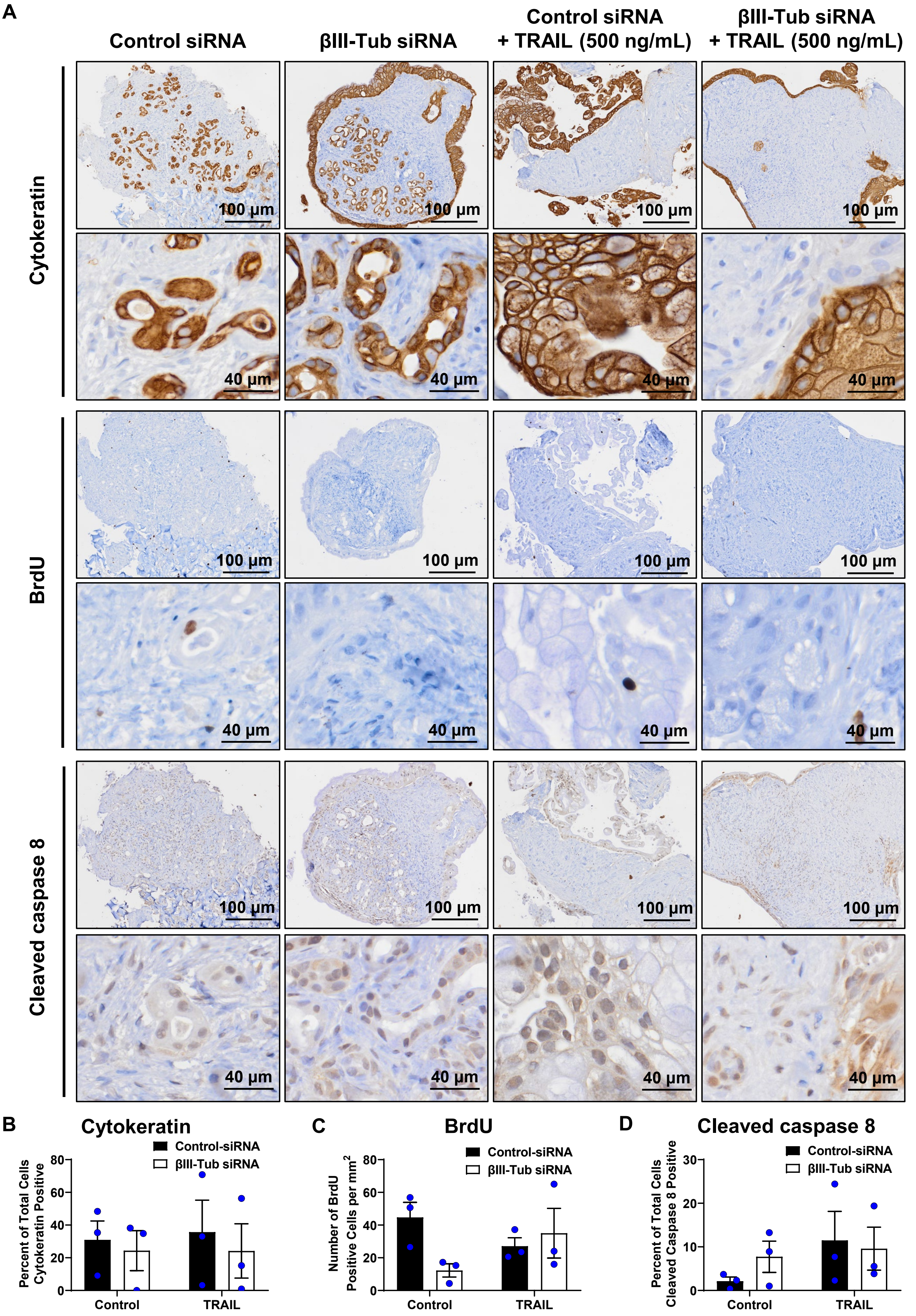


**Supplementary Figure 13: βIII-tubulin silencing combined with TRAIL in Patient 5 increased extrinsic apoptosis in pancreatic ductal adenocarcinoma (PDAC) patient-derived explants.**

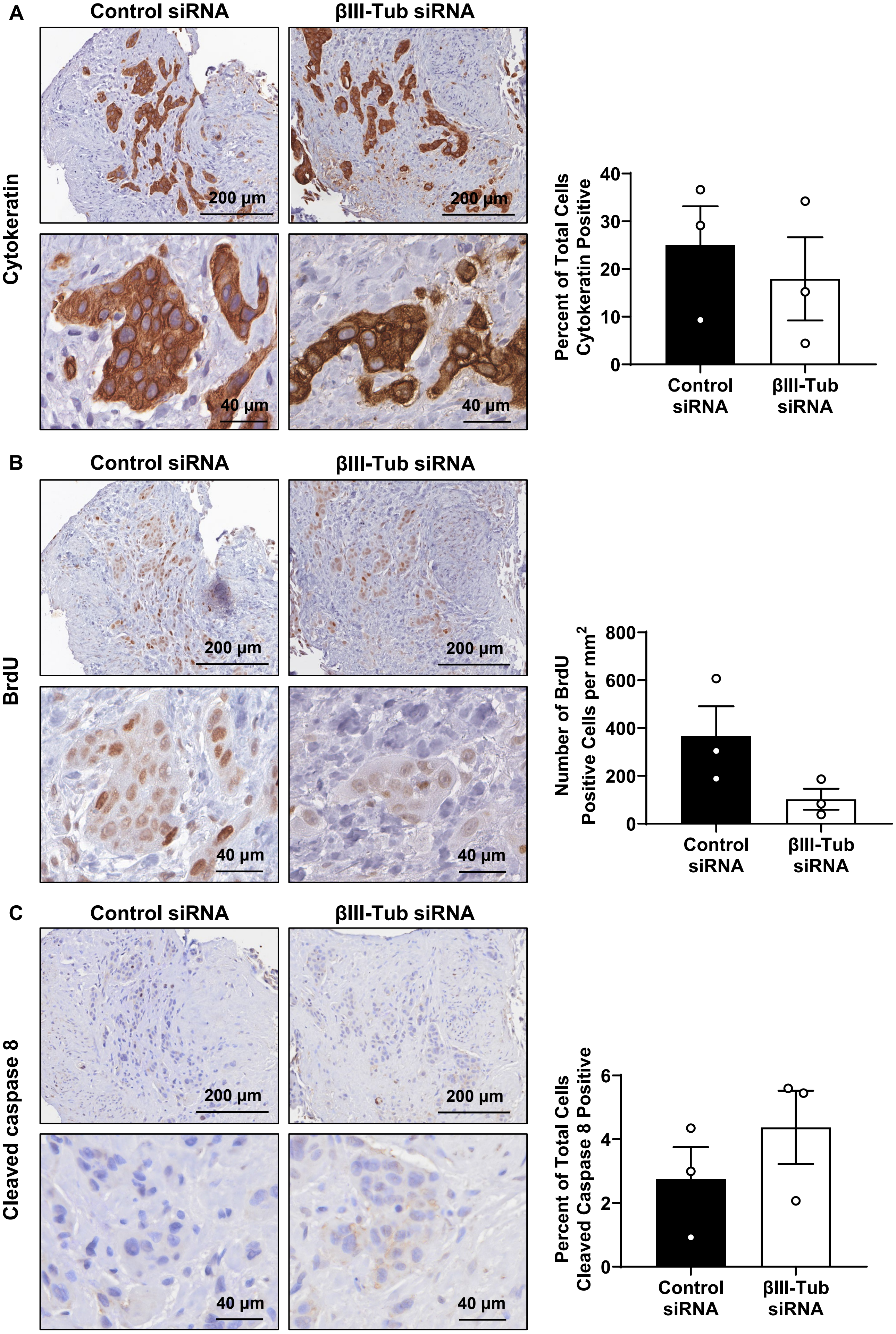


**Supplementary Figure 14: βIII-tubulin silencing in Patient 6 decreased cell proliferation and increased extrinsic apoptosis in tumour explants derived from a stomach metastasis of a patient with pancreatic ductal adenocarcinoma (PDAC).**

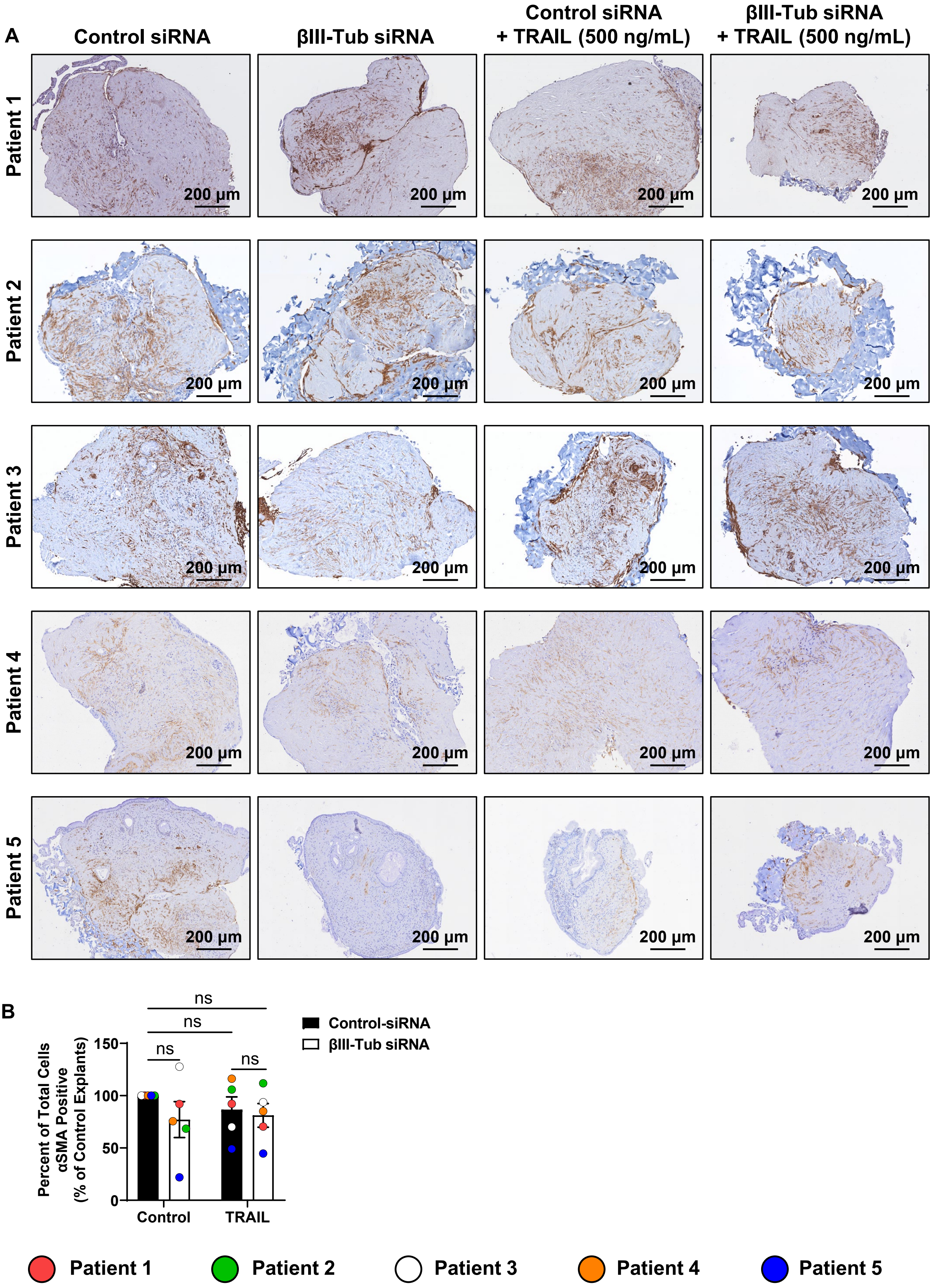


**Supplementary Figure 15: βIII-tubulin silencing and TRAIL treatment had patient-specific effects on cancer-associated fibroblast (CAF) cell number in pancreatic ductal adenocarcinoma (PDAC) tumour explants.**

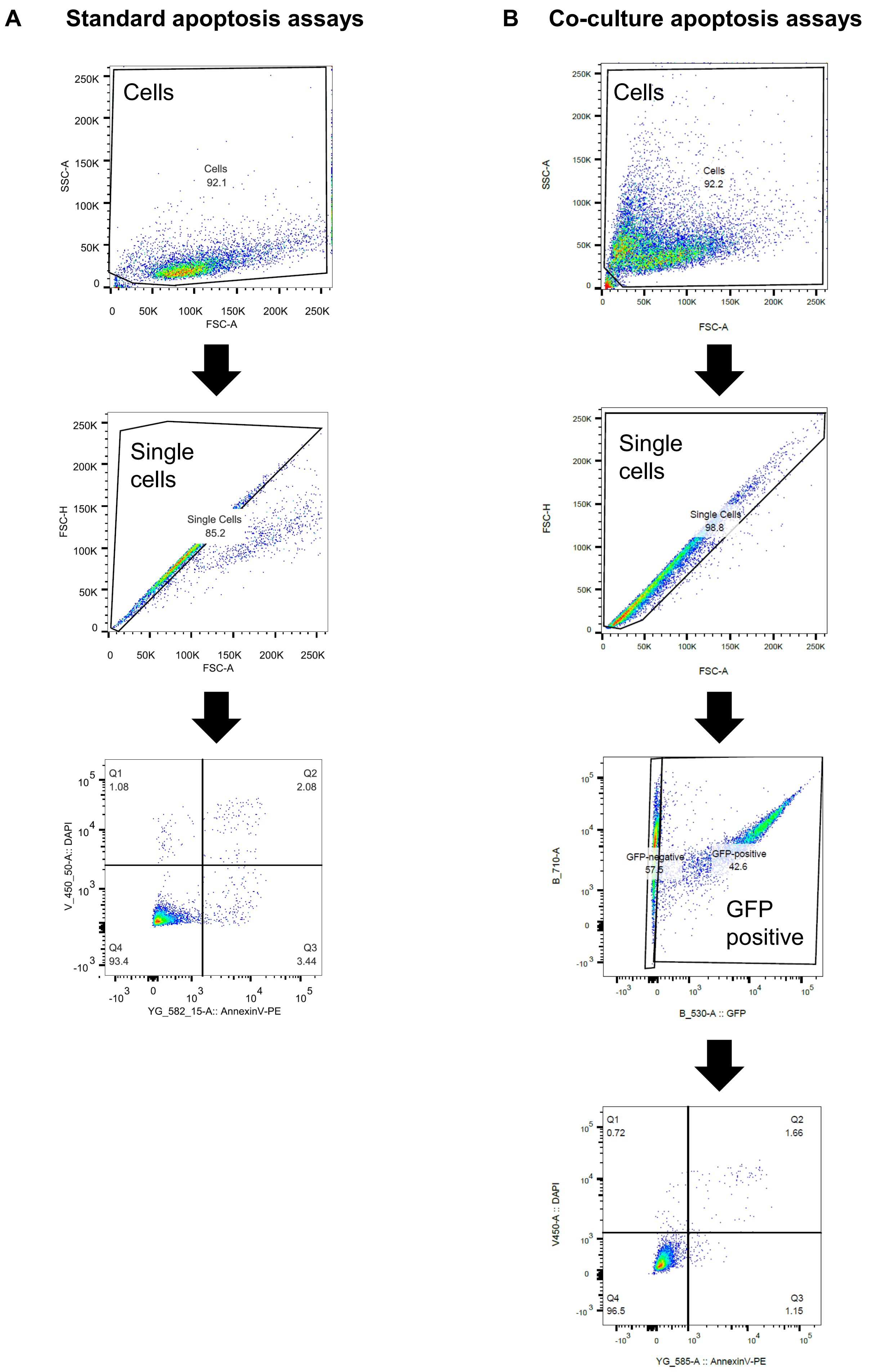


**Supplementary Figure 16: Gating strategy for apoptosis assays.**

**A** **Phase contrast**

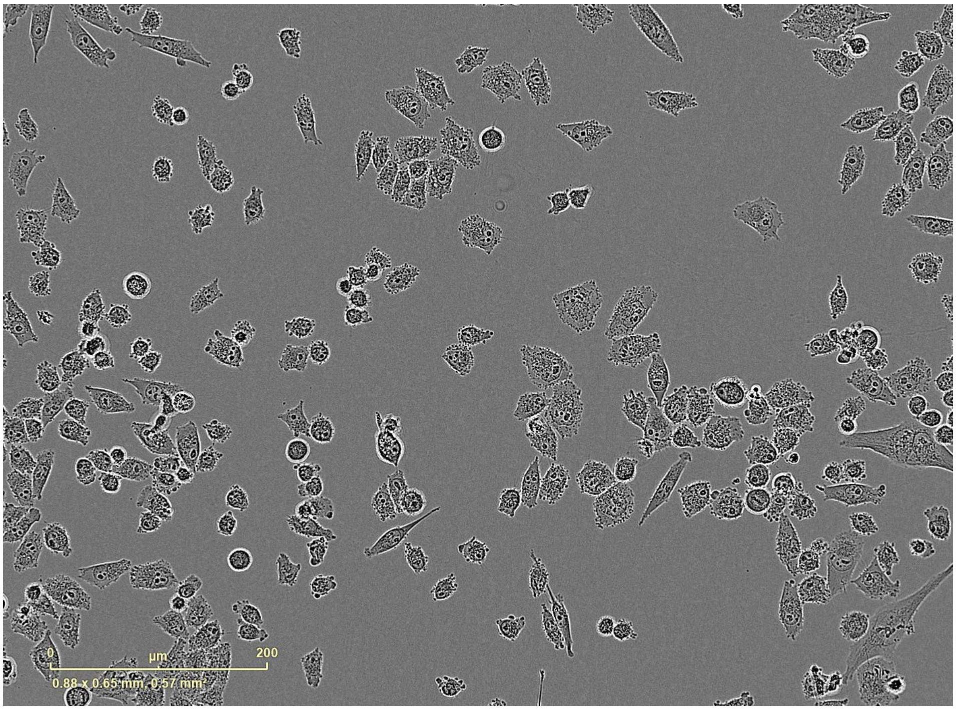


**B** **Confluence mask**

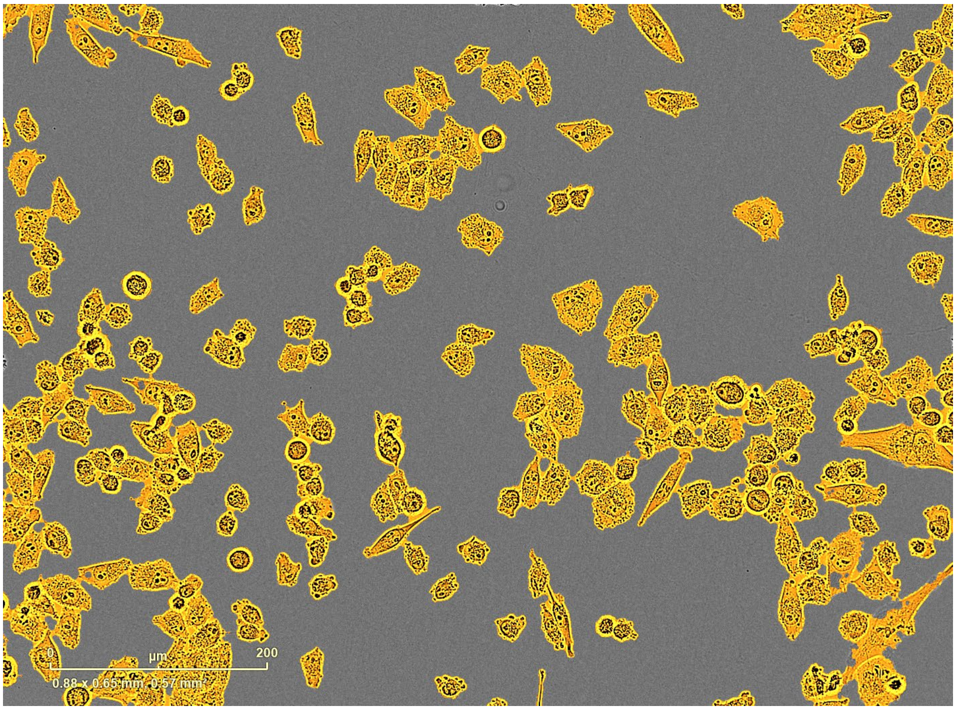


**Supplementary Figure 17: Analysis of cell proliferation on IncuCyte® S3 using confluence metrics.**

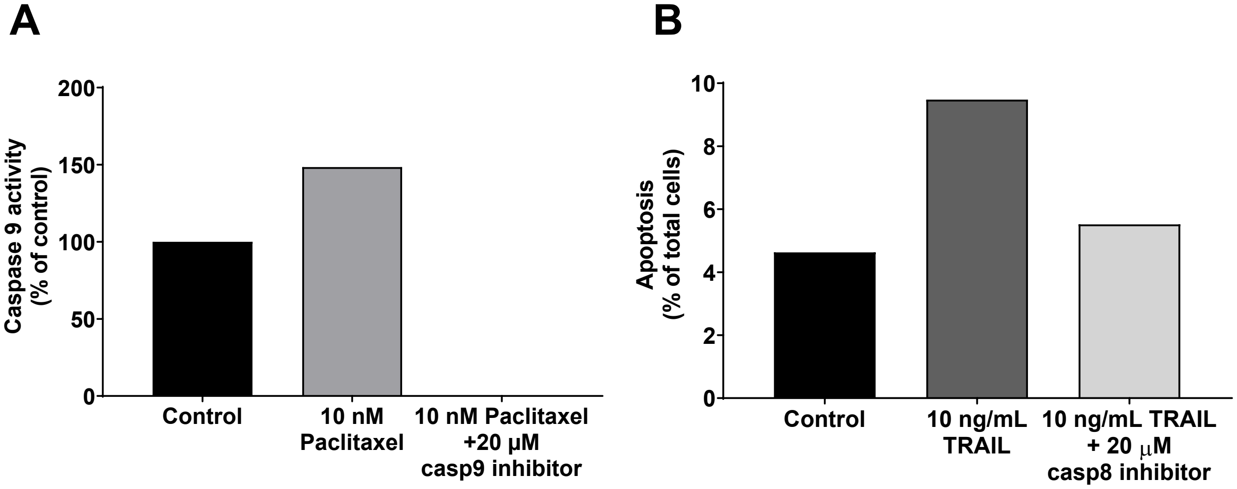


**Supplementary Figure 18: Validation of caspase inhibitors.**

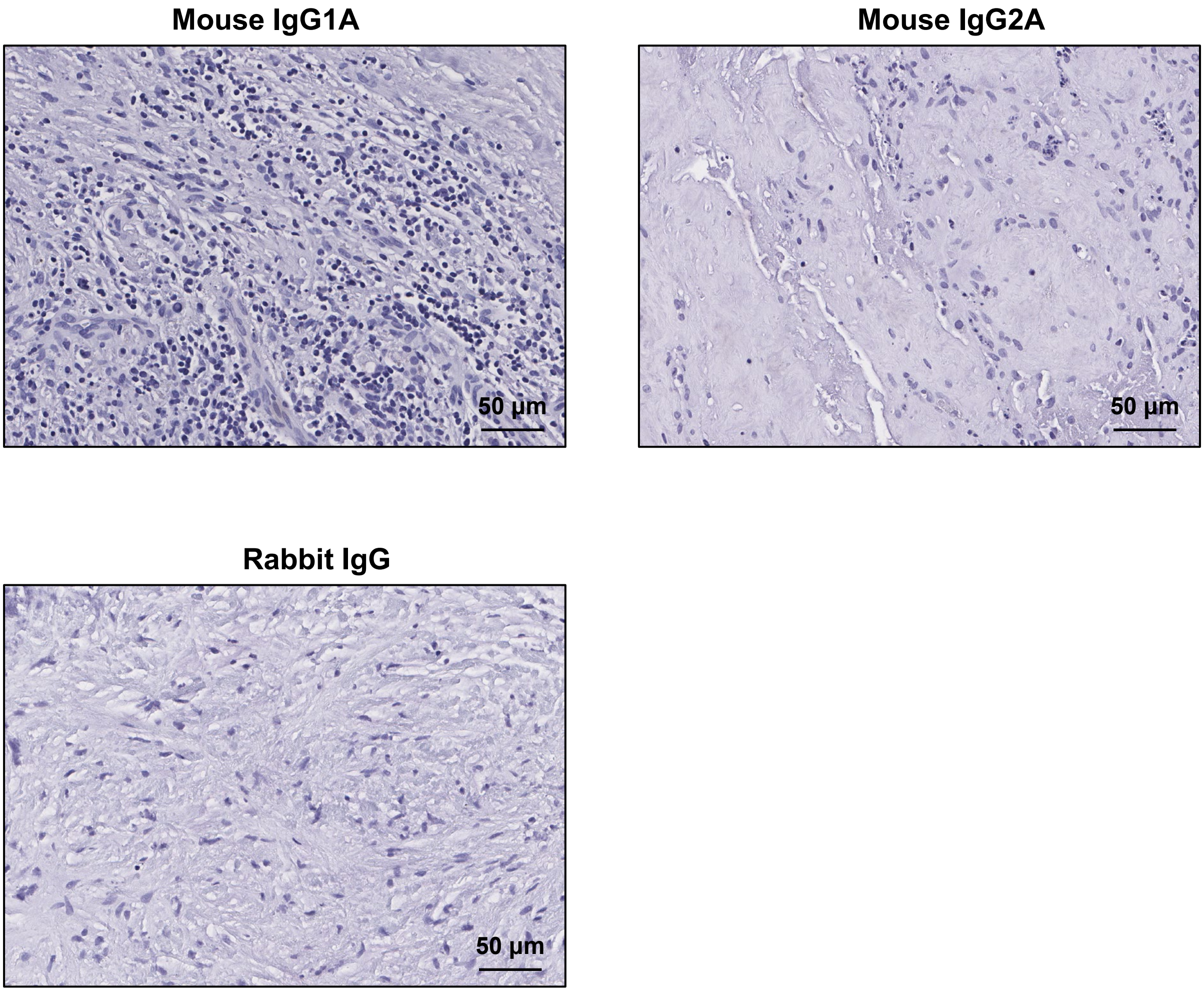


**Supplementary Figure 19. Immunohistochemistry isotype controls. Representative images of each isotype control antibody used for immunohistochemistry.**

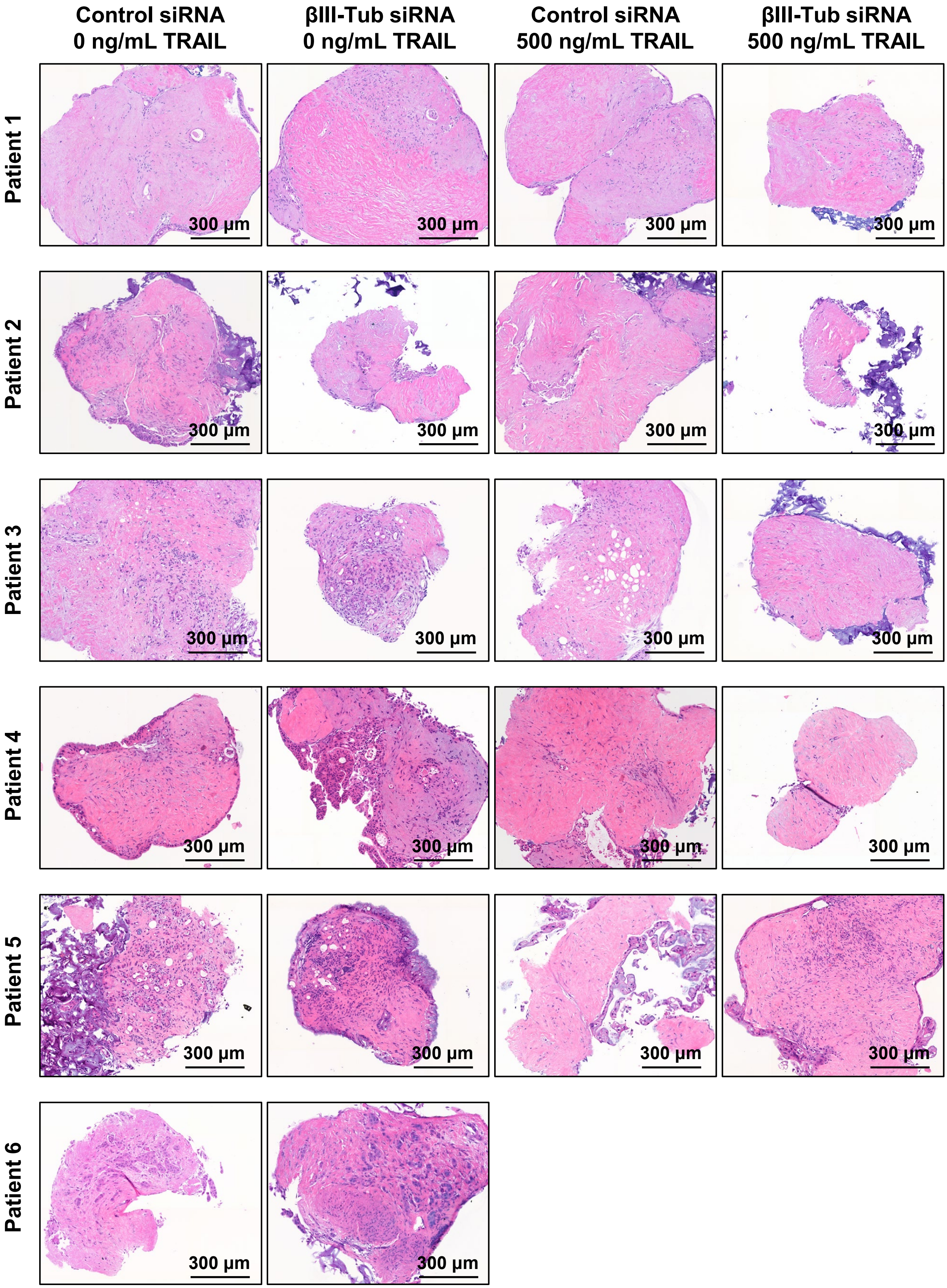


**Supplementary Figure 20. H&E staining of pancreatic ductal adenocarcinoma (PDAC) tumour explants from patients 1-6 used for βIII-tubulin experiments.**

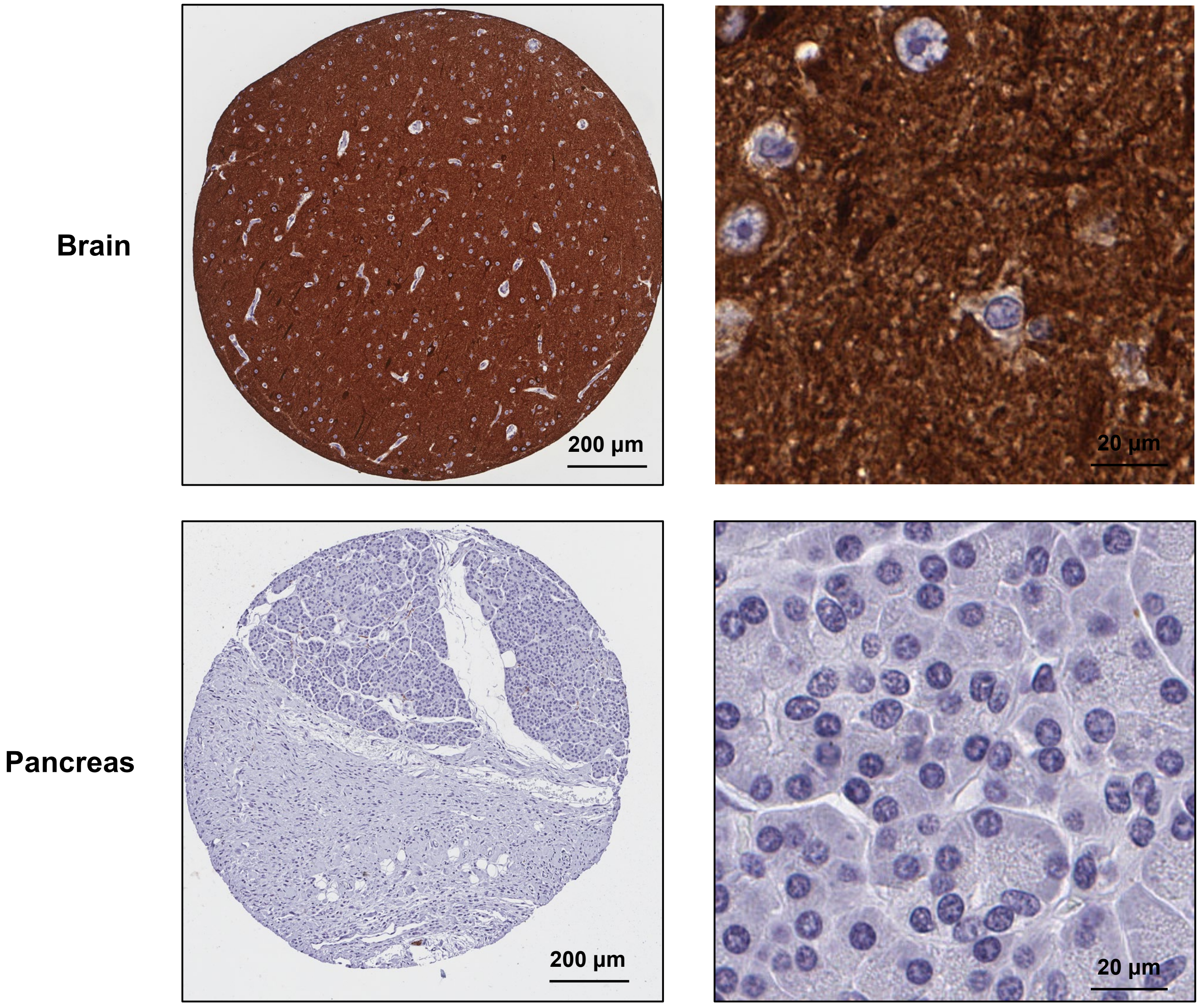


**Supplementary Figure 21. Validation of βIII-tubulin antibody.**
